# Modular Organization of Electrical Fluctuations in the Mouse Brain

**DOI:** 10.1101/2025.10.01.679820

**Authors:** Xingyun Wang, Jean-Claude Béϊque, Richard Naud

**Affiliations:** Department of Cellular and Molecular Medicine, University of Ottawa; Department of Physics, University of Ottawa; Center for Neural Dynamics and Artificial Intelligence, University of Ottawa; University of Ottawa Brain and Mind Research Institute University of Ottawa

## Abstract

A powerful computational feature of the brain is its ability to compartmentalize different functions in a way that can be flexibly recombined. Experimental evidence for such modularity arose from cytoarchitecture, connectivity, and electroencephalograms, while magnetic resonance imaging could attest that this modularity is dynamic. More spatially precise and more widely recordable, the local field potential (LFP) has an unknown brain wide organization. Here, we developed deep electrospectroscopy, a method to assess nonlinear spectral similarity between the LFP of different areas. A brain-wide application of this technique to the mouse brain revealed an organization composed of groups of spectrally similar areas. Such communities were mostly found in the fore-brain and showed an organization that did not strictly follow the cytoarchitecture. For instance, visual parts of the cortex and the visual parts of the colliculus were found in the same community. These electrospectral communities reshaped with context, showing splitting and merging operations, growing in size around brain areas required for a task. In particular, upper and lower limb parts of the somatosensory cortex were primarily in separate communities but merged during a task that required turning a wheel. Similarly, oculomotor reflexes and associative parts of the thalamus merged during the visuo-motor task. These analyzes show that LFPs are organized in a modular fashion, offering a window onto subcortical and layer-specific contributions to the compositionality of cortical brain functions.

## Introduction

Recent brain-wide recordings of activity have made clear that every part of the brain encodes almost every aspect of the world to some degree [1, 2]. These observations come as a puzzle if one consider the brain as a set of separated modules feeding into a centralized control of behavior [3]. Cognitive science and artificial intelligence investigations have proposed modules work in parallel [4], that motor control forms hierarchies [**?**], and that modules can reconfigure into compound functions [5, 6]. How does such a modular reconfiguration take place in the brain is therefore a central question in neuroscience. Brain modularity is observable in patterns of similarities and dissimilarities across different brain areas [7, 8]. Focusing on activity-related signals, coarse modules were found in brain-wide electroencephalograms (EEGs) and in functional magnetic resonance imaging (fMRI) [9, 10]. These modules were shown to be highly similar to the modules found in patterns of traced connectivity data [11], suggesting that the modular structure of brain connectivity is echoed by its connectivity. Studies in mice could further probe the question of what underlies this modularity. A modular organization was found in the cytoarchitecture, connectivity and transcriptome [12, 13, 14]. Yet, the recordings focused on cortex and often lacked both dynamics and layer precision. A picture of how modules reconfigure the different layers of cortical and subcortical elements is lacking.

With recent developments in silicon probes [15, 1, 16], local field potentials (LFPs) can, in principle, be recorded with high-density, and deep structures can be reached. Offering a microscopic picture of the electrical fluctuations across almost the entire mouse brain [17, 18], this signal is thought to reflect a mixture of activity patterns from afferents and the local population [19]. Synchronized afferent patterns, traveling waves, and inter-area synchronization [20, 21, 22, 23] are specific activity patterns that shape the LFP and can endow it with a modular structure. Yet, while many location-specific LFP patterns have been found [24, 25, 26, 27, 28, 29, 30, 31, 32, 33, 34, 35], little is known about how the LFP is organized across the entire brain, let alone whether its organization is modular.

Here, we reasoned that patterns of similarity could be found in the spectrograms of the LFP recorded across almost the entire mouse brain, but that these patterns may require analysis methods more perceptive than the correlation coefficient. This article is constructed in two sections. First, we trained deep artificial networks to localize LFPs, integrating possibly nonlinear relationships between features of the LFP and brain areas. Second, we use this model to assess the similarity and dissimilarity of areas, and thus to reveal the presence, structure and task-dependence of modules.

## Results

We used publicly available, multi-lab, brain wide Neuropixels recordings (International Brain Lab; IBL [17, 18, 36, 2]). In this standardized dataset, mice were cued with an auditory tone and a visual grating to steer a wheel. For each trial and for each channel, we extracted the spectrogram from one second of LFP recordings taken during the intertrial time period and at the onset of the task (Fig. 1a). The use of LFP spectrograms, or electrospectra, preserved some information about interactions between timescales, about transient synchronization to task features and about nearby multi-unit activity [37]. After quality control and shaping the data for our purposes (see Methods), we obtained LFPs from 472 different brain regions across 127 animals and 12 labs, for a total of 1 687 471 independent LFP segments (samples) taken during the intertrial period. While we present our analysis of this inter-trial period first, we will later present an analysis the other 1.7 M samples taken during the task period.

**Figure 1:**
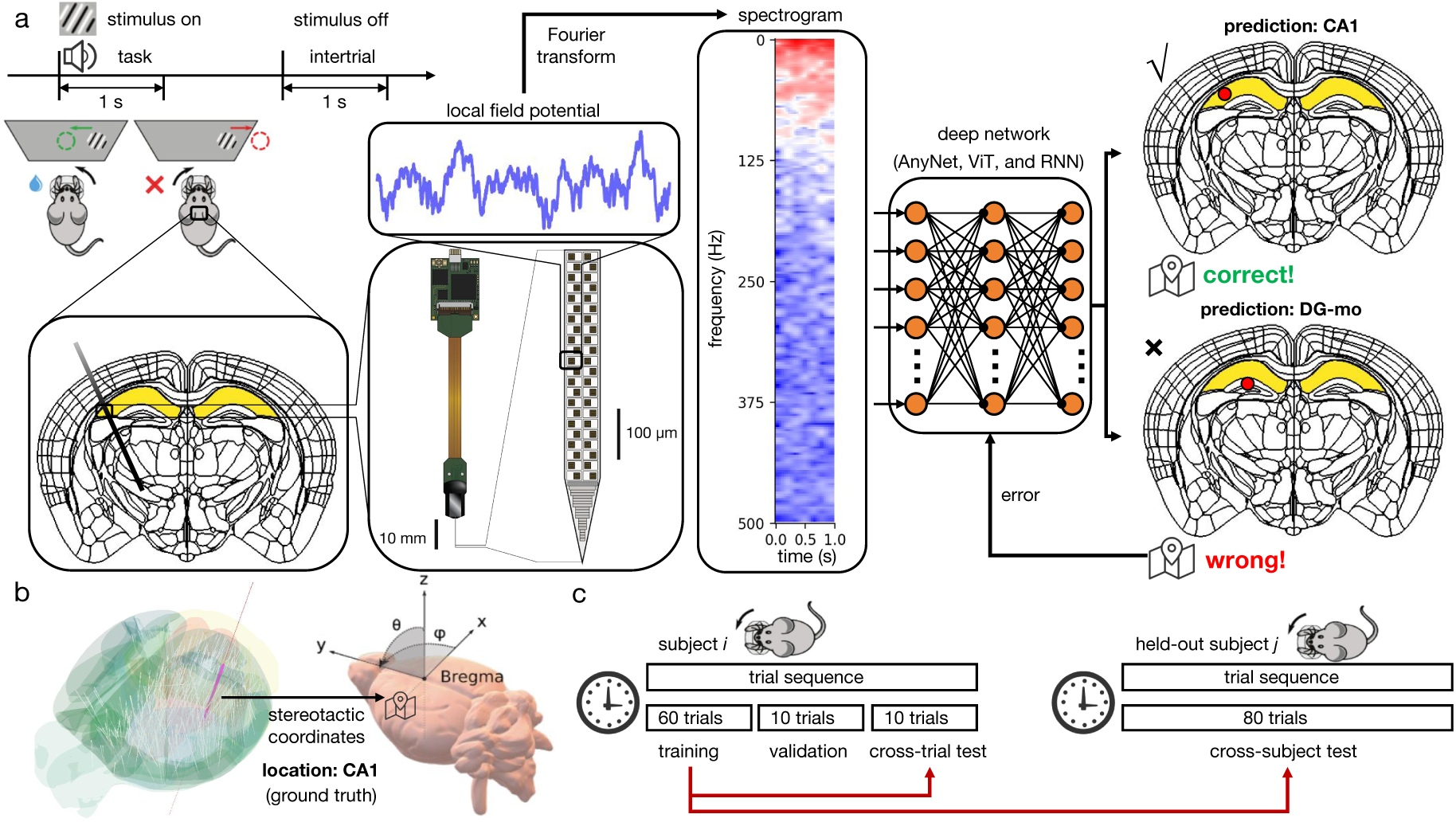
Deep learning LFP localization. **a.** Workflow schematic for brain area classification. Opensource Neuropixels data from the IBL was used for both task and intertrial states of a period of 1 second. We used LFP recorded during the inter-trial period. Each LFP on each electrode in each session was converted to a spectrogram and fed to an ANN (ViT, AnyNet or RNN) for localization. The ANN parameters were learned based on correct and erroneous prediction of the brain area of the recording, as illustrated here with a recording localized in cornu ammonis 1 (area ‘CA1’) but that could have been predicted to lie in the dentate gyrus (‘DG-mo’). **b.** The ground truth of a recording’s location were obtained by mapping its stereotaxis coordinates to a histologically-defined brain atlas (Allen Brain or Cosmos). **c.** Schematic of training and different test sets. For the first experiment of each subject except held-out subjects, we used first 60 trials to train and the following 10 trials for validation. The last 10 trials from each subject were were used to quantify the cross-trial test. The first 80 trials of held-out subjects consisted of the cross-subject test. Cross-trial and cross-subject tests correspond to in-distribution and out-distribution tests in machine learning.

### Nonlinear features of the LFP enable milimetric localization

As a preliminary investigation, we inquired about the spatial scale of changes in the electrospectra. A previous study has estimated the autocorrelation length (equivalent to the positional error of a linear model) of LFPs to a tenth or a third of the rostrocaudal length [38], depending on brain state. We therefore asked if using a method that can extract electrode positions from a nonlinear combinations of electrospectral features could do so with more precision than the average autocorrelation length. Accordingly, we trained deep networks (see Methods) in this task using samples from the inter-trial period and electrode position estimated using stereotaxis (Supplementary Fig. 2). We found positional error between 5 and 9 % of brain length, depending on axis (standard deviation of regression residuals 0.7-1.2 mm, compared to brain length of 14 mm). While preliminary, this observation supports the idea that our deep networks reveal a more precise structure than linear methods.

Since we are interested in the relationship between brain areas rather than raw positions, we then trained our networks to predict the source brain area for every sample [39] (Fig. 1a). To ensure reproducibility, we considered three different classifiers: a deep network developed for sound classification (AnyNet; [40]), a visual transformer (ViT; [41, 42]) and a recurrent neural network (RNN; [43]; Suppl. Fig. 1). Data was separated into a training set, a validation set and two test sets. One test set was made of different trials from the same animals used for training (cross-trial test), and the other test set was composed of different animals (cross-subject test; Fig. 1c), allowing us to test different types of generalization. We computed the average cross-trial accuracy for the three neural networks, along with a linear classifier. While a linear classifier was able to localize electrodes far better than chance (accuracy of 0.120 vs 0.002 for chance), its accuracy was surpassed by each of the three deep learning networks (Fig. 2a). Among these, performances were of the same order, with a small advantage for the ViT (0.389) and a small disadvantage for the RNN (0.312) and AnyNet midway (0.368). Therefore, deep learning captures combinations of features in the electrospectra that were not captured using linear methods.

**Figure 2:**
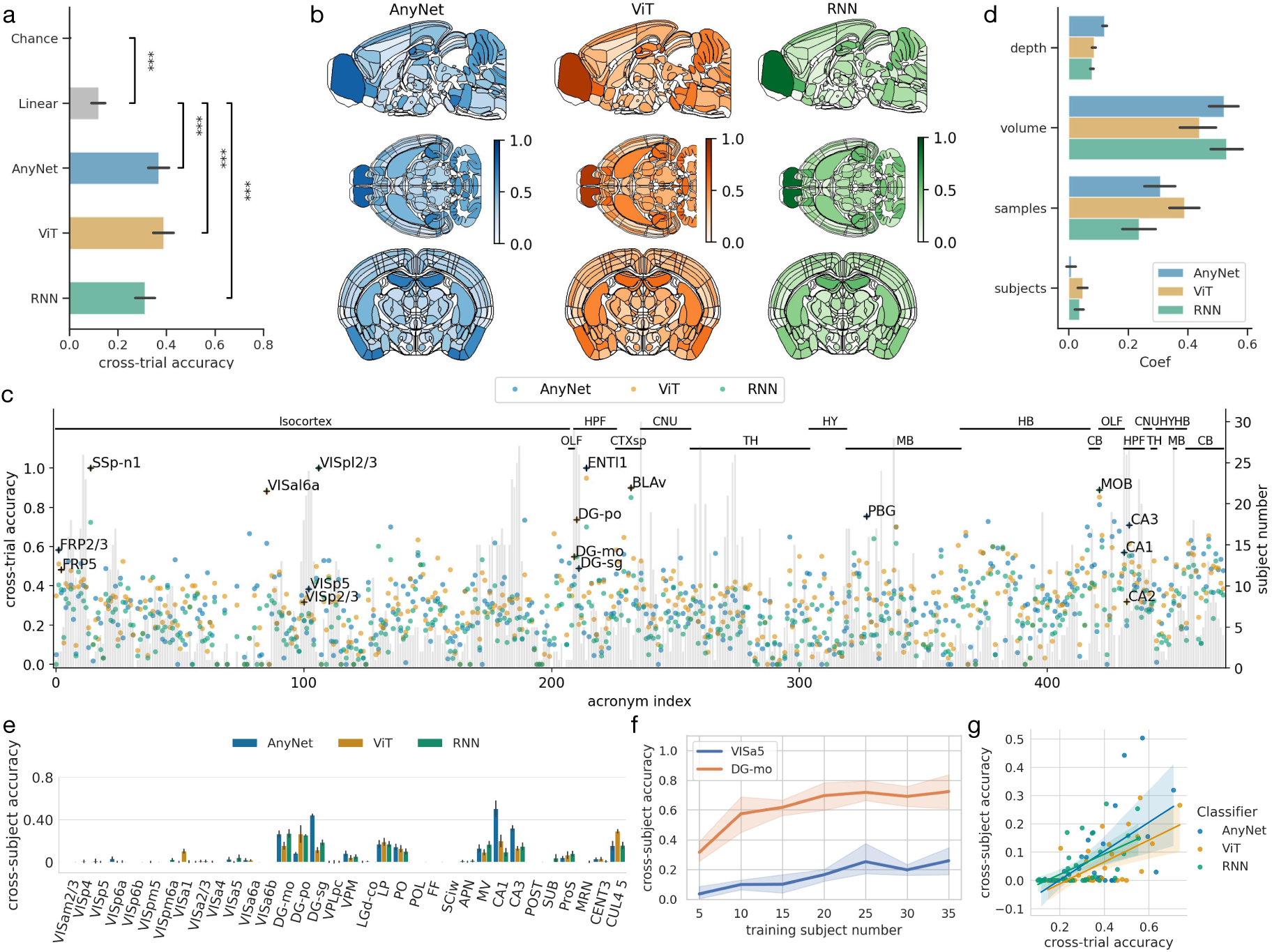
Electrospectral landmarks show high accuracy across trials and subjects. **a.** Overall cross-trial accuracy from distinct classifiers: linear network model, AnyNet, ViT, and RNN. Linear classifier performs significantly better than the chance level (*p <* 0.001). The three deep networks (AnyNet, ViT, and RNN) outperform the linear classifier (*p <* 0.001). **b.** Beta coefficients of multi-linear regression of cross-trial accuracy against depth, volume, samples and number of subjects. **c.** Spatial distribution of individual cross-trial accuracy from three classifiers across the mouse brain. **d.** Individual cross-trial accuracy (left axis) of the three deep learning models for the 472 brain regions. At the top of the figure is the label of the anatomical divisions. The subject number of each brain region is shown as gray bars referring to the right-axis. **e.** Cross-subject accuracy for brain areas having at least have 10 subjects. **f.** AnyNet cross-subject accurcay while holding out 5 subjects and gradually increasing the subject number in the training dataset. **g.** Scatter plot of cross-subject against cross-trial accuracies. Pearson Correlation between cross-trial accuracy and cross-subject accuracy are AnyNet *r* = 0.587, *p* = 0.00027; ViT *r* = 0.334, *p* = 0.0537; RNN *r* = 0.501, *p* = 0.00252. See Supplementary Table 1 for acronym definition.

To gain insights on the type of features exploited by the models, we also trained our models to classify time-averaged spectra instead of time-resolved spectrograms (Suppl. Fig. 3). In this case, the linear classifier upkept its accuracy (0.119), but the accuracy of the deep models dropped (AnyNet 0.213, ViT 0.357, RNN 0.176). This indicates that information increases from linear features to nonlinear feature combinations of the time-averaged spectra, and increases further with nonlinear feature combinations of spectrograms.

### Deep learning reveals multiple electrospectral landmarks

We then asked what makes one area more recognizable than another. Plotting the accuracy area-by-area (Fig. 2b,c), we found a considerable variability. We performed a multivariate regression of the region-specific accuracy on four factors: i) the volume of the brain area, ii) the number of samples (Fig. 2d), iii) the recording depth, which could decrease the accuracy via an increased stereotactic error but could also indicate more recognizable deeper structures either intrinsically or by bearing less artifacts of the surgery, and iv) the number of subjects. We found that the main factors affecting the cross-trial accuracy are the volume and the sample number, while the depth and subject numbers have comparatively negligible contributions (Fig. 2d). Surprisingly, depth was positively associated with accuracy, suggesting that stereotaxis errors are small and deeper are more recognizable. These factors, however, could not account for the majority of the variability (regression R^2^=0.14). Therefore, the considerable variability in localization accuracy is not fully accounted for by the size and number of samples and supports the idea that some areas are intrinsically more recognizable than others.

We call those more recognizable areas ‘landmarks’. Examples from the cross-trial test are the main olfactory bulb (MOB), cornus ammonis 1 (CA1) and the different layers of the dentate gyrus (DG; Fig. 2b,c). Turning to the cross-subject test, here we expect that variability in brain morphologies will necessarily reduce the maximal accuracy. Also, since most areas were not sampled consistently across animals, we could only perform this cross-subject test for a fraction of the areas (see Methods). Cross-subject accuracies for the remaining areas are shown in Figure 2e. A number of cross-subject landmarks can be identified by virtue of their relatively high accuracy. These include CA1, CA3 and each of the three layers of the DG. Other than these areas known for their distinctive LFP, we found the lateral posterior nucleus (LP), the posterior complex (PO) of the thalamus, hindbrain’s medial vestibular nucleus (MV) and the cerebellum’s Lobule IV (CUL4 5).

Given the added difficulty of the cross-subject test, it remains possible that these accuracies mainly reflect the number of subjects. To verify this, we selected two areas, non-landmark VISa5 (visual cortex, anterior area, layer 5) and landmark DG-mo (dentate gyrus, molecular layer), and gradually increased the subject number in the training set while testing in held-out subjects (Fig. 2f). As expected, the accuracy gradually increased with the number of animals in the training set. Yet, VISa5 required 35 subjects to match the accuracy achieved by DG-mo with only 5 subjects, further supporting that DG-mo is a landmark of the LFP. To more broadly test whether a landmark identified in cross-trial tests is predictive of landmark status in cross-subject tests, we calculated the correlation between cross-trial and cross-subject accuracies (Fig. 2g). We found a positive correlation, thus supporting the idea that we can identify subject invariant electrospectral structure from cross-trial tests.

### Electrospectral localization transfers to human

We further investigated the idea that our models trained on mice can then be fine-tuned on limited data to localize LFP from human brains, indicating a shared electrospectral structure across species. To investigate this, we used a human Neuropixels dataset, which was recorded from the dlPFC and temporal lobe [16]. In the absence of a precise mapping to a coordinate framework, we considered 21 labels based on electrode identity and used our three classifiers as we did for mice. We compared two training approaches: i) training classifiers from randomly initialized weights on the human dataset and ii) training classifiers that were pre-trained on the large mice dataset while only allowing the penultimate layer weights to be changed. If there is a shared structure across species, we should observe a faster learning in the pre-trained group. The final accuracy, in contrast, is more likely to reflect the restriction of which weight is allowed to be trained. We found that classifiers pre-trained with mice data could learn much faster than their counterparts trained from a random initialization on the human dataset (Suppl. Fig. 4). The final cross-trial accuracy did not show a significant difference between the transfer learning and random initialization group. This result further supports the potential for generalization of the electrospectral features learned by the models.

### Electrospectral similarities reveal robust communities

Inspired by spectroscopy approaches that rely on spectral matching to identify the composition of a sample, we hypothesized that a group of areas may share electrospectral features even when LFP spectra do not show well-separated peaks. Because the relevant spectral features may mix nonlinearly, we rely on the ANN representations of electrospectra rather than the electrospectra to assess similarity (Suppl. Fig. 5a). We found that the three ANNs converged to a consistent similarity matrix (Suppl. Fig. 5b).

We thus applied an approach from graph theory to delineate shared spectral features. The generalized modularity index [44] uses the ANN’s confusion matrices to create an undirected weighted graph and then quantifies the degree to which a given community structure is unlikely to arise from a random graph. The index is defined with a hyperparameter called the resolution (see Methods), which biases towards big communities when small. Given a resolution parameter, we used a greedy modularity search [45] to find a clustering that maximizes modularity (Suppl. Fig. 5c). Furthermore, to ensure that we find a community structure that does not depend on the specific ANN architecture, we performed the modularity search on each network independently and then focused on areas that are at the intersection between the communities found using each classifier (Fig. Suppl. 5d), the ensemble communities.

Before diving into the structure of the resulting electrospectral atlas, we explored different resolution parameters. Suppl. Fig. 5e shows the optimal modularity as a function of resolution. As a rule-of-thumb for modularity-based approaches, the index should be higher than 0.3 [9, 46], which is achieved for resolutions higher than 0.5. As higher resolutions yield smaller communities, we checked the robustness of the communities at different resolutions by bootstrapping the similarity matrix (Suppl. Fig. 5f) and found a monotonically decreasing robustness with an overlap ratio at 75% when using a resolution of 1. We also computed the cross-trial and cross-subject accuracies for the classifications of communities found at different resolutions (Suppl. Fig. 5g-h). Here the resolution showed only a small effect on robustness, and only the non-modular resolution (0.4) was markedly more robust, an observation explained by the very large communities that bias the accuracy due to class imbalance (Suppl. Fig. 6a). The localization accuracies of communities matched those of the areas that were individually most recognizable (landmark areas in Fig. 2). The resolution parameter therefore controls a trade-off between low robustness and low modularity, such that a resolution of one offers a good compromise.

Next we sought to compare the localization accuracies of an atlas based on the electrospectral communities and one based on a coarse histological atlas. We thus trained our classifiers on an atlas of ten mouse brain regions (e.g. olfactory bulb, isocortex, see Fig. 3b). Referring to this map as the Cosmos Atlas [17], we found again that a linear classifier was more apt than chance but surpassed by deep networks (Suppl. Fig. 5i, Suppl. Fig. 7). The deep networks achieved a cross-trial accuracy around 0.65. We could then directly compare the Cosmos Atlas with the 10 most recognizable communities, thus matching the task difficulty. We found that even when separating the brain into smaller regions than Cosmos, it is easier to localize communities of the electrospectral atlas (Suppl. Fig. 5j).

**Figure 3:**
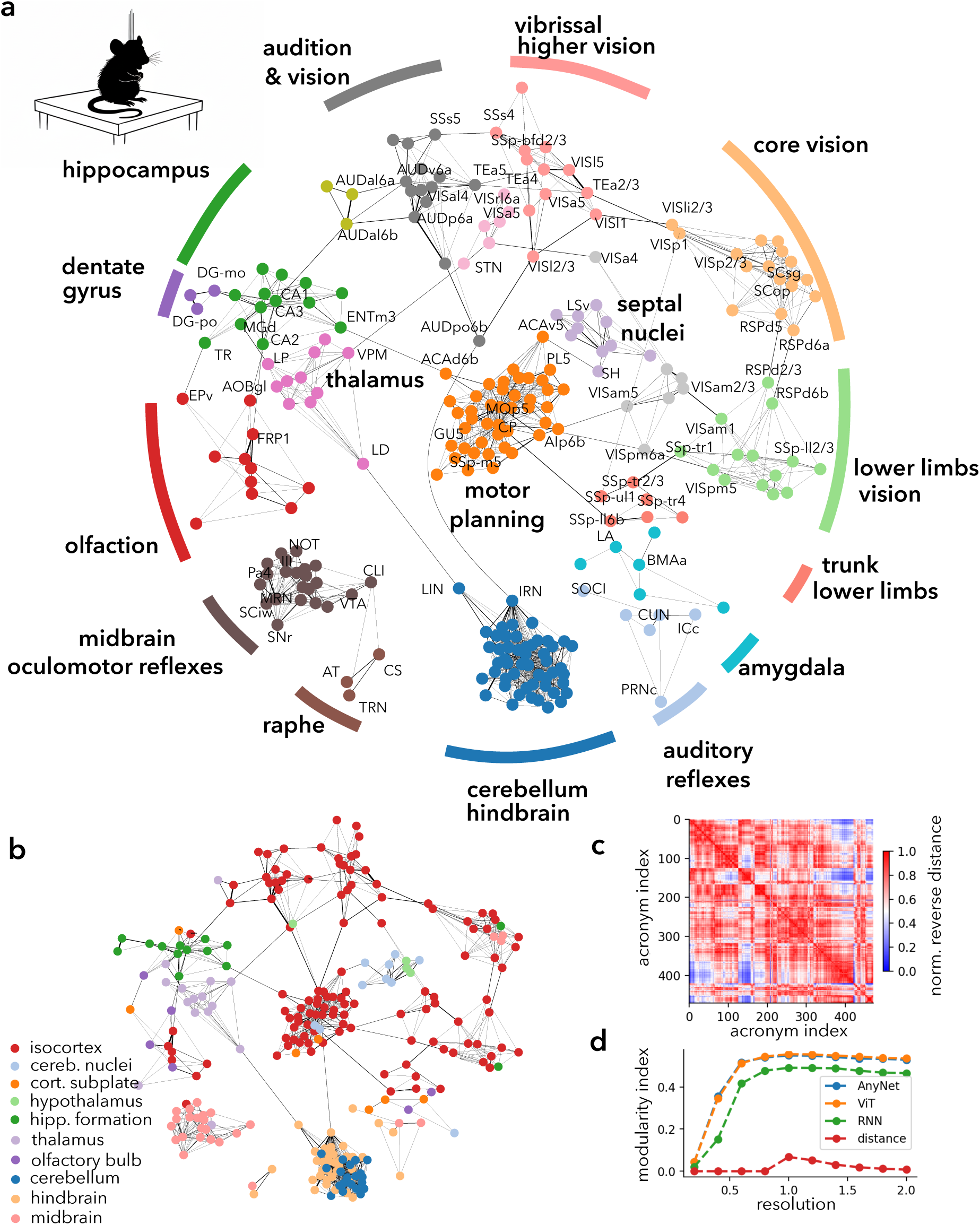
Electrospectral communities during the inter-trial period. **a** A selection of electrospectral communities (different colors) identified during the inter-trial state. Each circle represent an Allen atlas area. Graph links represent thresholded elements of the confusion matrix, with thicker lines corresponding to higher values of the confusion matrix. **b** Same graph as panel a, but colors representing coarse histological atlas. **c** Matrix of reversed normalized distances between area centers. **d** Comparison of modularity indices with the deep learning algorithms on electrospectra with those obtained from distances between areas centers. The modularity index mean and standard deviation are obtained by bootstrapping. Standard deviations are not visible because they are smaller than the marker size.

### Electrospectral communities of the intertrial period

Focusing on a selected set of communities, Figure 3 shows the resulting network graph with communities denoted by different colors (see also exemplar spectrograms in Suppl. Fig. 8 and full list in Suppl. Table 1). We stress that the links in the graph denote spectral similarity, not synaptic connections between areas. Graph theory analysis defines hub and connector nodes. Defining a hub node as the node with the largest in-degree, we looked at the probability that a given cortical layer was a hub (Fig. 15; Table 3). We found a layer specific probability where L6a and L5 were most likely and L4 and L6b the least likely. Performing the same analysis for connector nodes did not show a significant layer specificity (Fig. 15).

Many communities suggested an alignment with function. For instance, we found an ‘audio-visual community’ composed of the union of auditory cortex, parts of higher-order visual cortex, temporal and parietal association areas, a ‘hippocampus community’ which regrouped CA1, CA2, CA3 with entorhinal cortex, the ‘olfactory community’ regrouped parts of the olfactory bulb with frontal pole and orbitofrontal cortex, or a ‘visuo-vibrissal community’ that unites lateral and anterior visual cortex with temporal association areas and barrel fields. The area at the top of the visual hierarchy [13], labeled ‘VISam’, formed a relatively isolated community. The alignment with function was also suggested by subcortical communities such as a ‘auditory reflexes community’, which united superior olive (SOCI), inferior colliculus (ICc), a nucleus for the acoustic startle response (PRNc), and one for the tensor tympanic muscle (PC5). The ‘amygdala community’ regrouped the lateral amygdalar nucleus (LA) with the piriform amygdalar area of the olfactory bulb (PAA) and the cortical amygdalar area (COA). Alignment with function was not manifest for all communities, such as the large ‘cerebellum-hindbrain’ community, or the union of three small and tegmental areas we called ‘raphe’ (Fig. 3a).

At times, however, communities were not strictly circumscribed by the broad cytoarchitectural regions of the brain (Fig. 3b). For example, the putamen was the hub node of a community that unites nucleus accumbens, gustatory cortex, parts of the motor, somatosensory, anterior cingulate and insular cortex. Because many of these areas are part of the cortico-basal-ganglia loop, receiving inputs from the parafascicular nucleus, we labeled this community ‘motor planning’. The layer resolution of the LFP allowed to match specific layers of the colliculus with layers of the cortex: The ‘core visual community’ contained the optic parts of colliculus, primary visual cortex, parts of retrosplenial cortex and area prostriata, but not many other associative parts of the visual cortex, nor the other layers of the colliculus. The motor parts of the superior colliculus were instead separated into two communities (Suppl. Fig. 10), one containing many hypothalamic areas, and the other containing oculomotor parts of the midbrain. Therefore, we find that electrospectral communities can deviate from the cytoarchitecture in a way that suggests an alignment with function.

### Electrospectral communities reshape according to the task

To establish the community structure in a different context, we used the other set of models trained on the 1.7 M LFP samples of the task period. The IBL task required visual and auditory acuity, lower and upper limb coordination for turning a wheel and licking for obtaining rewards [18, 36, 2]. The communities were established anew using the same resolution parameter (Fig. 3c, exemplar spectrograms in Suppl. Fig. 9, and subsystems task-intertrial comparisons in Suppl. Fig. 10 - Fig. 13). We found that some communities appeared (a union of lower and upper limbs), some disappeared (visuo-vibrissal, visuo-tactile), some communities stayed approximately the same (visuo-auditory, visual-am, septal nuclei, amygdala) and others reshaped by adding or removing areas for a set of core areas.

Consistent with the appearance of the drifting grating, vision-related communities showed much reshaping. The core visual community increased in size (17 areas in the intertrial period vs. 25 in the task), with the inclusion of visual areas VISpm and VISa and reducing areas from VISli and VISl. The visual parts of the colliculus and area prostriata remained in this community, which also remained centered on layer 5 of the primary visual cortex (VISp5). Plotting the network graph of all communities containing subcortical visual and auditory areas (Suppl. Fig. 10), we observed that SCsg (a visual layer of superior colliculus) preserved some similarity with SCiw (intermediate white layer). SCiw belongs to a large community that contains motor and association parts of the superior colliculus along with most of midbrain (including oculomotor areas) and a large portion of thalamus, a merger of two communities of the intertrial period. To verify that this merge was significant, we performed a bootstrap test on probability of selected areas to be in the same or a different community (Fig. 4b, see Graph Theory Methods). Interestingly, a new small community appeared that contained both the visual and auditory relay thalamus despite these areas being dominated by categorically different afferents. These relay areas did not belong to any community during the intertrial period. Together, the picture that arises is that much reshaping of communities took place, both in terms of cortical and subcortical areas, at least around areas connected to the task.

**Figure 4:**
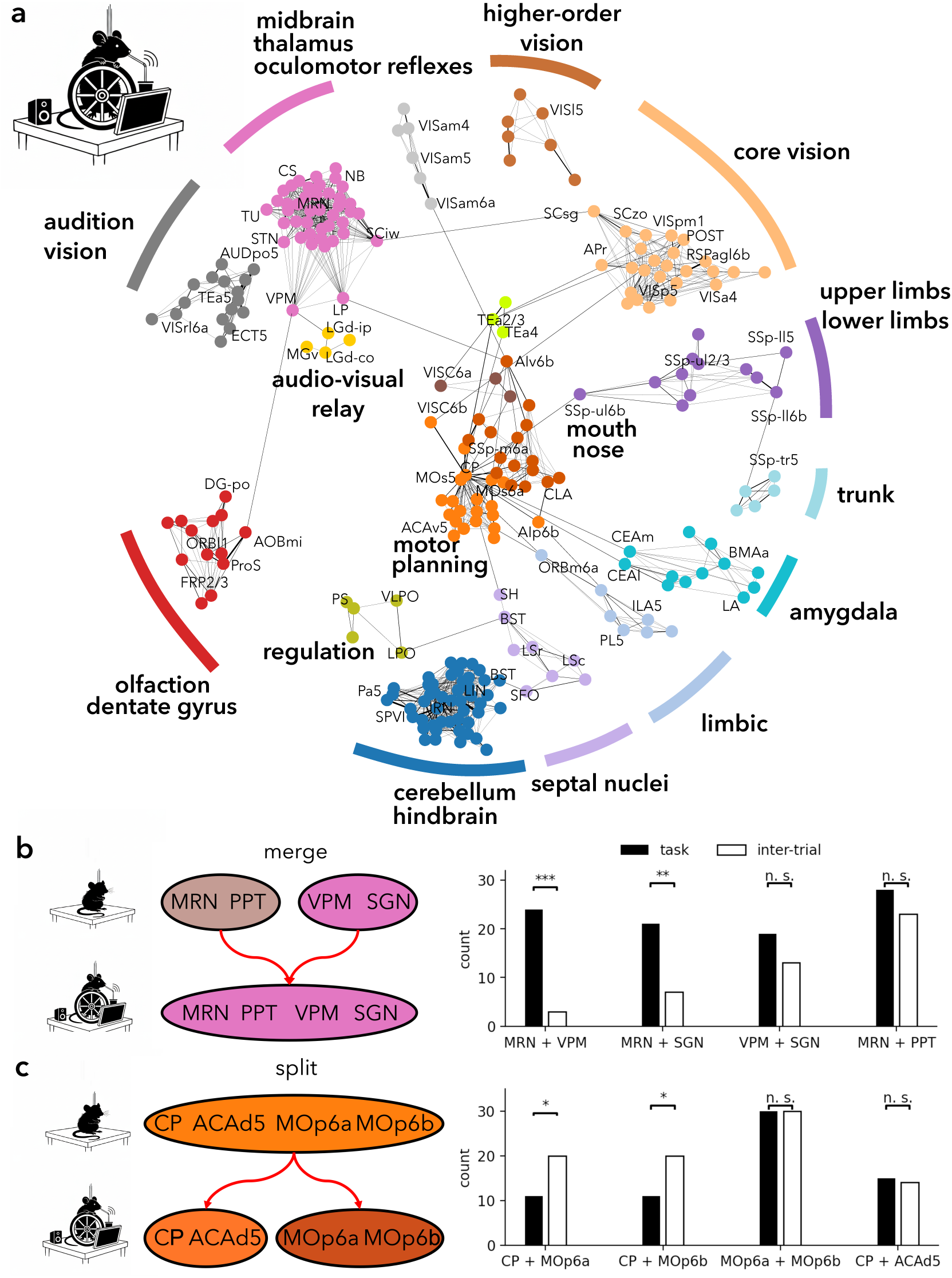
Electrospectral modules recombine during the task. **a** A selection of communities obtained during the visuo-motor task, as in Fig. 3. **b** Bootstrap test using Fisher’s exact method for computing the probability that two communities are in the same community in the task (black bar) and during the intertrial period (white bar). The MRN is in the same community as PPT but not VPM nor SGN during the intertrial period, while it is also in the same community as VPM and SGN in the task period, indicating a merge of communities in the task period. **c** Same as b, but for areas CP, ACAd, MOp6a and MOp6b, indicating a split of communities in the task period.

Potentially reflecting the licking and wheel-maneuvering components of the task, we found that motor and planning areas were marked by a split. During the intertrial period, the motor planning community centered on the putamen and contained the nucleus accumbens (ACB), frontal pole (FRP), anterior cingulate area (ACA), the secondary motor cortex (MOs), agranular insula (AI), primary motor cortex and orofacial somatosensation (SSp-n, SSp-m). During the task, putamen was still part of the community that integrated ACA, MOs and ACB, but the orofacial and primary motor areas formed a distinct community. We performed a bootstrap test to verify the significance of this split (Suppl. Fig. 5c). Interestingly, while most similar to the hippocampus or the septal nuclei community during the inter-trial state, the motor planning community became most similar with the visual and limbs communities. Together, these findings support that the LFP is organized in modules which can reconfigure with behavior.

## Discussion

Brain-wide brain wide graphs of Suppl. Fig. 5 and 3 appear centered on the caudoputamen, a subcortical structure involved in motor control, emotion and cognition. Graphs that contain such hub structure are naturally highly sensitive to subsampling. Therefore, we expect that an even wider sampling of areas can alter the precise hierarchical graph that can be inferred from the LFP. Similarly, there different hyperparameters or different modularity search will likely provide a slightly different community structure.

The default mode network is a group of areas identified in humans that are active specifically during the resting state. One may ask if any of the electropectral communities matches the mouse DMN. Following Whitesell et al. (2021) [**?**], the cortical part of mouse core DMN regroups areas along the midline (ACAd, ACAv, PL, ILA, ORBl, ORBm, ORBvl, VISa, VISam, RSPd, RSPv, SSp-tr, SSp-ll and MOs). Except for the sensory areas (RSP, SSp and VIS), these areas are all found in the motor planning community. Therefore, the largest cortical community aligns substantially with the default mode network. This community splits during the task, when the default mode network is inhibited.

Given that the LFP is shaped by both the activity patterns of the afferents and those in the local population [19], one may ask if either of those factors influence more the community structure. If the structure of the LFP was dominated by afferents, we would expect that the electropectral community should reflect the modularity of the connectivity. Previous research has revealed community structure in the connectivity of intracortical connections [13, 12], and these clearly show a high degree of overlap with the electrospectral communities. If, alternatively, the LFP was dominated by local properties, we would expect that the local transcriptome would reflect the modularity of the electrospectra. Again here, the modularity structure of the local transcriptome shows a high overlap with that of the LFP. While these comparisons are ambiguous, it is worth to note that our findings indicate a dynamical structure of the LFP. Within the connectivity and cytoarchitectural constraints, the LFP re-organizes with the behavior, with layer precision, both across cortex and subcortical structures.

## Supplementary Materials

### Methods

#### International Brain Lab data

We used brain-wide Neuropixels recordings from International Brain Lab (IBL) [47], which was recorded across 139 mice in 12 labs. During the experiment, the mouse did a visual discrimination task requiring it to turn a wheel to move a visual stimulus (an oriented Gabor having variable contrasts) to the center of the screen. The mouse needed to hold the wheel still for 0.4-0.7 s to start a new trial, which was announced by an auditory cue. All mice were first trained and rewarded with sweetened water when they made correct decisions. Wrong decisions would bring a noise burst and longer inter-trial interval (2 s) as a punishment. Before the visual stimulus was turned off, the mouse had enough time for collecting the reward. Additional information for surgical methods, experimental materials, and probe tracking and alignment can be found in [18, 48]. All animal experiments were conducted according to local laws and approved by the institutional ethics committees [47]. Details for mice like housing, weights, and genders can be found in [47]. To collect samples from as many areas as possible brain areas, we used brain areas in Beryl mapping in case some recorded areas couldn’t be localized precisely under the Allen brain atlas (e.g., CA1so, CA1sp and CA1sr were labeled as CA1). We didn’t distinguish between hemispheres. The time period of interest was 1 second. For the intertrial period it is immediately after the visual stimulus was turned off and for the task period it is immediately after the visual stimulus was turned on. In some rare cases, the inter-trial time interval was smaller than 1 second, such that the inter-trial period contained part of this stimulus period. This only occurred on 0.06% of the trials. In all, we collected 3.4 million LFPs (≤ 500 Hz) samples from 472 brain areas for 1 second in the inter-trial and the task states.

#### Data preprocessing

##### Spectrogram

For each 1 second sample of the LFP, acquired at 2500 Hz sampling frequency, we performed a Fast Fourier Transformation (FFT) with 401 bins, window length 800 sample points (320 ms), and hop of window length 400 sample points (160 ms), which results in frequency resolution of 3 Hz. We limited the frequency for analysis between 3 and 500 Hz. This results in a time-resolved spectrum (spectrogram) of dimension 224 × 28, where the first dimension is the frequency and the second dimension is time. We used the square root of the power so as to better preserve activity in the ripple band and beta band. This frequency range should still bear signatures of the action potentials nearby [**?**] and harmonics of rhythms slower than 3 Hz. The time-resolved spectra should also still keep partial information about rhythm nesting.

##### Cross validation

For each brain region, we assembled a dataset from multiple subjects. For each subject, we kept only the first 80 trials to help balance data across subjects. As a training set, we used only the first 60 trials and only for a fraction of the animals. The following 10 trials of each subject constituted the validation test. The last 10 trials of the same animals made the cross-trial test set. Some animals were held out to constitute the cross-subject test. To determine which animals were held out and which were used for the training set, we calculated the subject number for samples of each recorded brain area. We only held out subjects that would not make the sampled subject numbers for each recorded area less than 10. The cross-subject test set is then made up of the first 80 trials in the first experiment of each held-out subject.

##### Human data

We used publicly available Neuropixels recordings in humans, which were acquired from the dlPFC and the temporal lobe [16]. To refine the recording location, for one penetration in one coarse brain region, which had recordings from 384 channels, we only preserved one channel every 60 channels and labeled it as a new brain area index. Finally, we got 21 refined brain regions in all. All preprocessing from LFPs to the input spectrogram of the human data was the same as mice data. For each fine brain region, we had 2 000 samples, each for 1 second. We used the first 1400 samples for training, the next 200 as validation, and reserved the last 400 for testing. In all, we had 42 000 labeled samples of human LFP.

#### Neural networks

##### Classifier architectures

The Recurrent Neural Network (RNN), the Convolutional Neural Network (CNN), and the Transformer are three basic types of modern deep network architectures. The RNN focuses on extracting sequential information with the risk of ignoring global information. The CNN extracts the local features and assembles them into a global representation by the convolution process. The self-attention mechanism in the Transformer enables the algorithm to integrate global sequential information; its quadratic computational complexity makes the training expensive.

For the RNN architecture, we treated the size of the frequency dimension of the input as the input size and the time dimension as the sequential dimension, as shown in Suppl. Fig. 6. We chose a two-layer, gated recurrent unit (GRU) module with 512 hidden units per layer. The last dimension of the output from the GRU module [49], which integrates previous sequence information, is fed into a linear layer that had the same number units as the number of brain regions. The RNN is not bi-directional because of our causal assumption of the spectrogram.

For the CNN architecture, considering the input as an image, we implemented the AnyNet framework to design our model [40]. AnyNet is a semi-auto neural architecture design framework for computer vision tasks, as shown in Suppl. Fig. 6. AnyNet consists of a stem, a body and a head. The stem was used to implement the initial processing for the spectrogram; multiple blocks exist in the body to generate the final representation by sequential nonlinear transformations; the head uses the linear readout to convert the output representation from the body to the brain region category. The architecture parameters include depth *d_i_*, the number of the output channels *c_i_*, the number of groups *g_i_*, and bottleneck ratio *k_i_*for stage *i*. The AnyNet design principles are the bottleneck ratio *k_i_*= *k*, group width *g_i_* = *g* for all stages *i*; increase network width, depth across stages, which *c_i_* ≤ *c_i_*_+1_, *d_i_* ≤ *d_i_*_+1_. In this proposal, considering the principles and the limitations of our hardware, we designed our AnyNet as having no bottleneck and the group width is always set to be 1, in which *k_i_* = 1 and *g_i_*= 1; the channel number *c*_0_ for the stem was 64, and the channel number for the remaining blocks was *c_i_*= *c*_0_ ∗ (*i* + 1); the depth for all stage was *d_i_*= 2.

For the Transformer architecture, we still treated the input as an image and used the Vision Transformer (ViT) model [42], as shown in Suppl. Fig. 6. We separated the input into 8 patches, with each patch size being 28 × 28. A *< cls >* token was also fed with 8 patches. After cascade transformation, its representation will be mapped into the output brain region label by a linear layer combined with the Softmax function. We implemented 8 heads and 4 Transformer encoder blocks in our ViT model.

##### Classifier training

As a typical method, we implemented cross-entropy to measure the loss between the label and model output for all three architectures. We used the Adam algorithm as the optimizer for training our models and set the batch size as 256. To avoid overfitting, the validation set was used for early stopping, in which we only retained models which had the best validation accuracy with training accuracy ≥ 0.6.

##### Computing resources

Our deeplearning based classifiers were trained by one A100 GPU. For an AnyNet classifier, training would require 4-6 hours, while for the ViT or the RNN, it required around 2 hours.

#### Graph Theory Methods

##### Greedy community search

For each classifier, we treated the confusion matrix as a weighted graph, allowing us to use theoretical tools from graph theory. To find latent communities based on the classifier’s confusion matrix, we implemented Clauset-Newman-Moore greedy modularity maximization. The modularity is calculated as 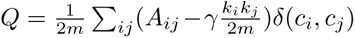, where *m* is the number of edges, *A* is the adjacency matrix, *k_i_* and *c_i_* the weighted degree of node *i*, *γ* is the resolution, and *δ*(*c_i_, c_j_*) equals 1 when *i* and *j* are in the same community otherwise equals 0. The greedy-based method begins with each node in the graph as its own community and repeatedly merged the pairs of communities that “lead to the largest modularity until no further increase in modularity is possible (a maximum)”. We performed this greedy search under different resolution (*γ*) conditions. We used the NetworkX package to complete above functions https://networkx.org/ [50].

##### Merging and splitting

To test community merge or split during distinct tasks, we examined the coexistence of chosen brain areas. We sampled 128 recordings of each brain area. (For a each brain area, if the number of samples was less than 128, we just used all samples we had). We regenerated confusion matrices 30 times. For those 30 confusion matrices, we performed greed community search as described above to obtain 30 community partitions. We counted the partitions for which the two brain areas were within the same community in task state. We repeated the computation for inter-trial states. We then used Fisher’s exact test to compute the probability of rejecting he null hypothesis that there is no difference of coexistence count between two states.

##### Hub and connector nodes

## Author Contributions

XW and RN designed the study, interpreted the results and wrote the manuscript. XW performed the computational analyses. JCB and RN acquired funding. RN supervised the project. JCB provided a verification of the approaches.

---

**Supplementary Figure 1:**
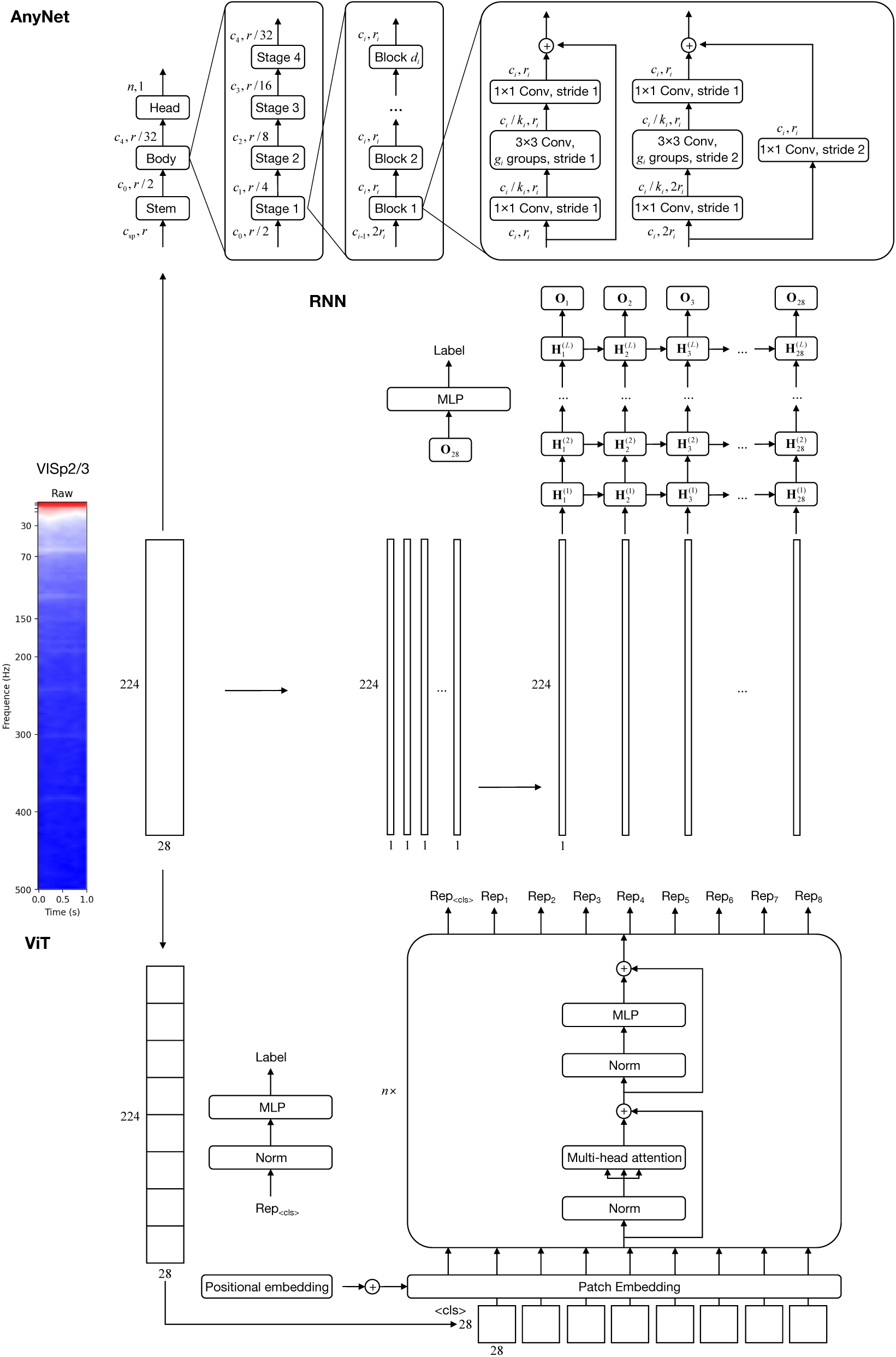
Architectures for the AnyNet, ViT and RNN. As shown above, for AnyNet, the input spectrogram is processed as a image and fed in the classifier; for ViT, the input is separated as several patches for the attention processing; for RNN, the input is cut as temporal slices and given to the classifier sequentially.

**Supplementary Figure 2:**
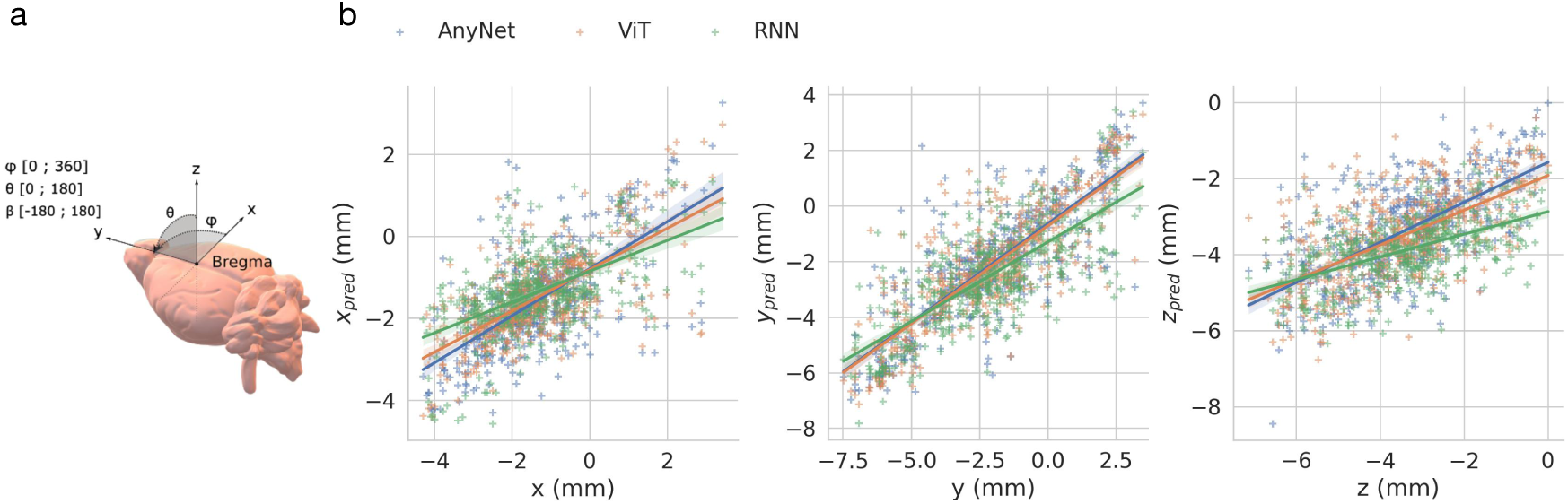
Deep learning-based regression for stereotactic coordinates using LFPs. **a.** Illustration of stereotactic coordinates in mouse brain. **b.** Regression for x, y and z axis using LFPs. Deep learning methods, AnyNet, ViT, and RNN are able to predict stereotactic coordinates. For x axis, AnyNet (*R*^2^ = 0.356*, r* = 0.656*, p <* 0.001), ViT (*R*^2^ = 0.433*, r* = 0.672*, p <* 0.001), and RNN (*R*^2^ = 0.2889*, r* = 0.553*, p <* 0.001). For y axis, AnyNet (*R*^2^ = 0.655*, r* = 0.812*, p <* 0.001), ViT (*R*^2^ = 0.683*, r* = 0.828*, p <* 0.001), and RNN (*R*^2^ = 0.527*, r* = 0.746*, p <* 0.001). For z axis, AnyNet (*R*^2^ = 0.395*, r* = 0.650*, p <* 0.001), ViT (*R*^2^ = 0.373*, r* = 0.622*, p <* 0.001), RNN (*R*^2^ = 0.197*, r* = 0.534*, p <* 0.001)

**Supplementary Figure 3:**
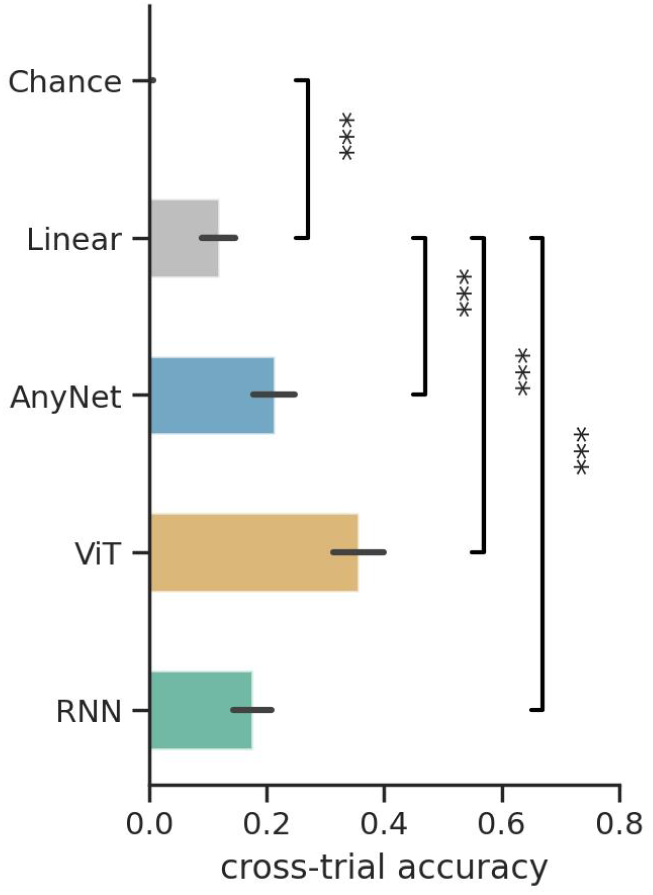
Classification using mean spectrogram of inter-trial state. Overall cross-trial accuracy from distinct classifiers: linear network model, AnyNet, ViT, and RNN. Linear classifier performs significantly better than the chance level (*p <* 0.001). The three deep networks (AnyNet, ViT, and RNN) outperform the linear classifier (*p <* 0.001). Overall cross-trial accuracy: linear model (0.119), AnyNet (0.213), ViT(0.357), and RNN (0.176).

**Supplementary Figure 4:**
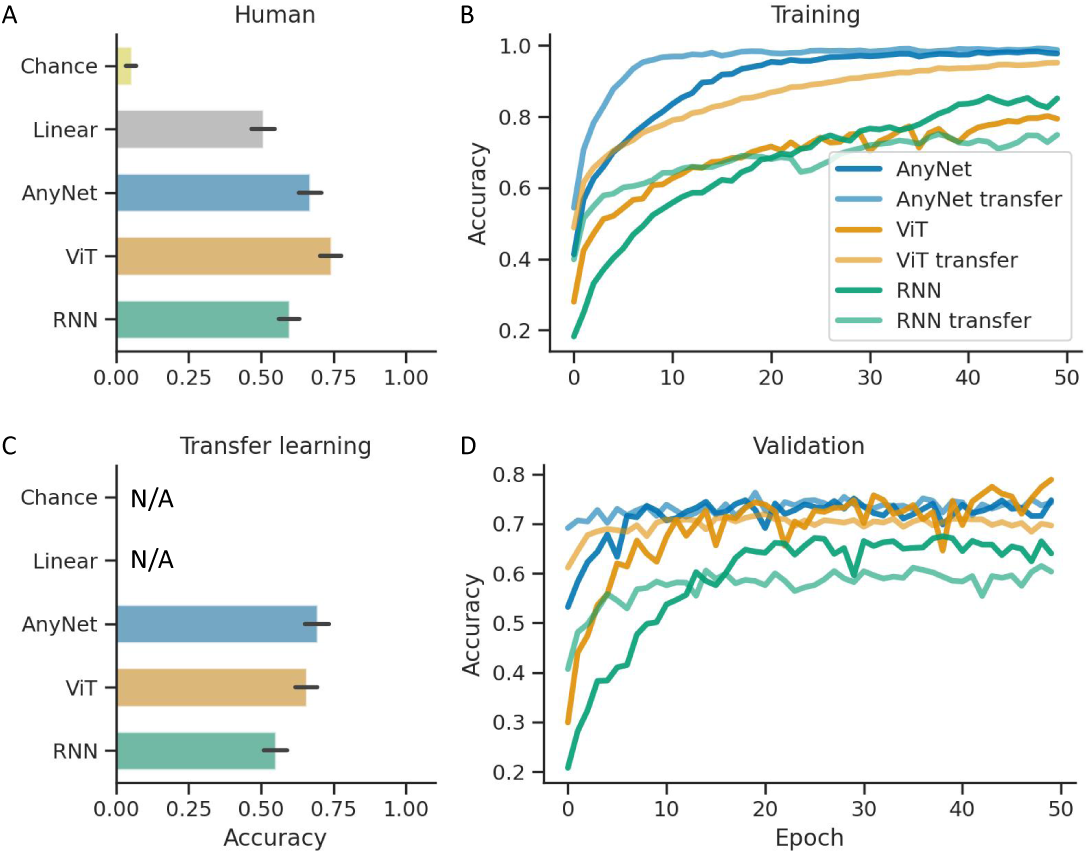
Pre-training on mouse data accelerates training on human data. **a.** Overall cross-trial accuracy for the human Neuropixels dataset trained from random initialization. **b.** Training accuracy of three classifiers for the transfer learning and training from the random initialization. **c.** Cross-trial accuracy for the transfer learning of three classifiers. **d.** Validation accuracy curves during training for three classifiers during transfer learning.

**Supplementary Figure 5:**
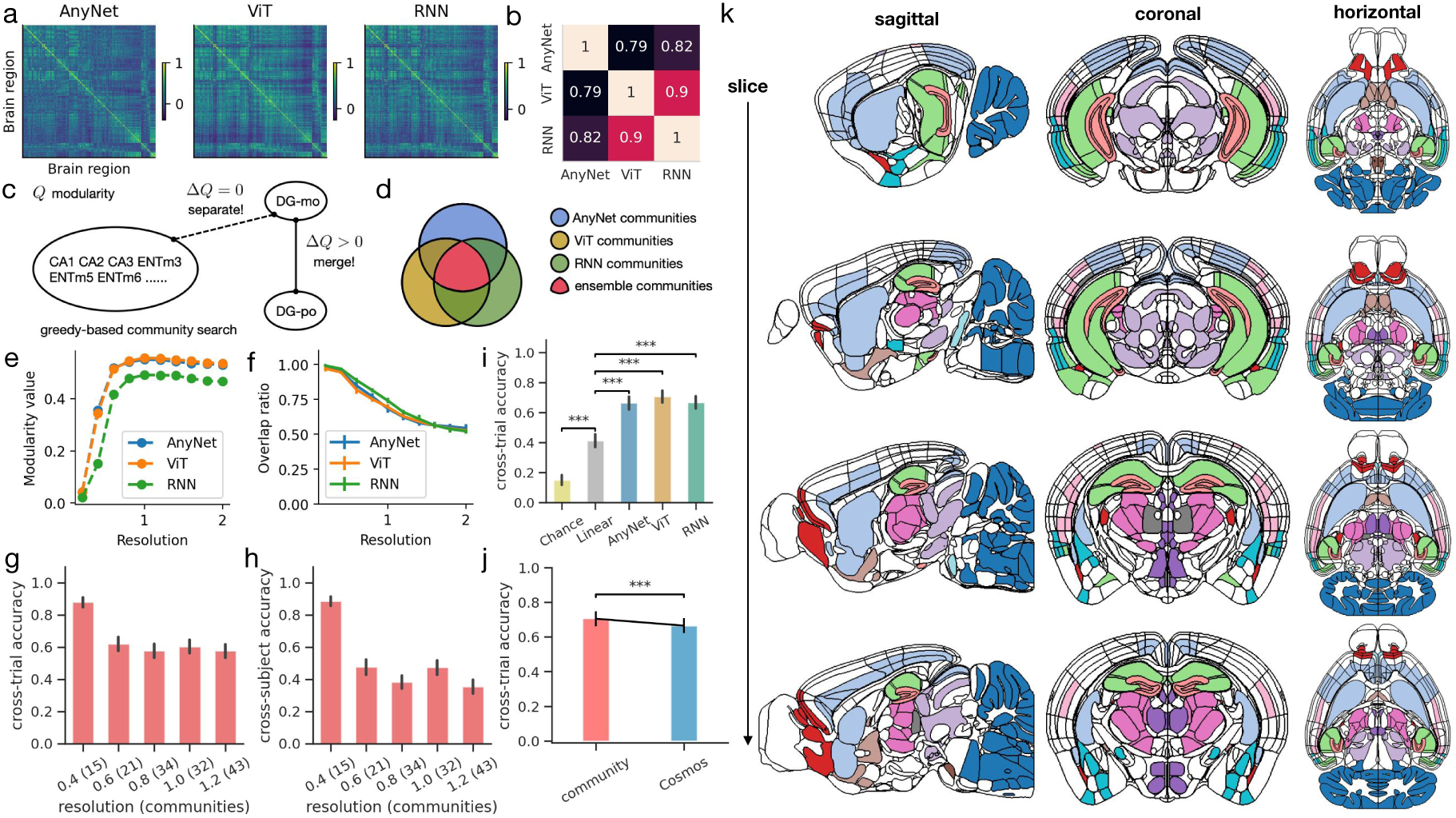
Spectral similarities define communities. **a**. Heat map showing the correlation coefficient of the classifier’s penultimate layer activations of one Allen brain atlas area to another. Ordering follows Fig. 2c. **b**. Similarity analysis for the correlation matrices of the three different models, each square shows the correlation coefficient between the matrices shown in a. **c**. Schematics of greedy modularity search for communities in 472 brain regions. If merging two separated communities increases overall modularity, then a new community is generated; otherwise, those two communities remain isolated modules. **d**. Schematic for establishing a consensus across ANNs. Ensemble communities are obtained by first aligning each community across ANNs and then only keeping the brain areas that were found to belong to that community consistently across the three ANNs. **e**. Final modularity values of greedy search is shown for different resolutions. **f.** The overlap ratio between communities found using all brain areas and communities using 80 % of the brain areas. **g**. Average cross-trial accuracy for ensemble communities at different resolutions, the number of communities is shown in parentheses. **h**. Same as g but for cross-trial accuracy. **i**. Overall cross-trial accuracy for the different algorithms trained to classify the 10-region anatomical atlas ‘Cosmos’. **j**. Comparing community with Cosmos at a matched number of 10 classes in terms of the cross-trial accuracy. **k**. Sagittal, coronal and horizontal slices showing 18 ensemble communities out of the 32 using different colors. Only 18 are shown for enhanced visualization, hence the white areas are either areas that did not belong to any community, or areas that were in the 14 communities that could not be shown simultaneously.

**Supplementary Figure 6:**
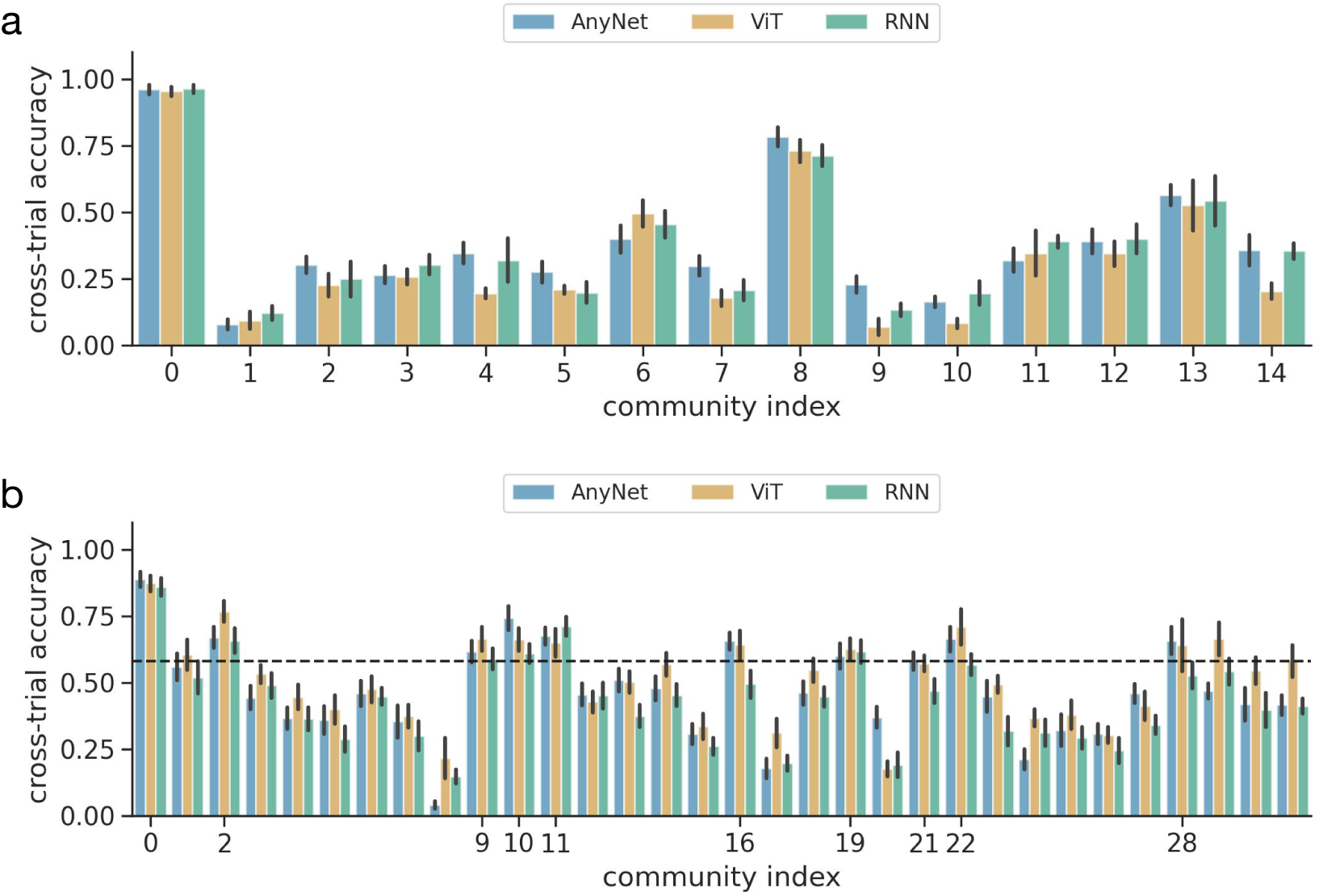
Individual cross-trial accuracy for communities under different resolutions. **a.** Individual cross-trial accuracy for communities under 0.4 resolutions. **b.** Individual cross-trial accuracy for communities under 1.0 resolutions. Communities which individual accuracy for AnyNet are in Top 10 (above the dash line) are selected as landmark communities.

**Supplementary Figure 7:**
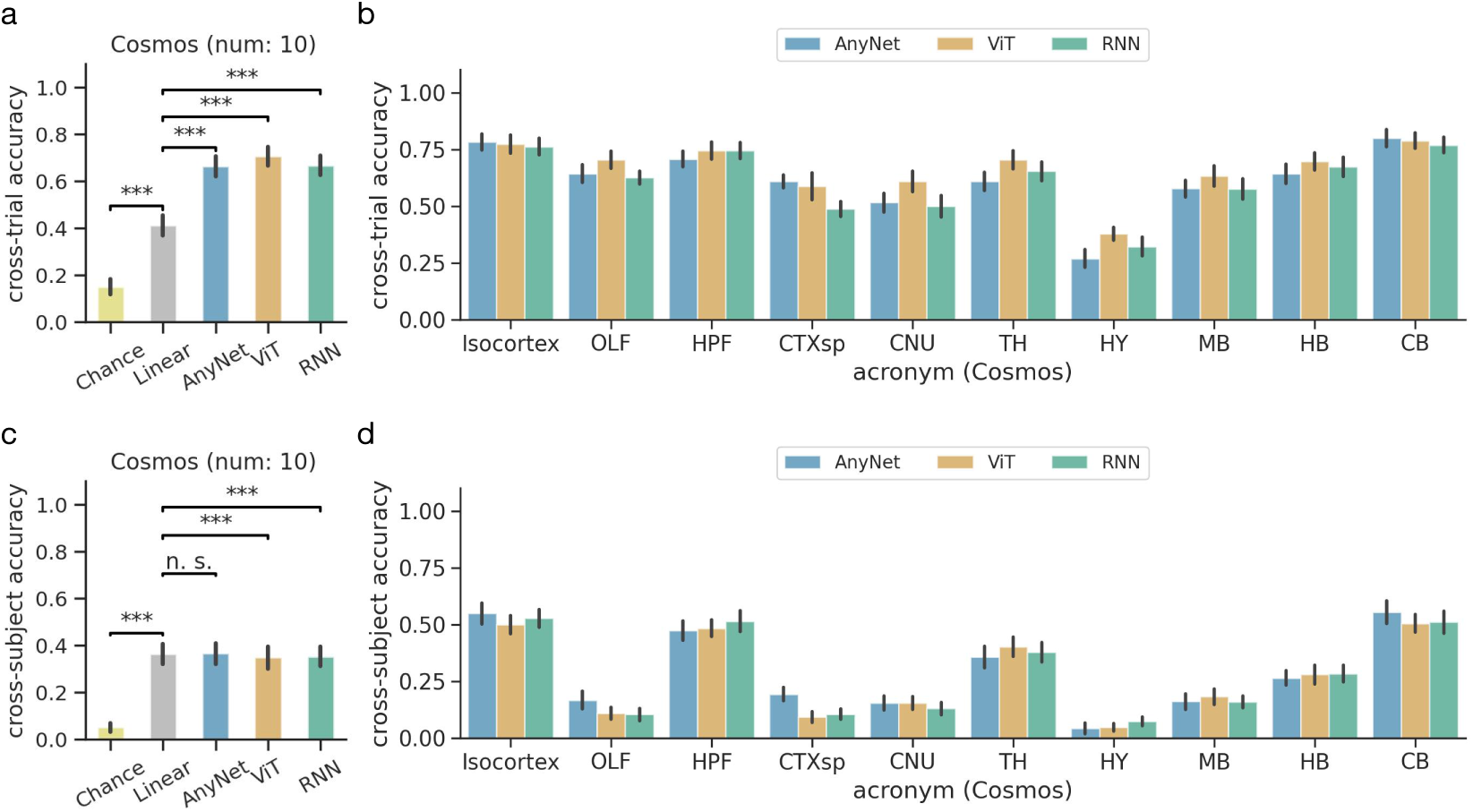
Classification of Cosmos mapping. **a.** Overall cross-trial accuracy for Cosmos mapping. **b.** Individual cross-trial accuracy for Cosmos mapping. **c.** Overall cross-subject accuracy for Cosmos mapping. **d.** Individual cross-subject accuracy for Cosmos mapping.

**Supplementary Figure 8:**
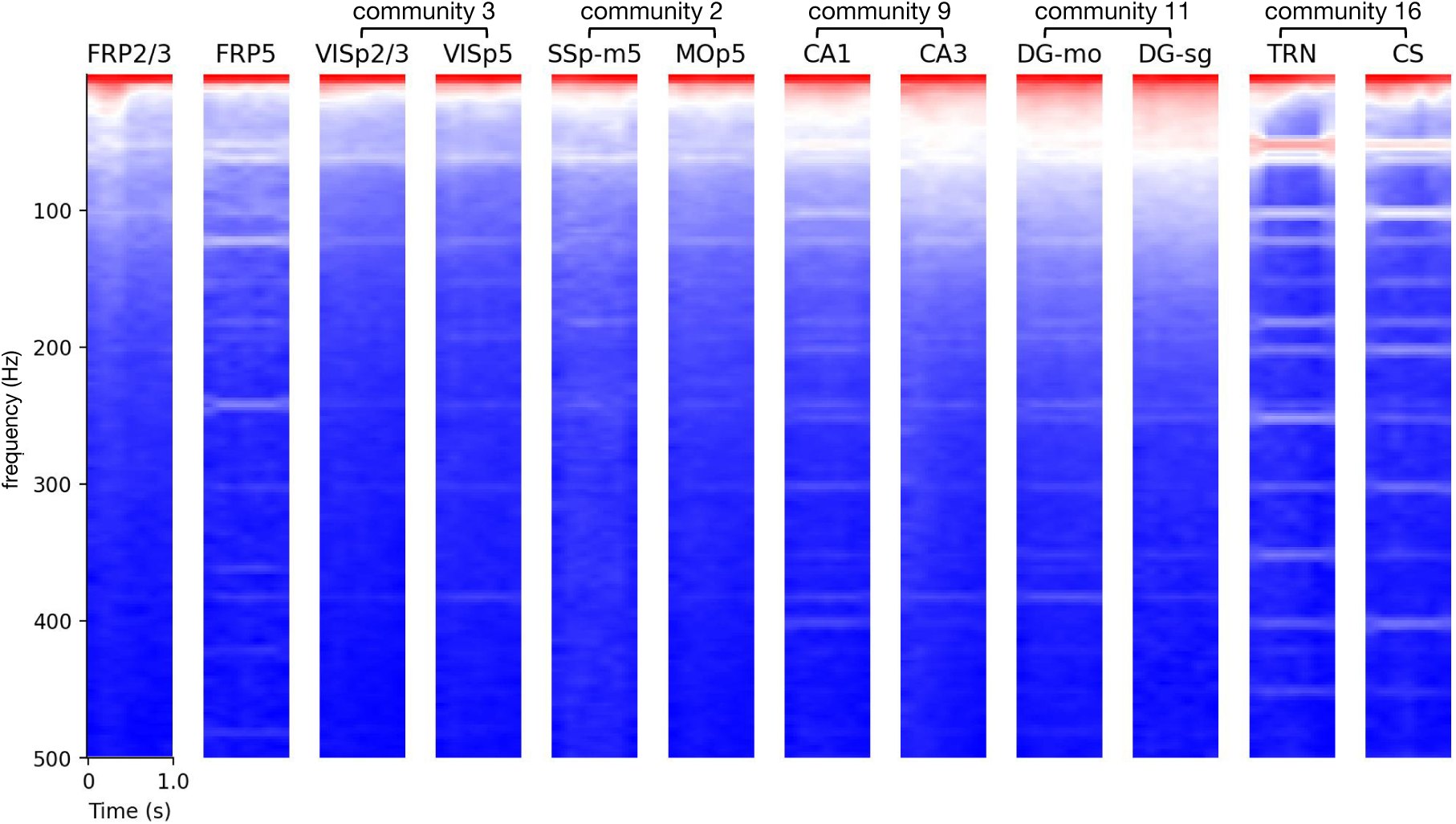
**Mean spectrogram of distinct brain areas of inter-trial state.**Power spectra were normalized and read out with a logarithm for visualization. Red means high amplitude; and blue represents low amplitude. Brain areas from the same community are marked. Much of the bar structure seen in, for instance, TRN arises from scaling a lower power signal that contained the alternative current artifact (at 50 Hz for European labs and 60 Hz for American labs) and its harmonics.

**Supplementary Figure 9:**
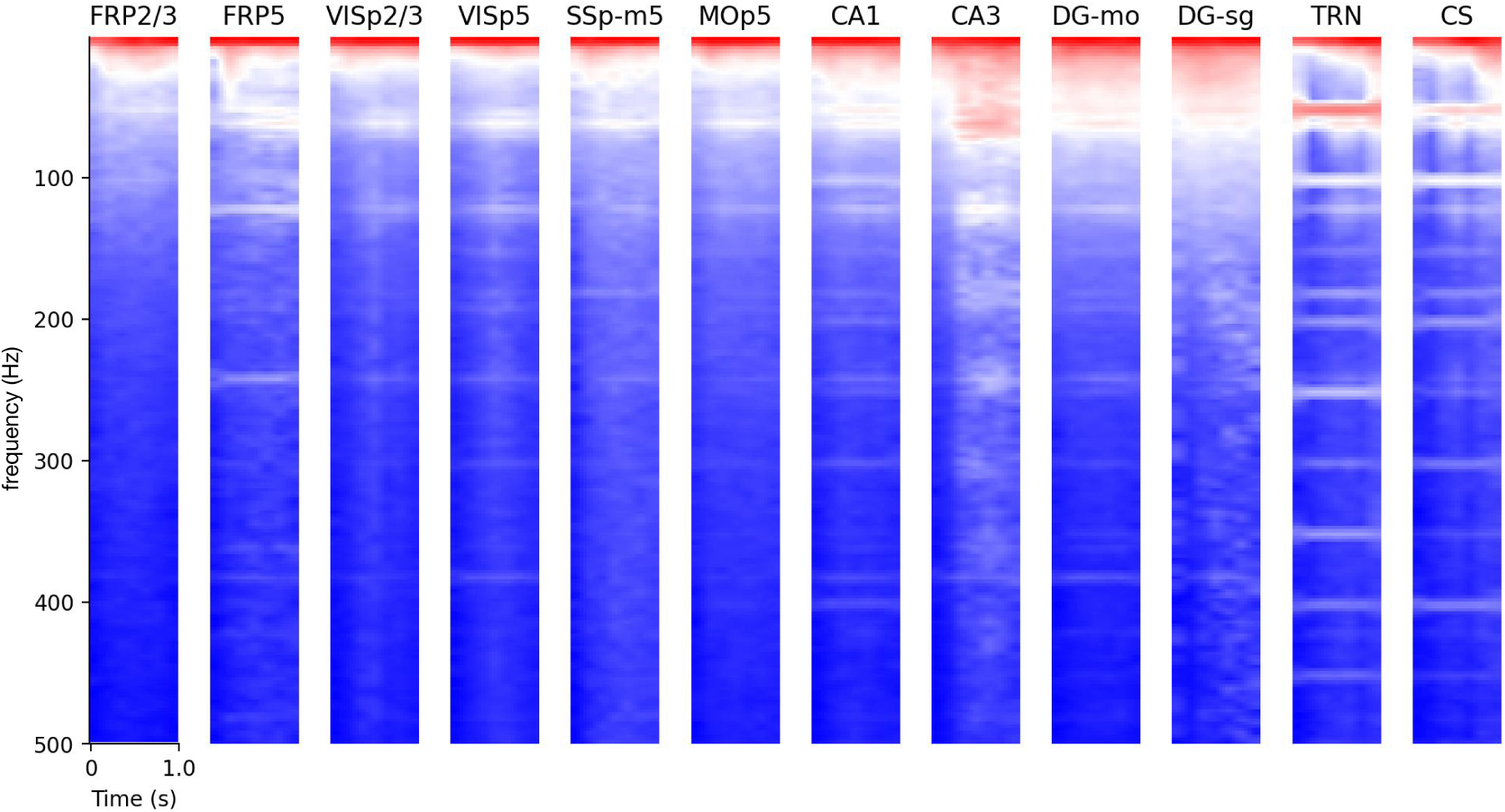
Mean spectrogram of distinct brain areas of task period. Same as Supplementary Fig. 8 we show the logarithm of the mean spectrogram of task period.

**Supplementary Figure 10:**
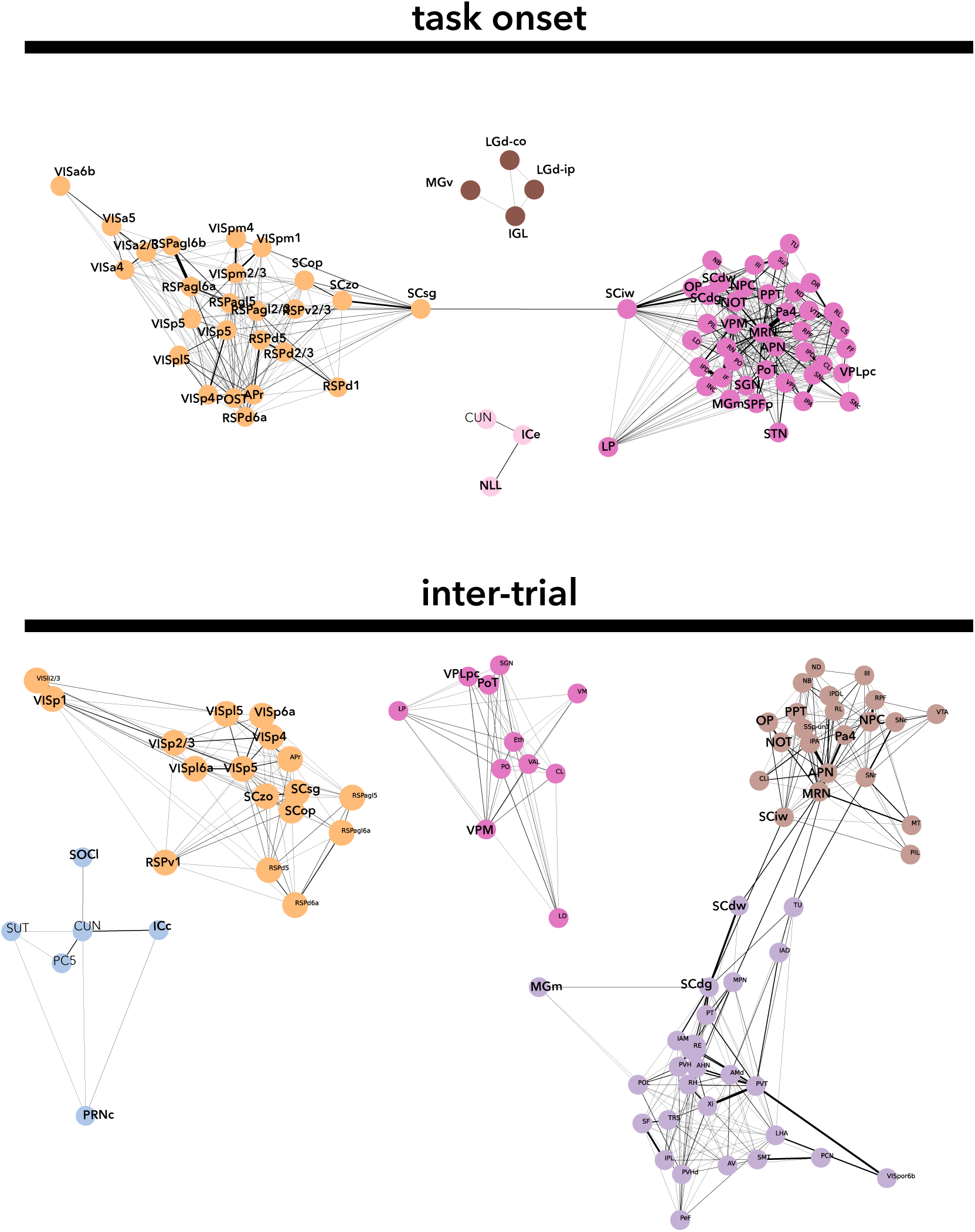
Electrospectral for basic audio-visual areas. Selection of communities containing areas associated with lower order subcortical visual sensory (lateral geniculate nucleus, LGd, superior colliculus, SCop, SCzo and SCsg), auditory (inferior colliculus ICc, medial geniculate nucleus, MGm, MGv, Nucleus of the Lateral Lemniscus NLL, superior olive SOCl), oculomotor (MRN, APN, Pa4, NOT, IP, PPT, superior colliculus, SCiw, SCdw and SCdg) and auditory reflexes (caudal part of pontine reticular nucleus PRNc) information. Graph links represent thresholded confusion matrix element with line thickness.

**Supplementary Figure 11:**
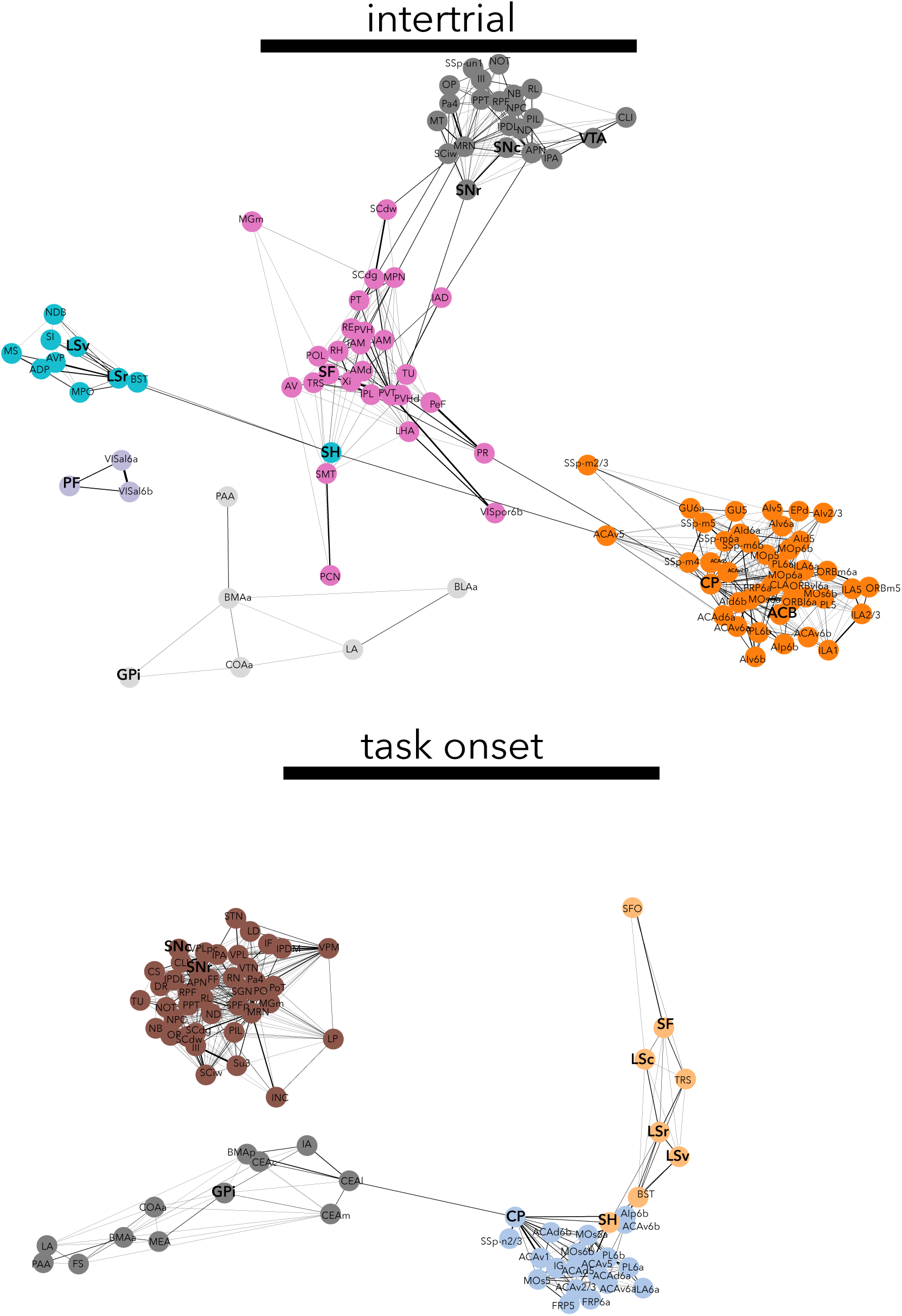
Electrospectral for kinematics system. Selection of communities containing areas associated with kinematics: putamen (CP), parafasacicular nucleus (PF), lateral septum (LSr, LSv), globus pallidus (GPi), nucleus accumbens (ACB), septofimbrial nucleus (SF), septohippocampal nucleus (SN), Substantia Nigra (SNr, SNc).

**Supplementary Figure 12:**
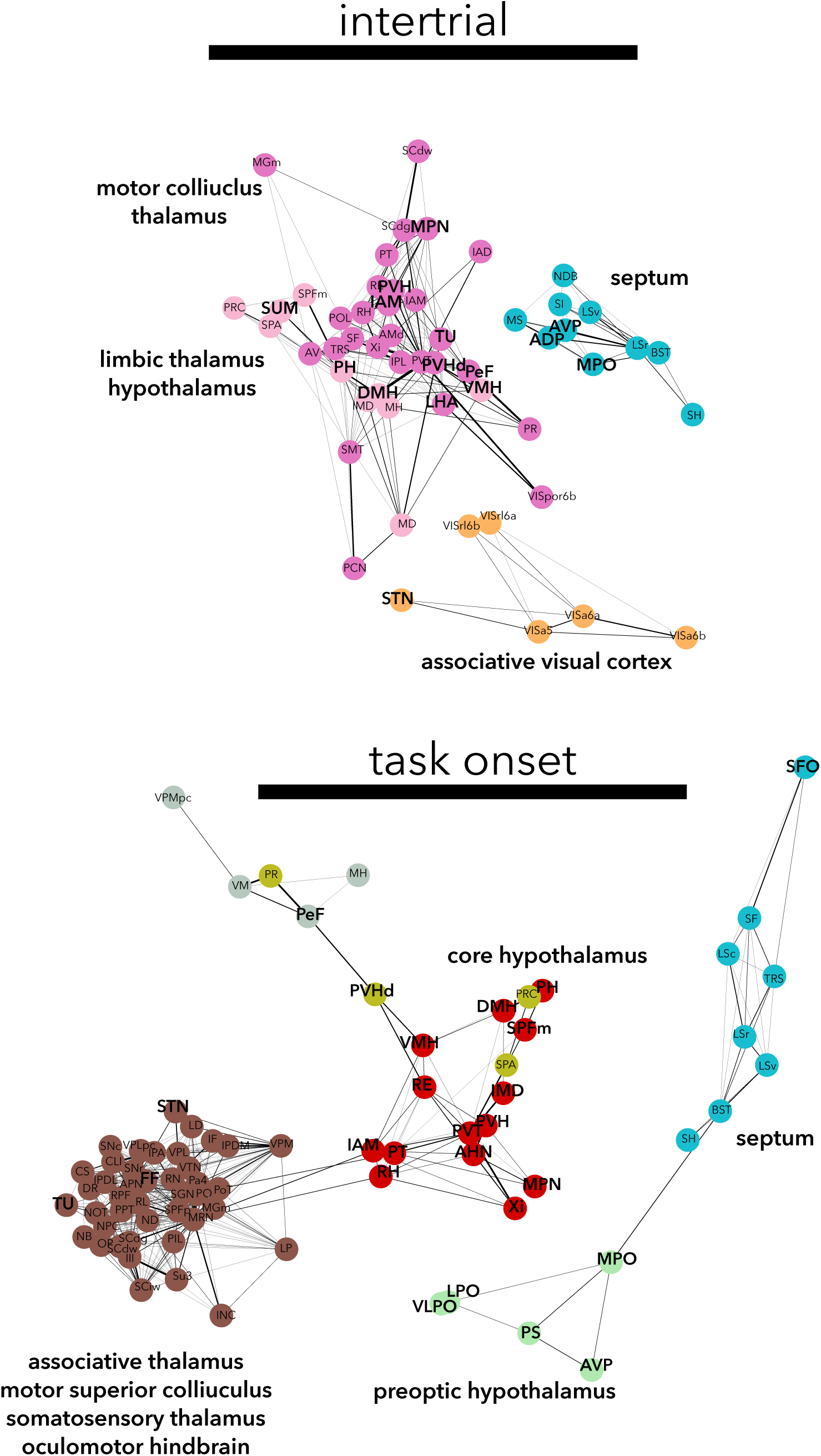
Electrospectral for the hypothalamus. Selection of communities containing areas that belong to the hypothalamus.

**Supplementary Figure 13:**
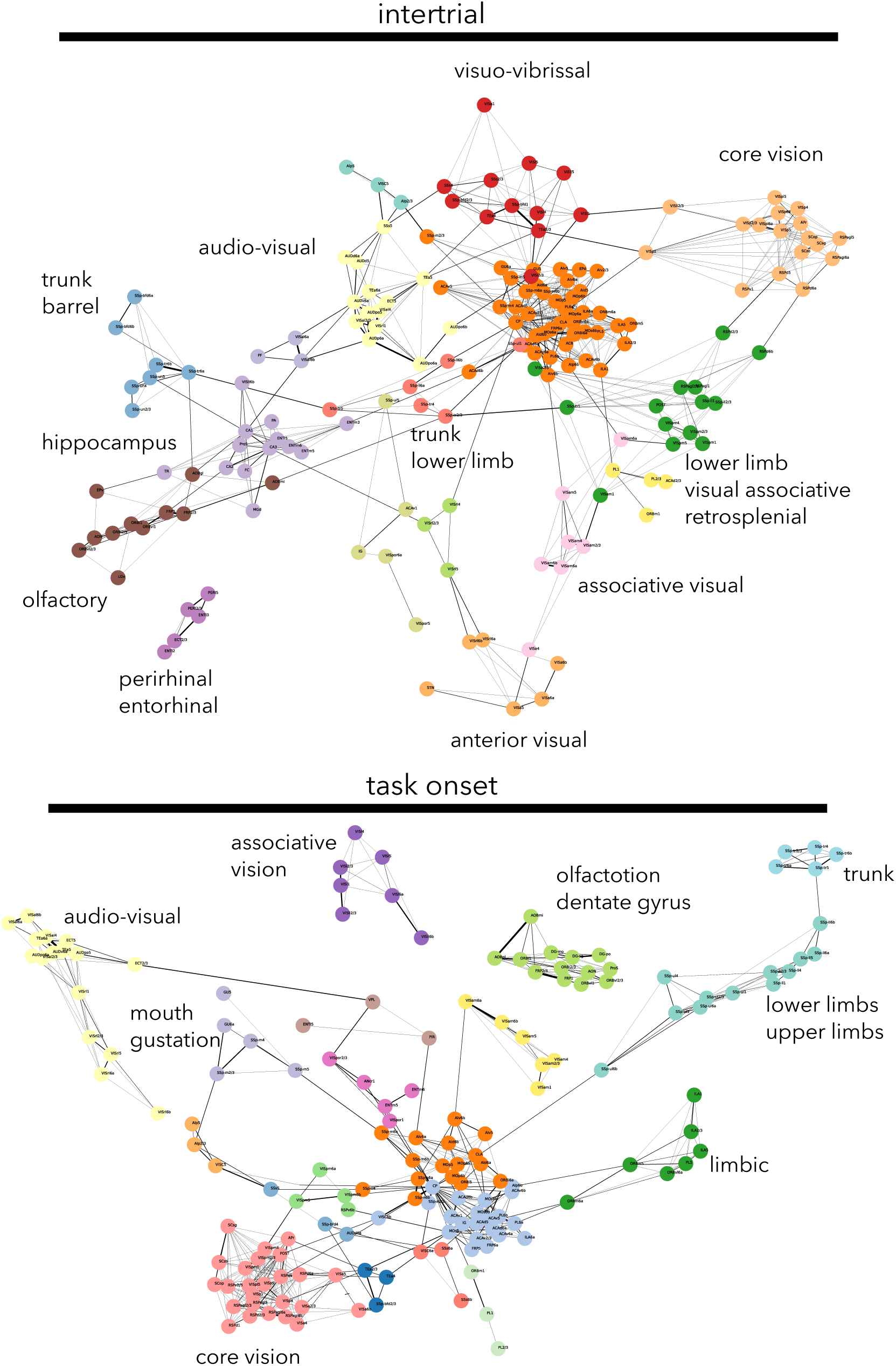
Electrospectral communities for cortical areas. Selection of communities containing areas that belong to the isocortex.

**Supplementary Figure 14:**
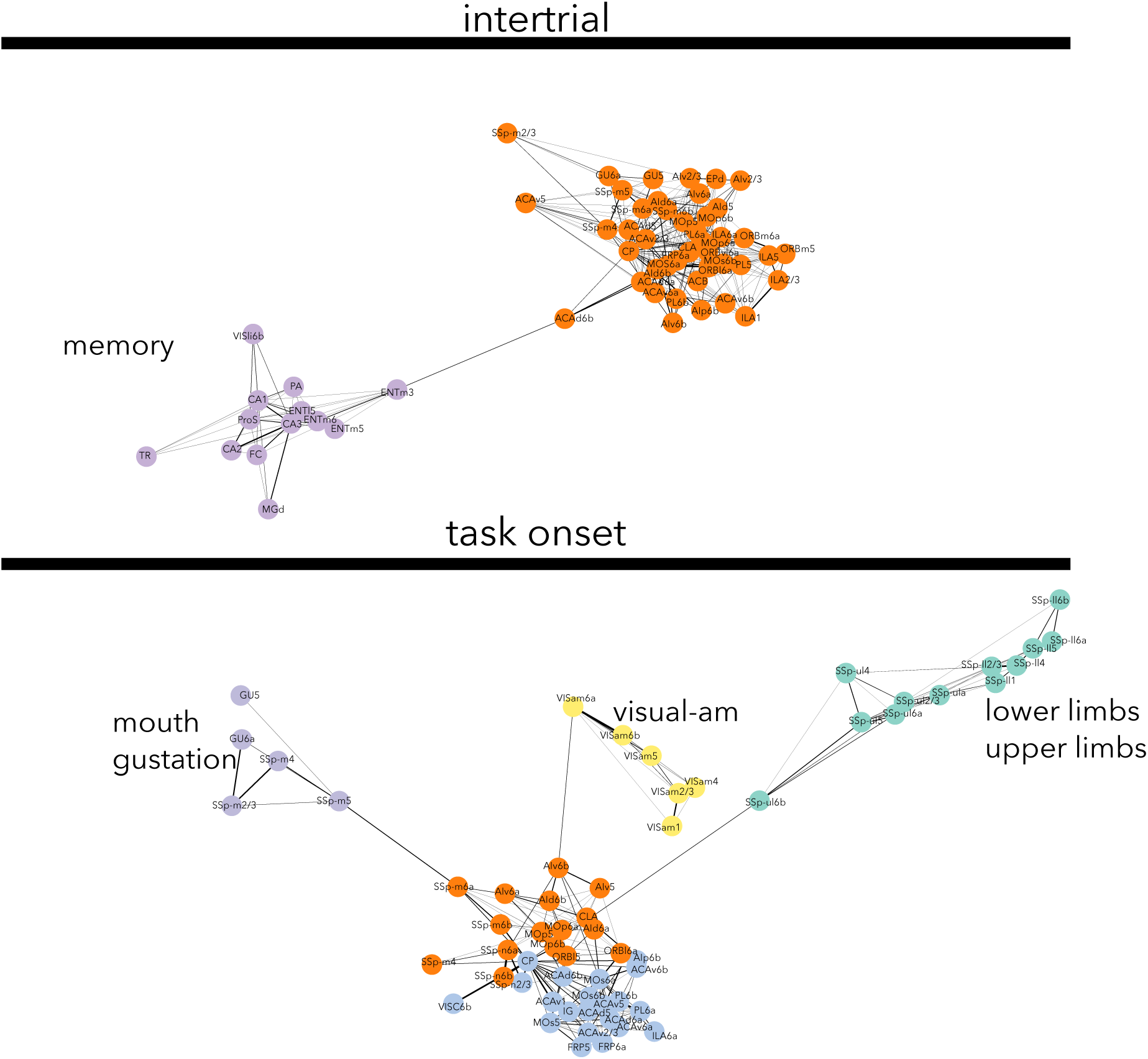
Selection intercommunity links with single connector nodes. Top: for intertrial communities, the memory community links to the motor planning community through layer 3 of the medial entorhinal cortex (ENTm3) and layer 6b of the anterior cingulate area (ACA6b). Bottom, for the task communities linking to the two motor planning communities, the gustatory community hinges on the layer 5 and 6 of mouth somatosensory cortex, visual-am through layer 6 of anteromedial visual cortex and agranular insular cortex layer 6b. The limbs community links through putamen and somatosensory upper limb layer 6b.

**Supplementary Figure 15:**
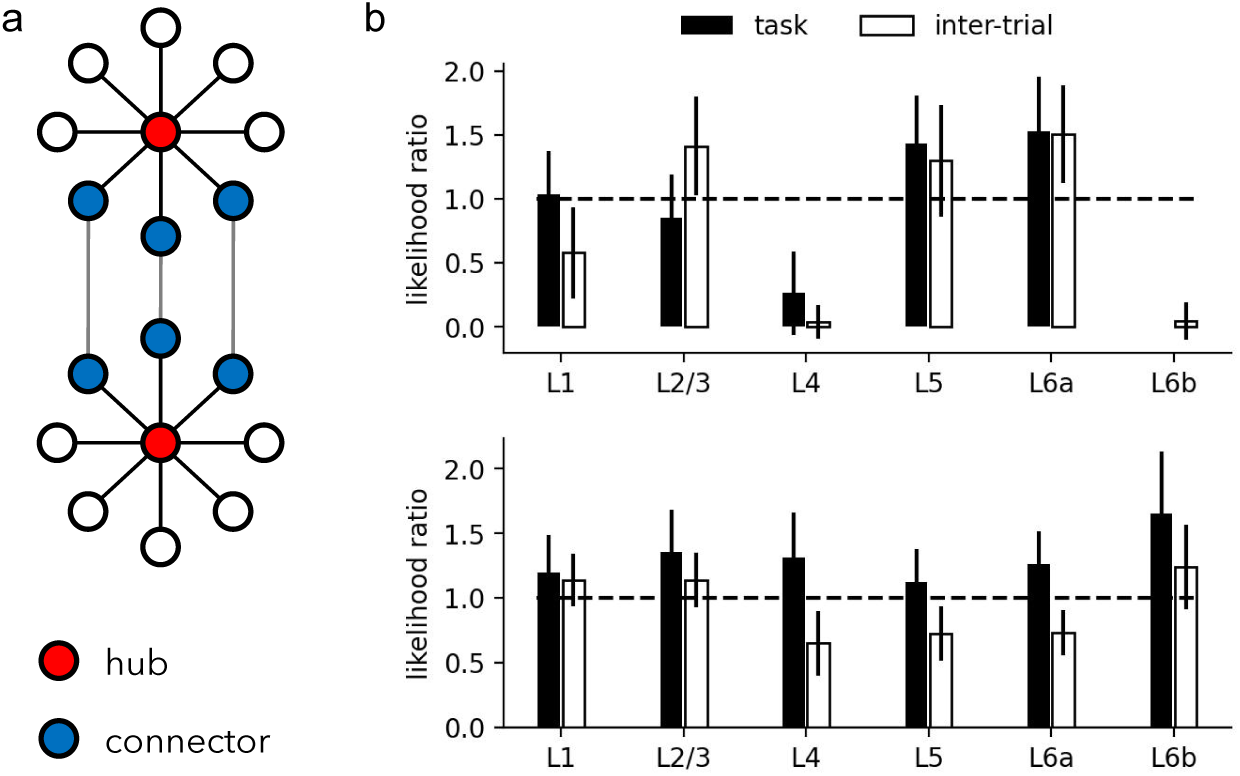
Layer specificity of hub and connector nodes. **a** Schematic definition of hub and connector nodes. **b** The ratio of probability (likelihood ratio) is defined as the bootstrapping estimate of the probability that a layer is attributed as a hub divided by the bootstrapping estimate of the probability that a layer is attributed to any community. **c** The likelihood ratio for a given layer to be a connector node. Errorbars represent standard deviation over 30 subsampled confusion matrices (Methods; Graph Theory Methods

**Supplementary Table 1:**
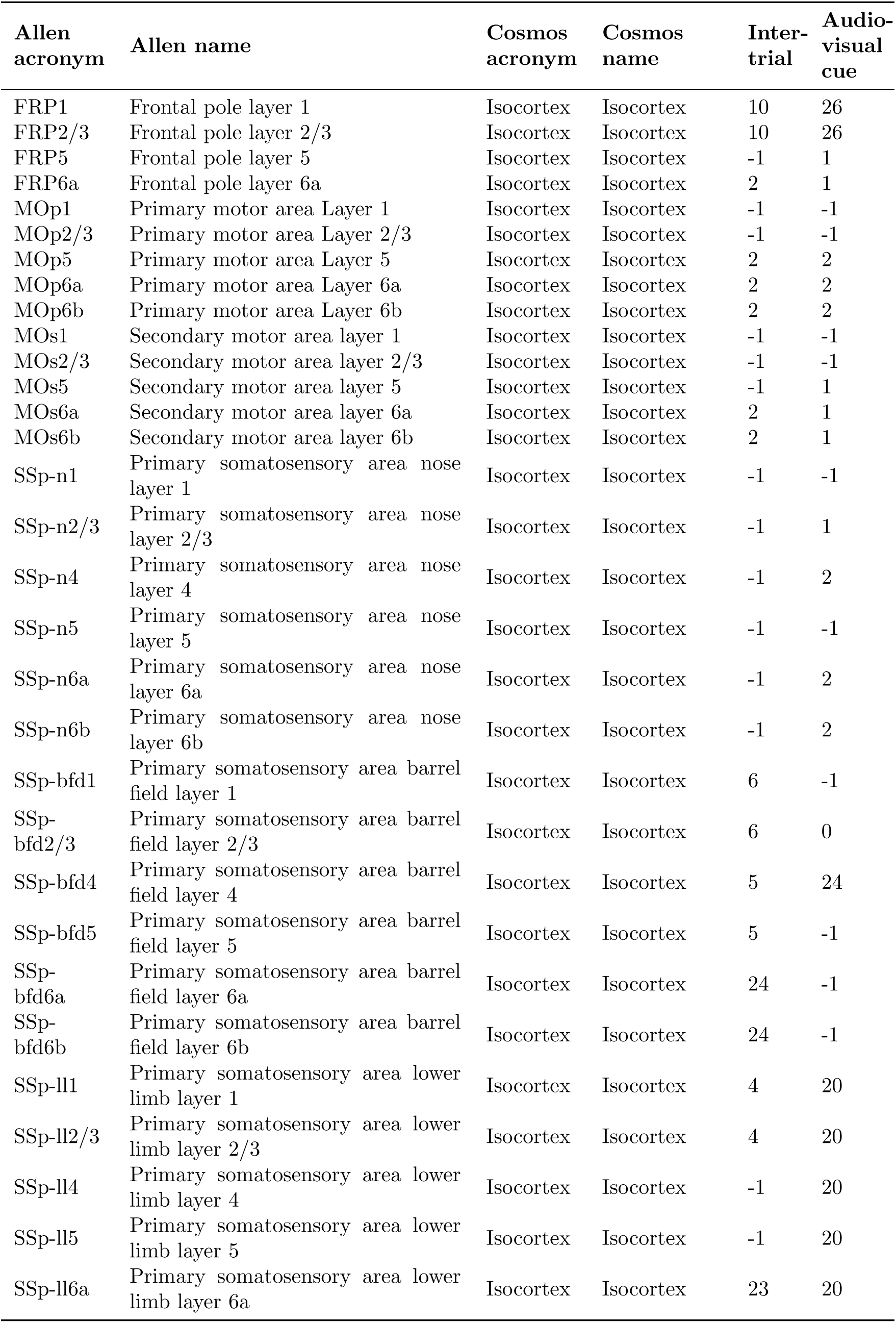
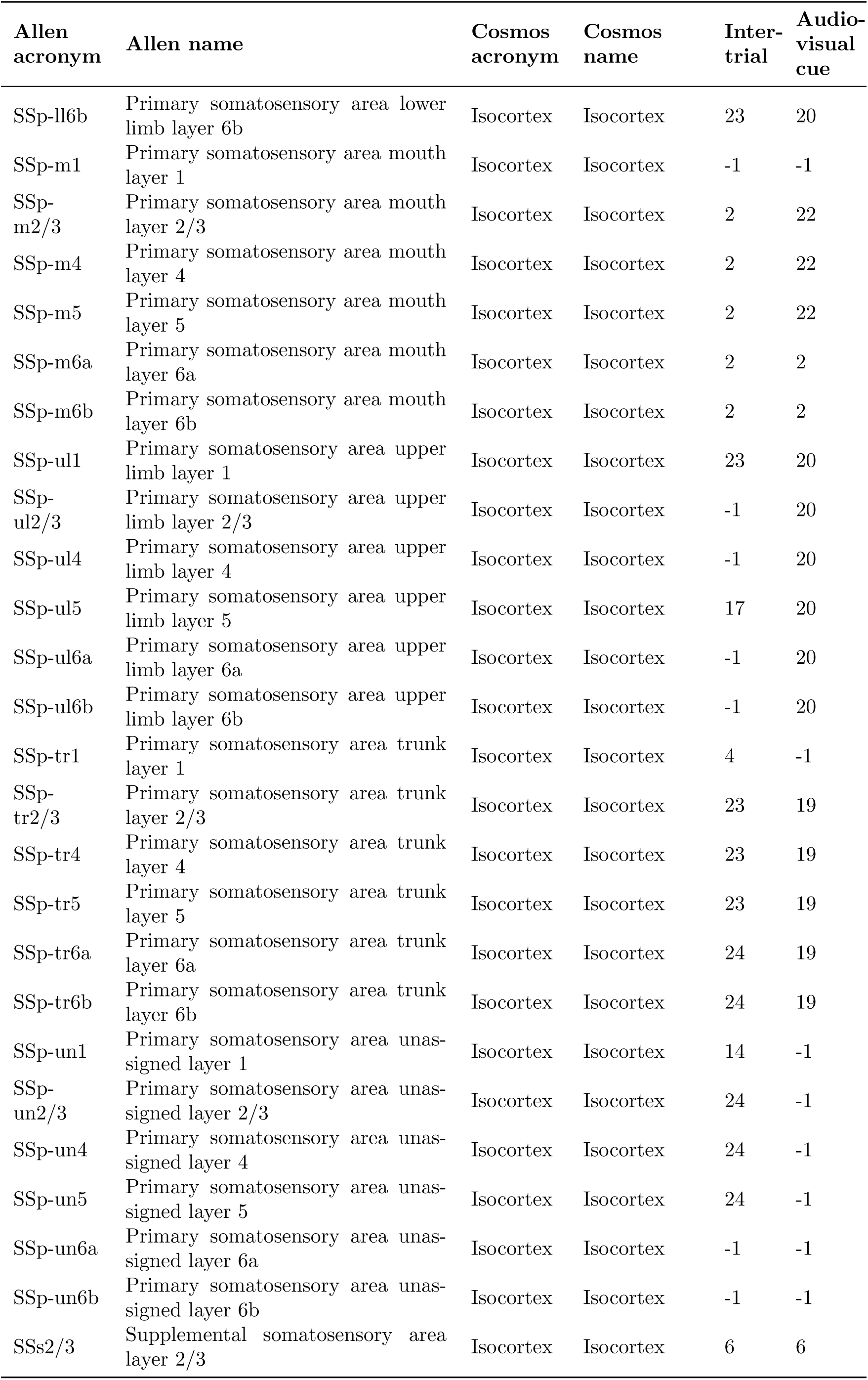
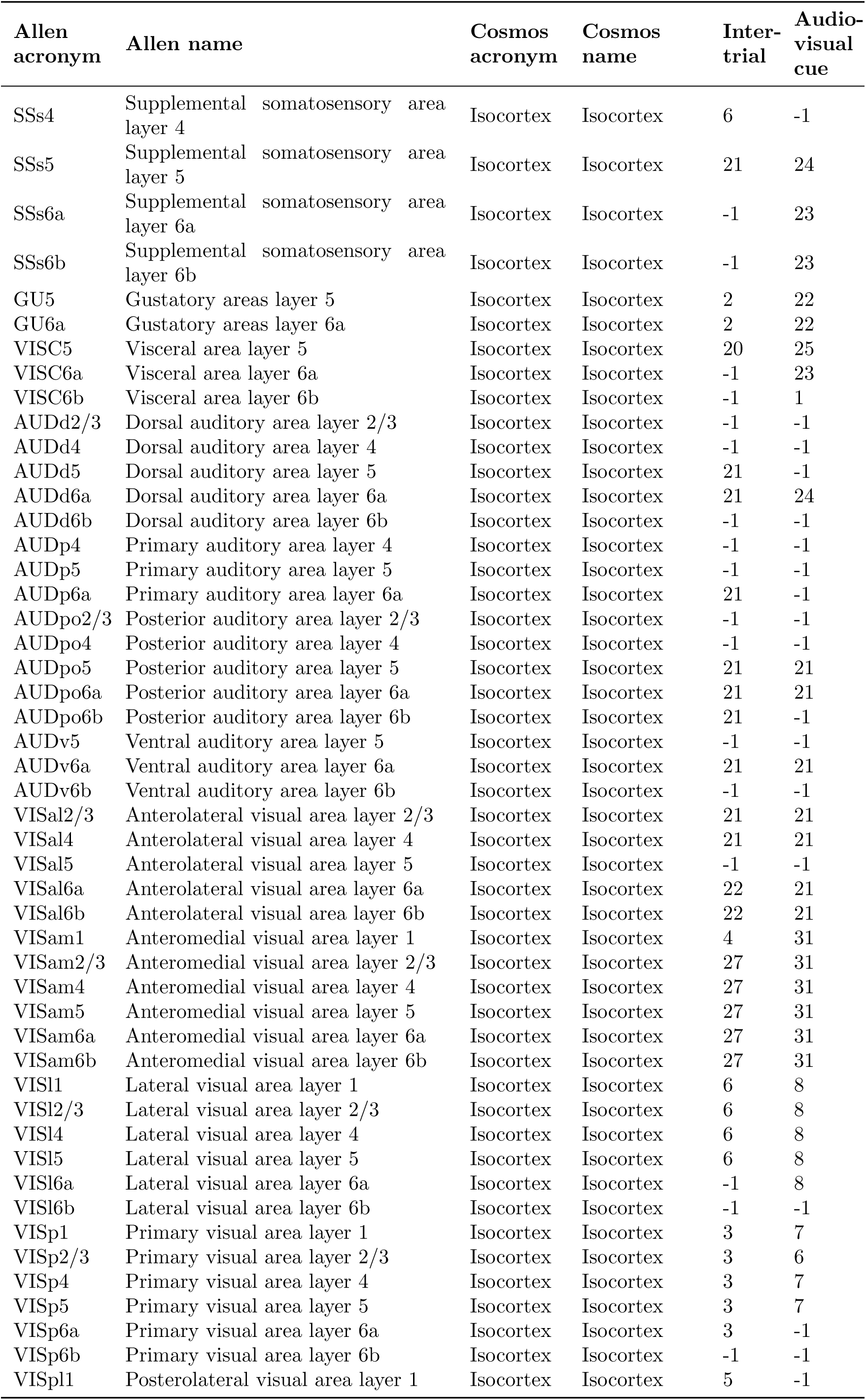
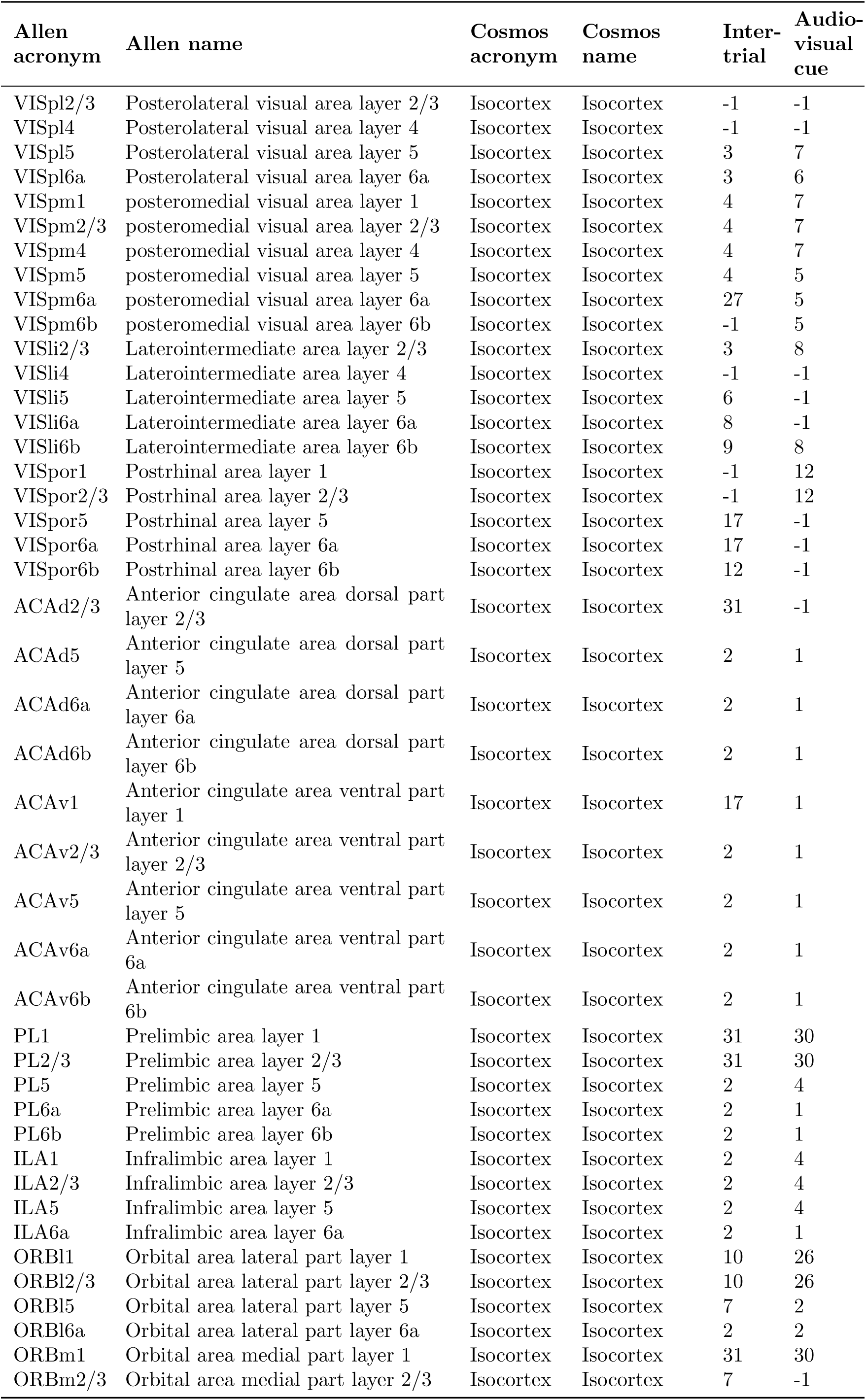
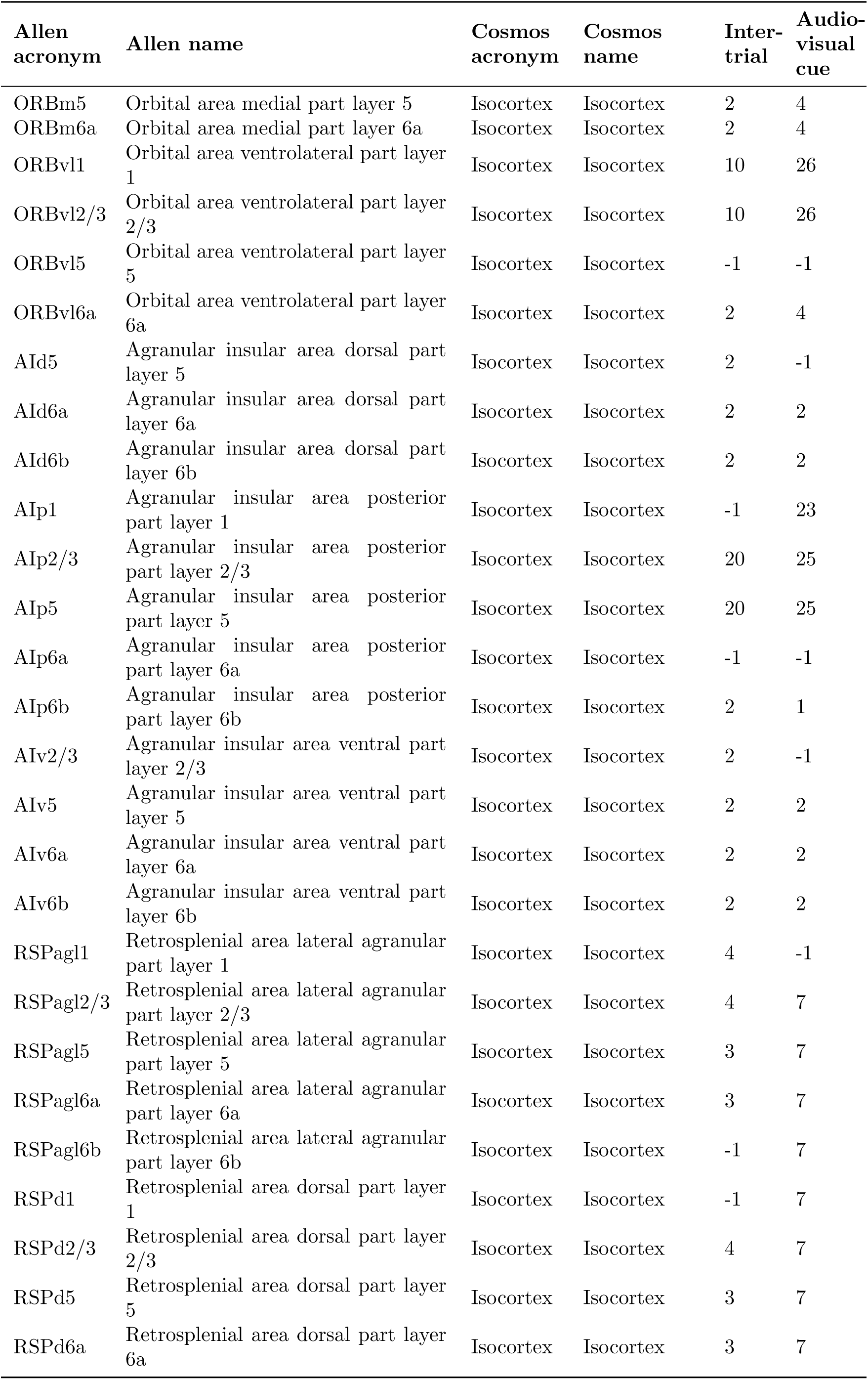
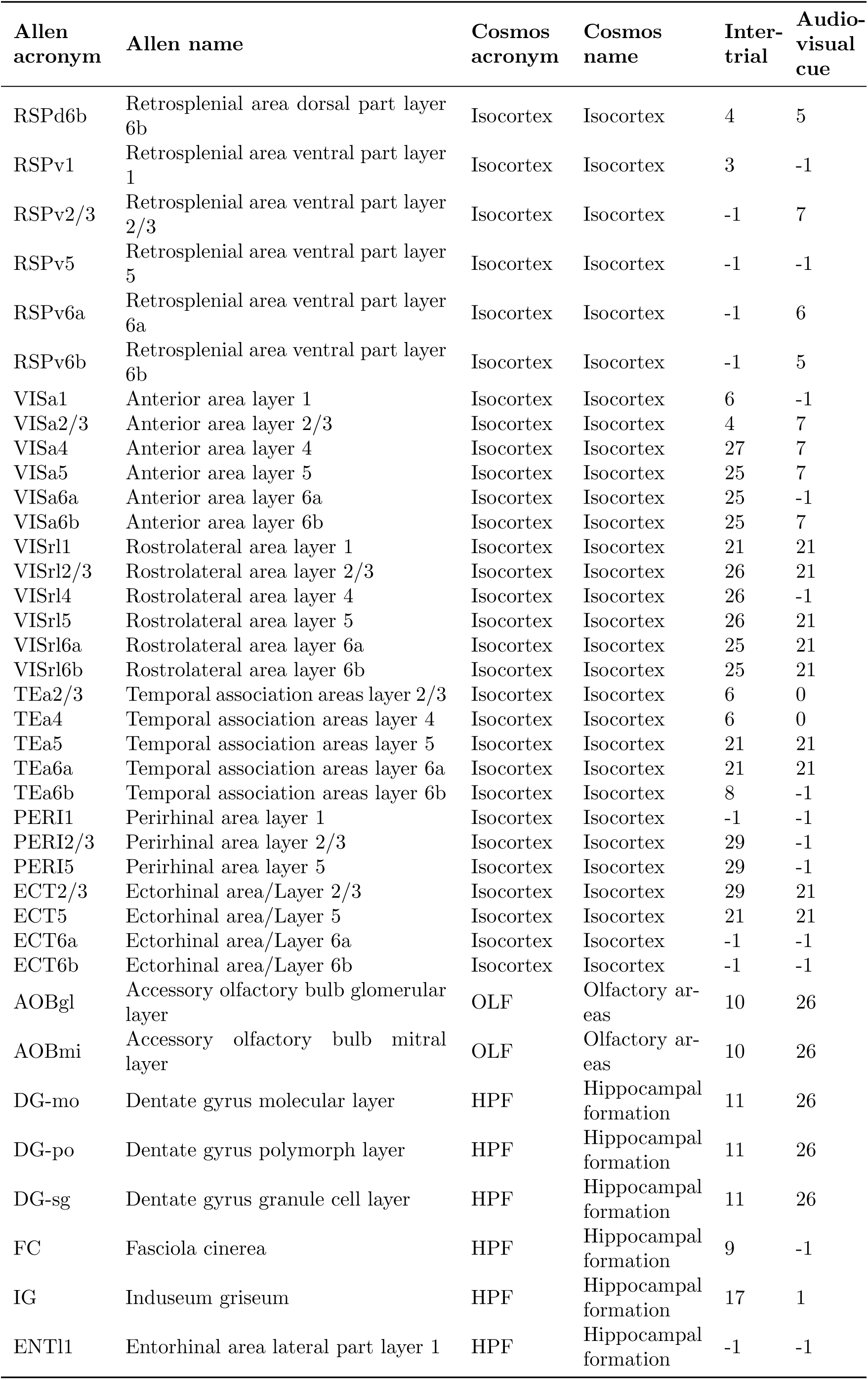
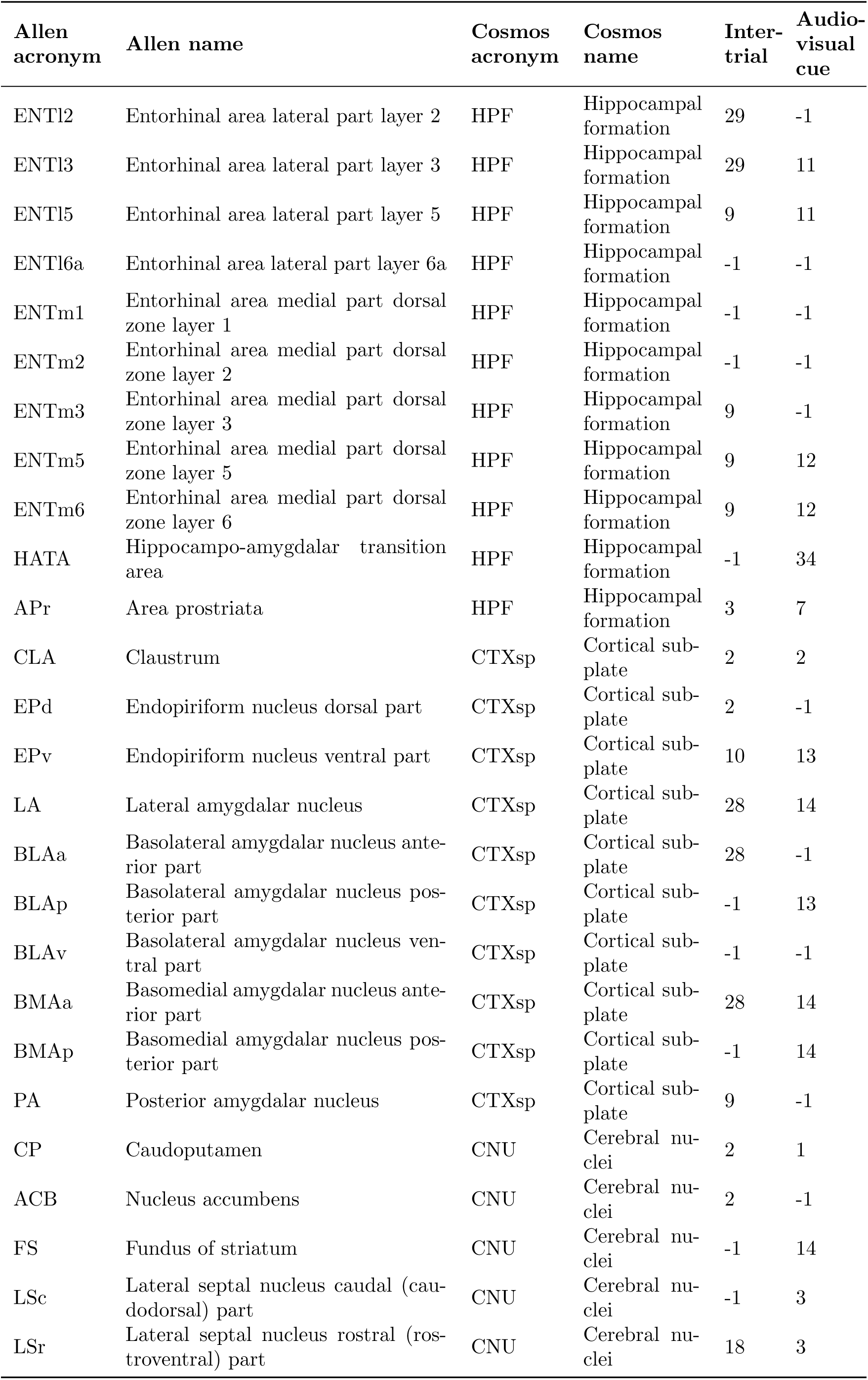
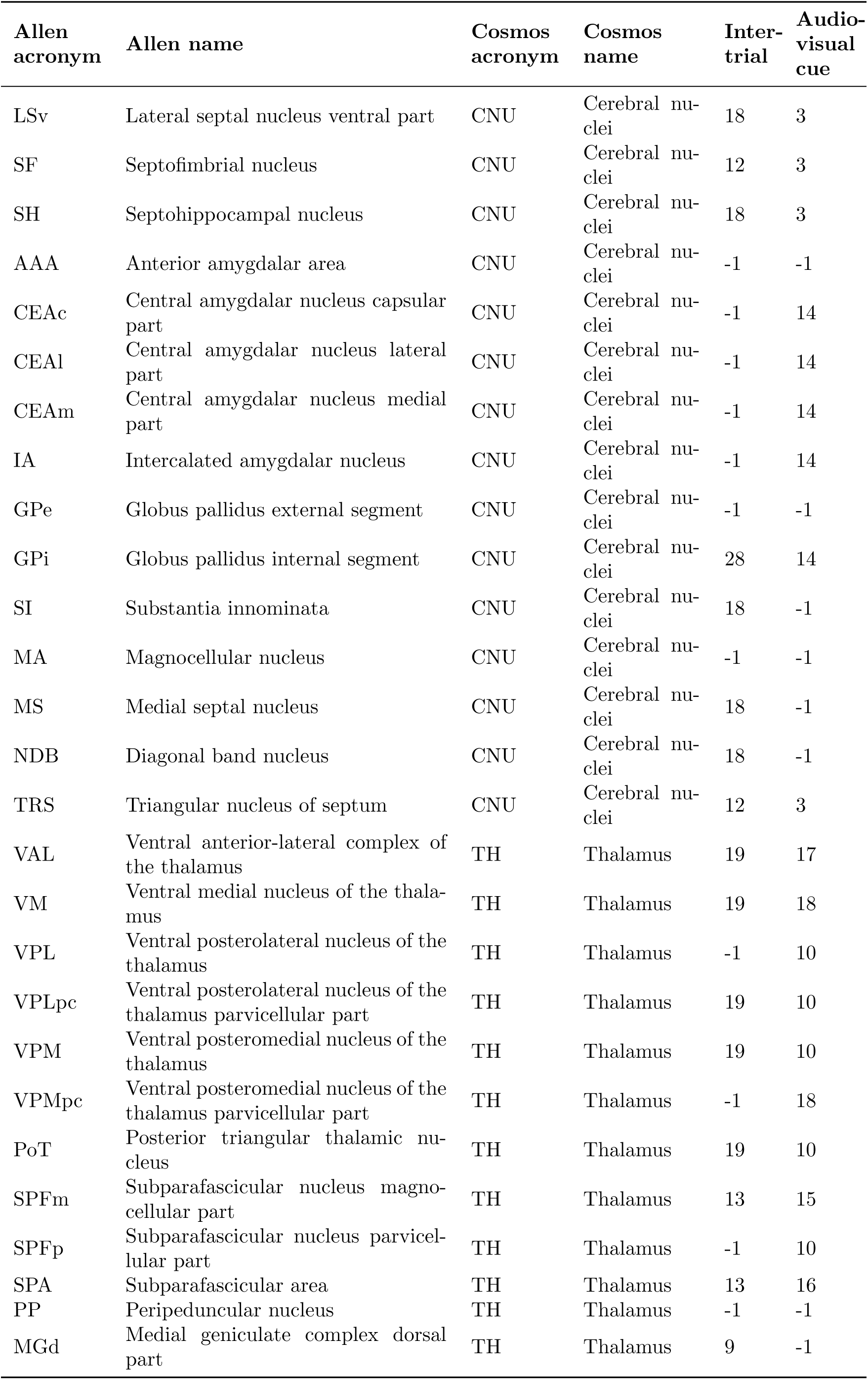
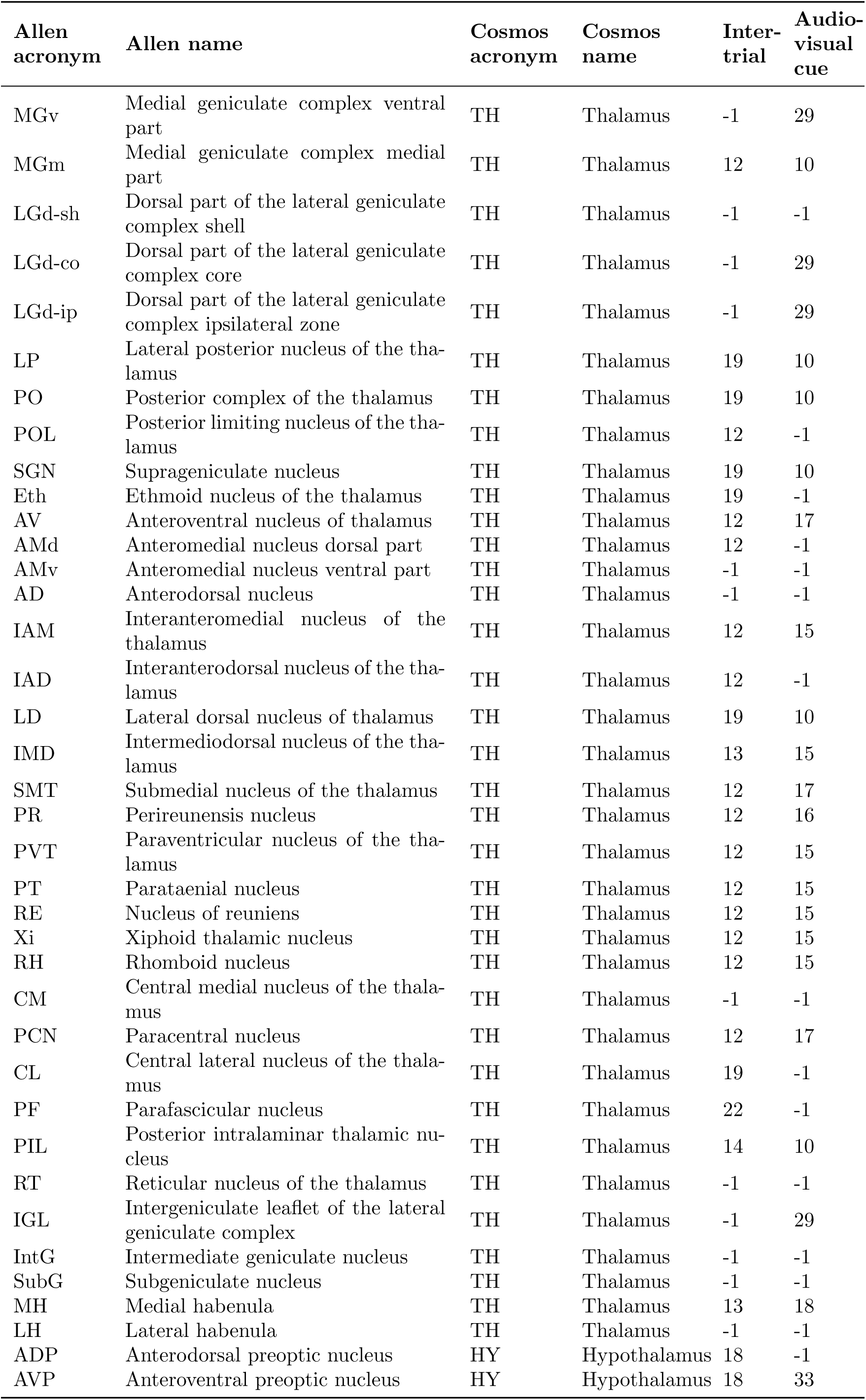
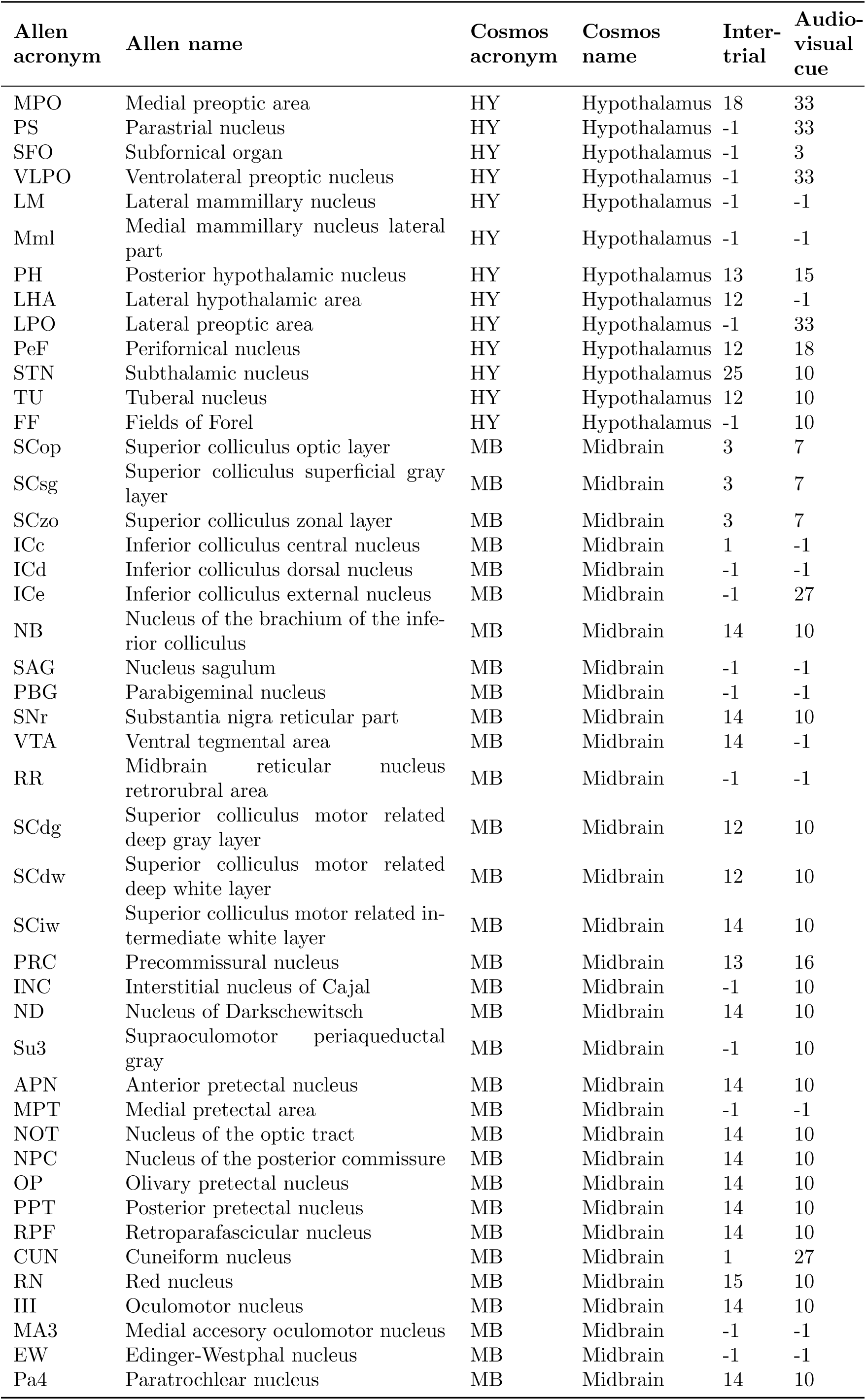
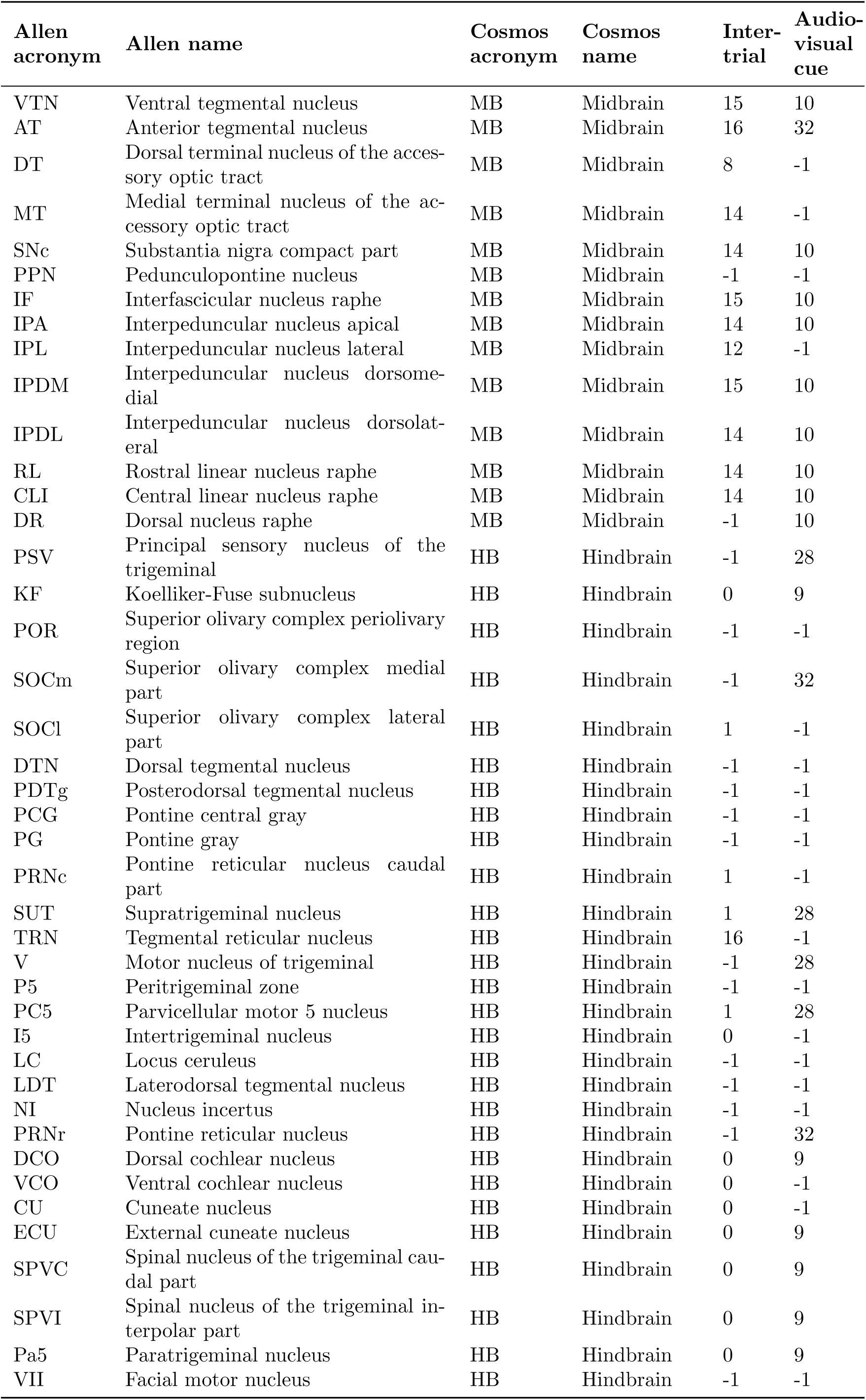
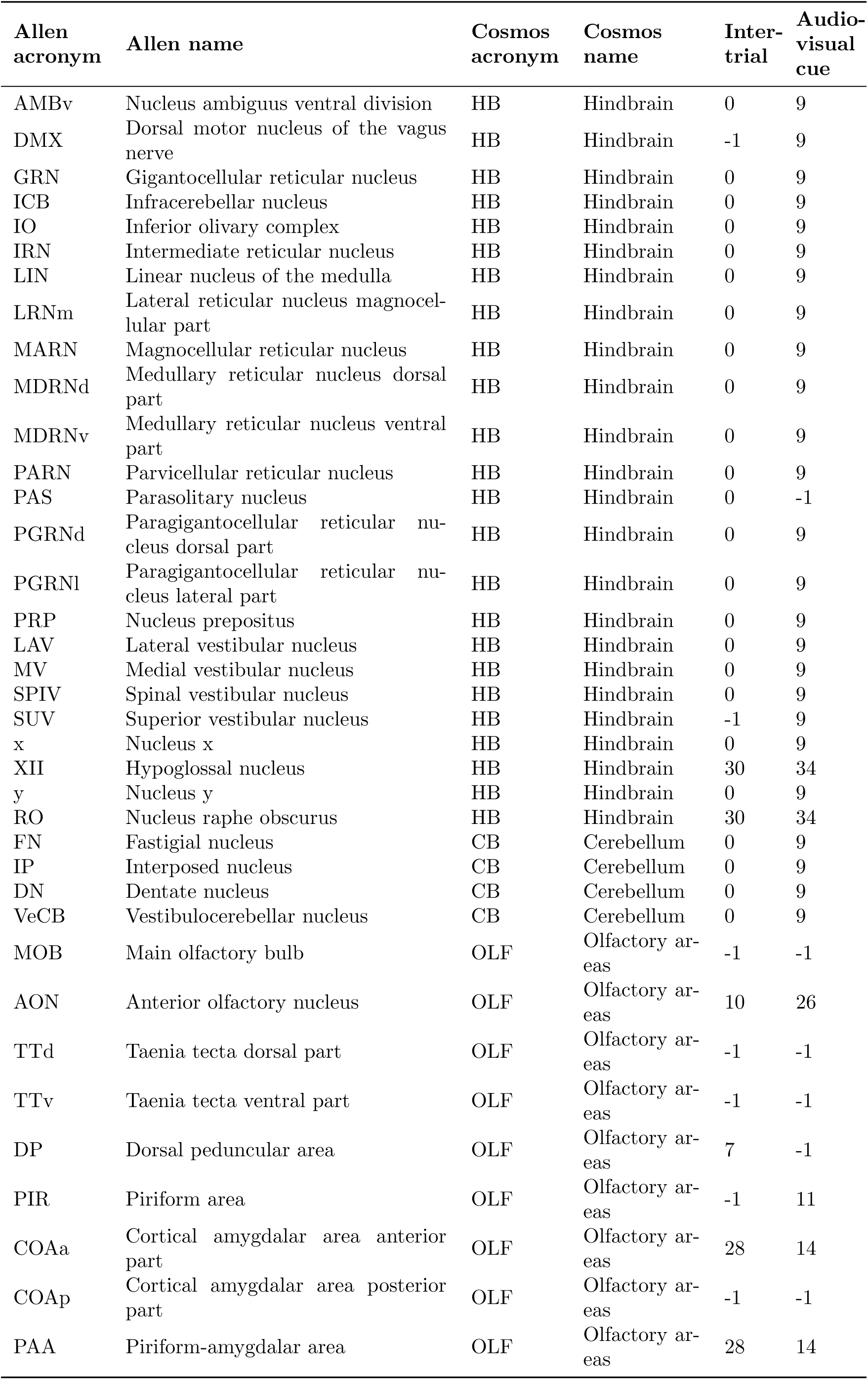
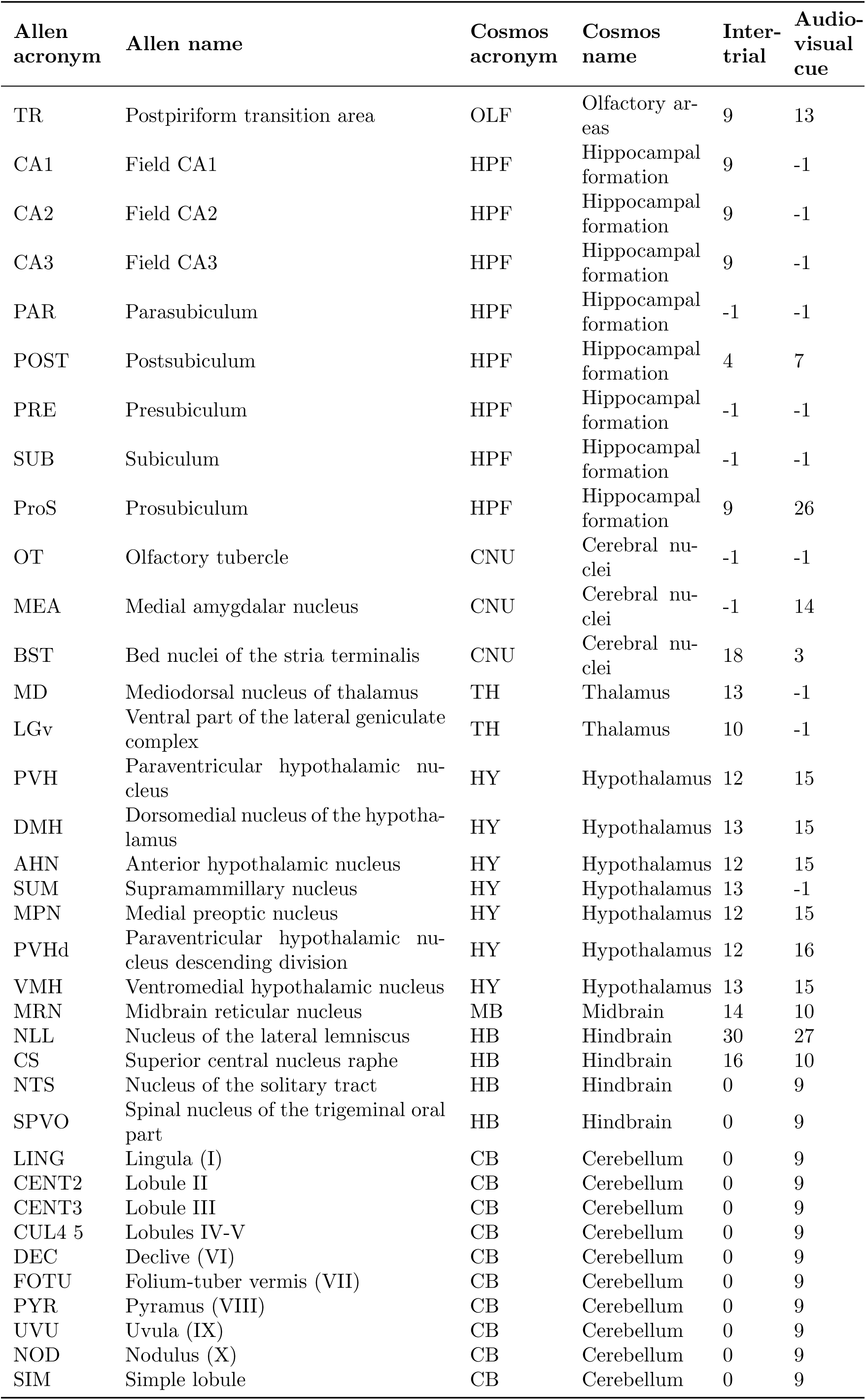
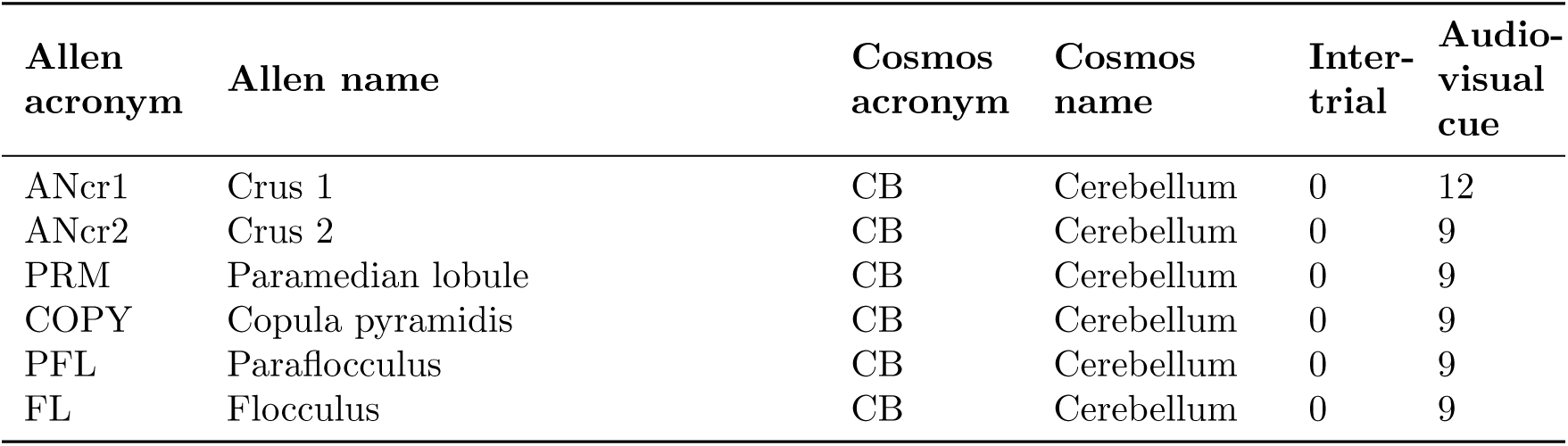
Table for community labels of each area.

**Supplementary Table 3:**
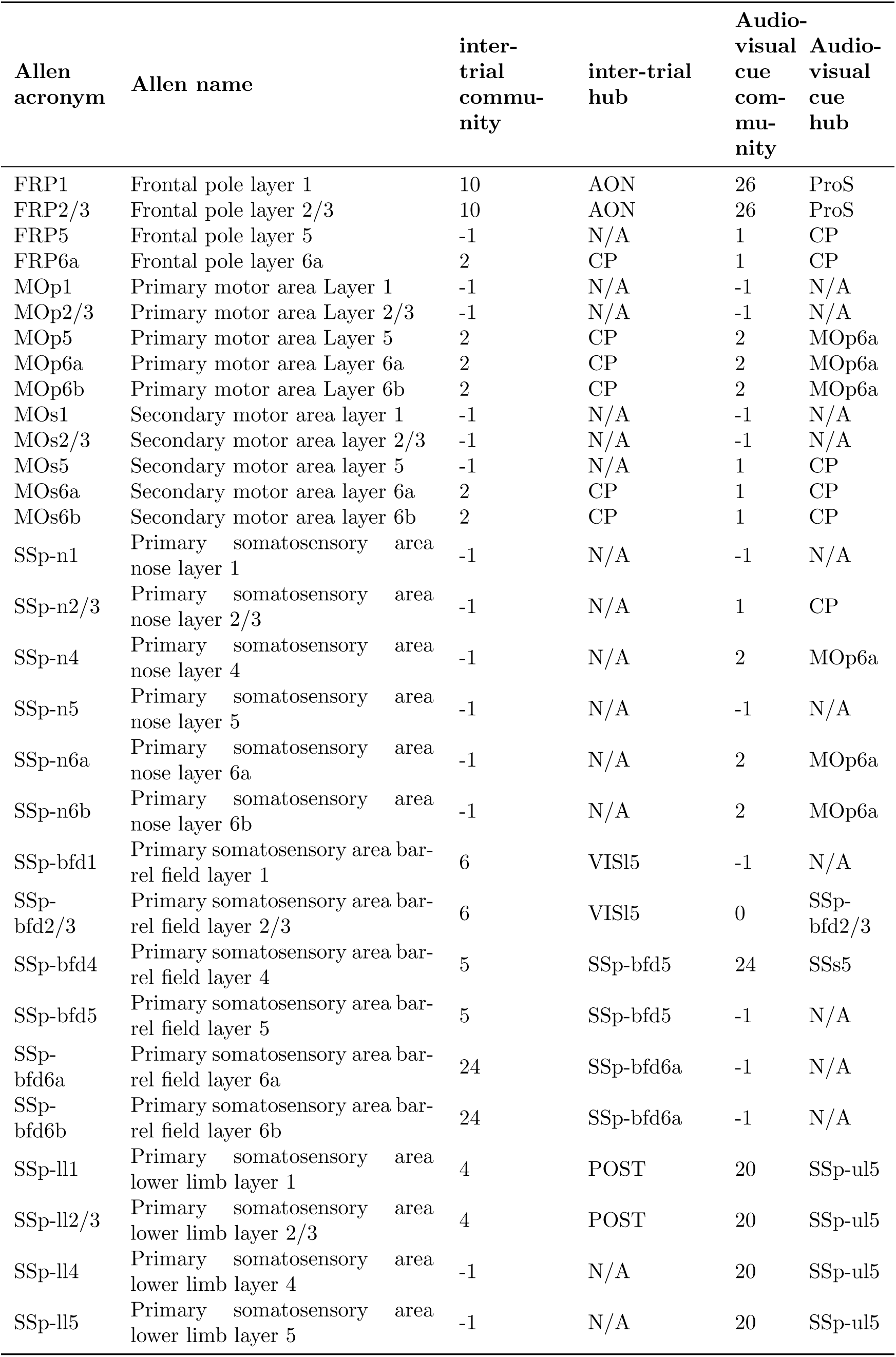
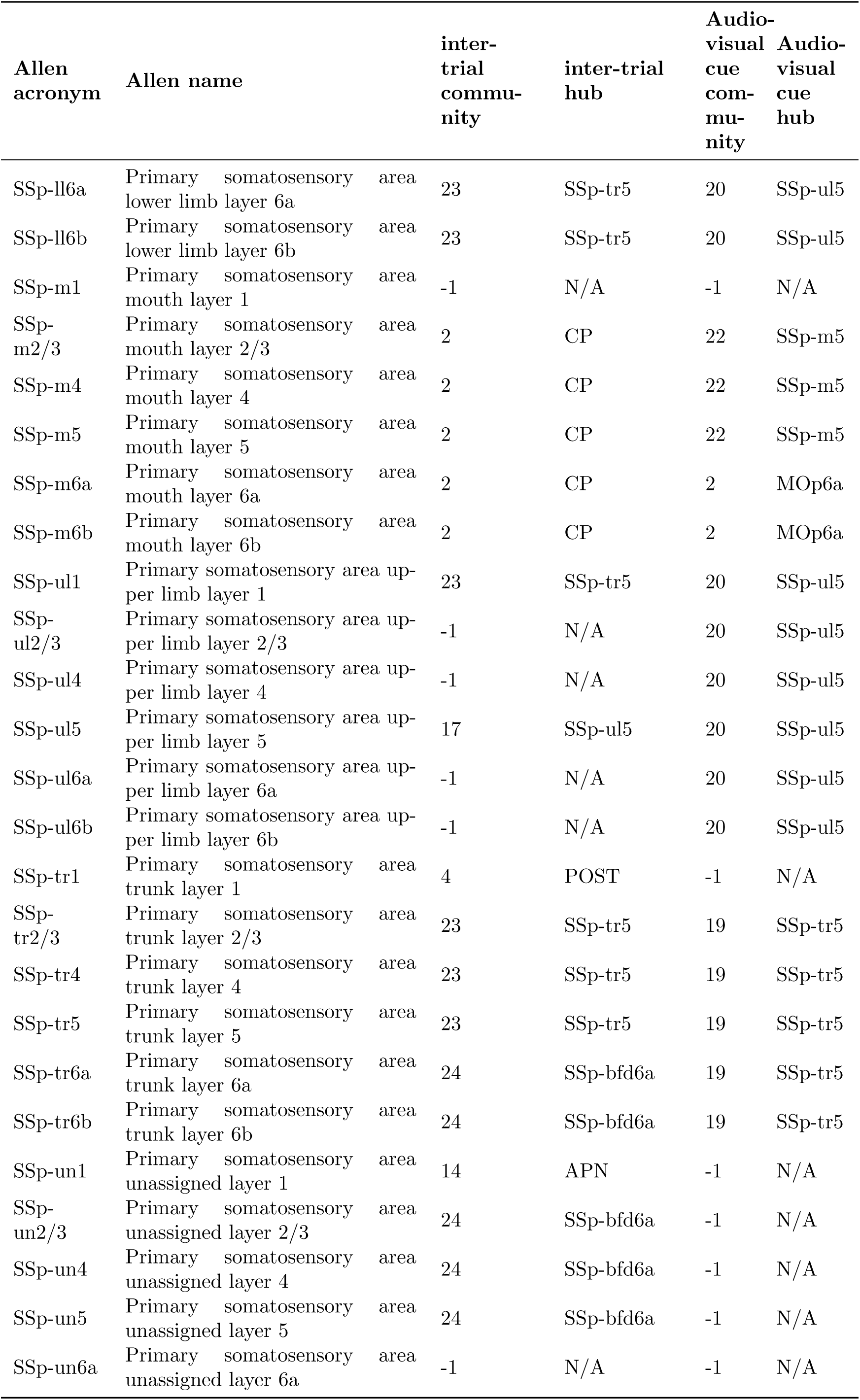
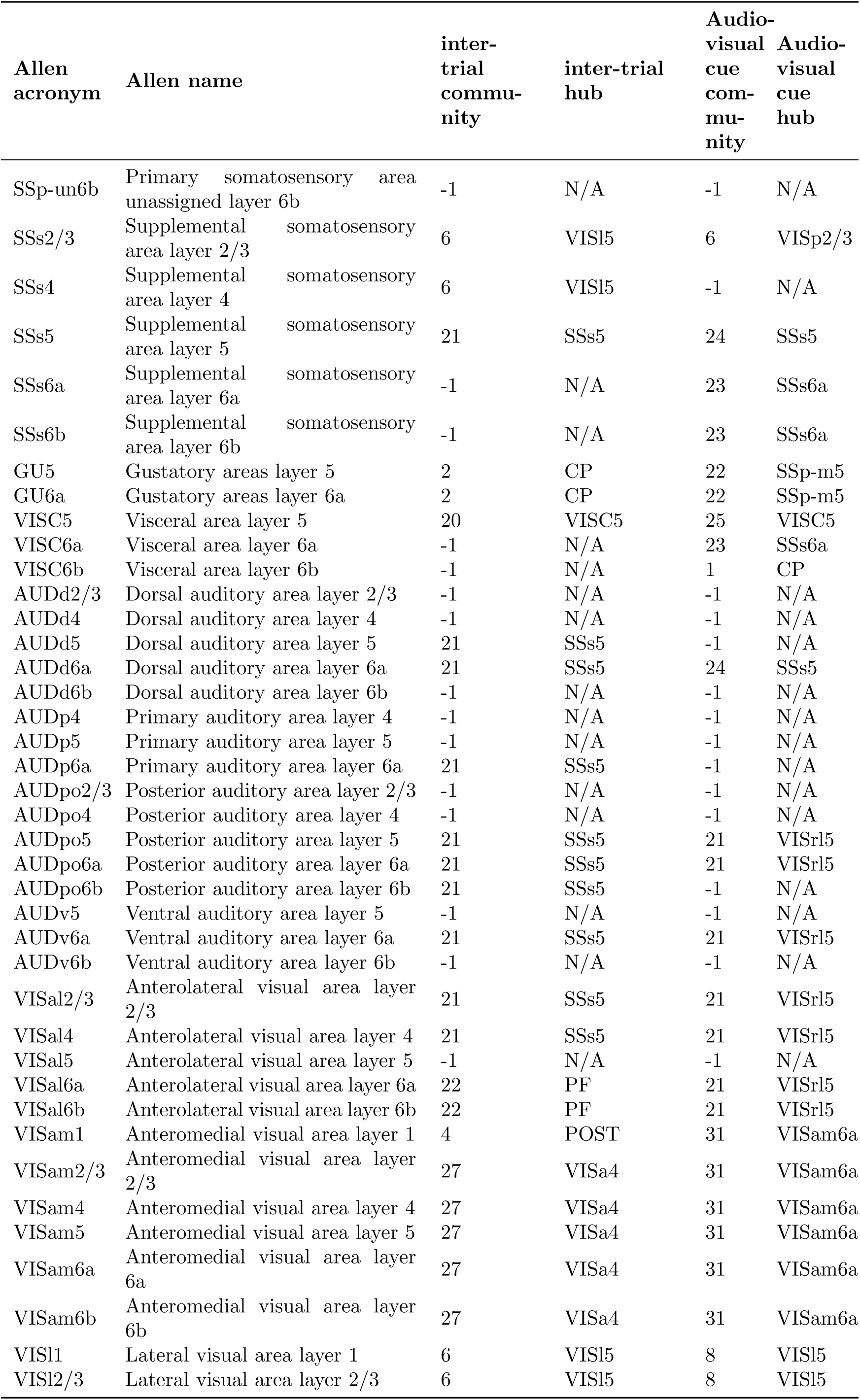
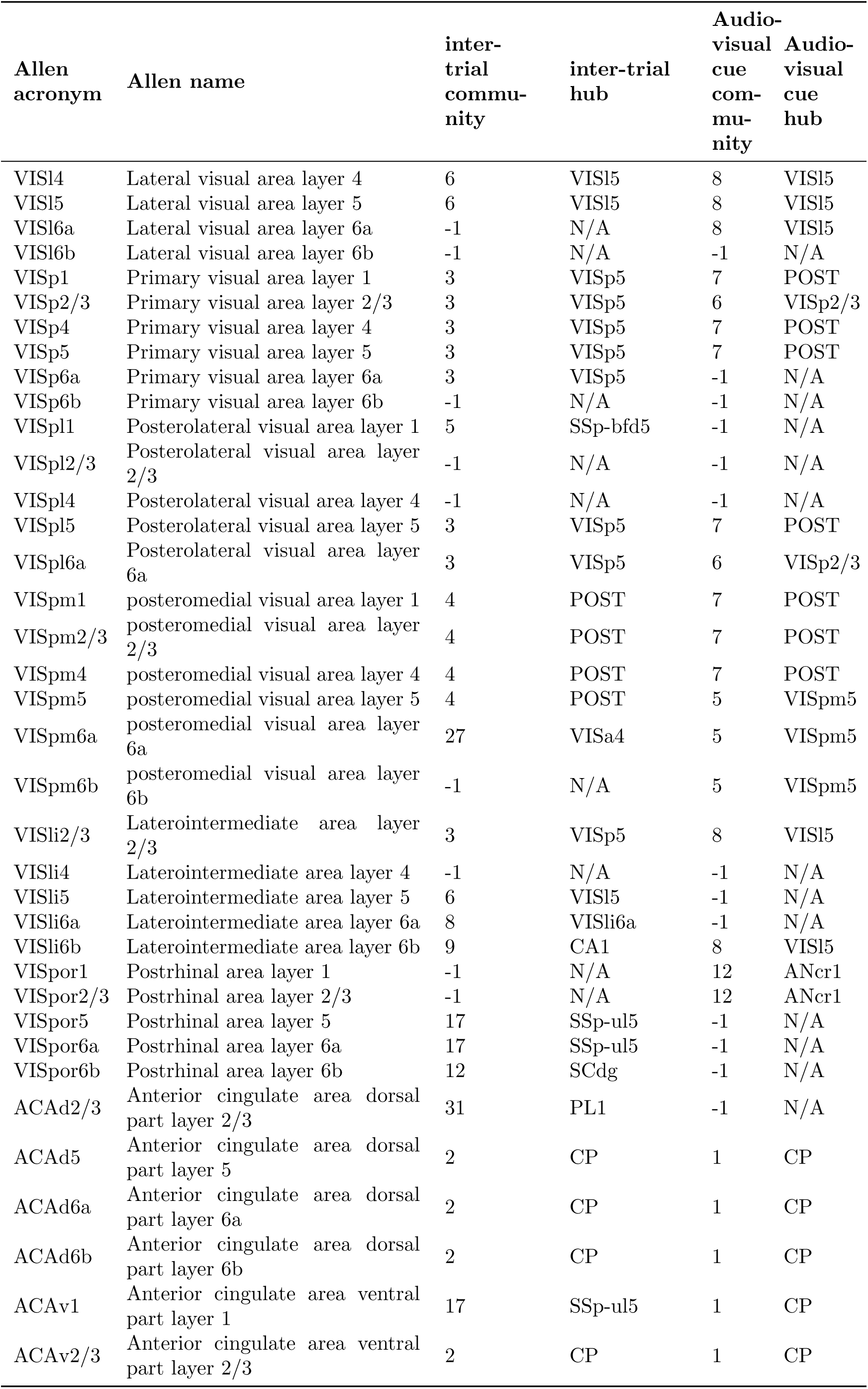
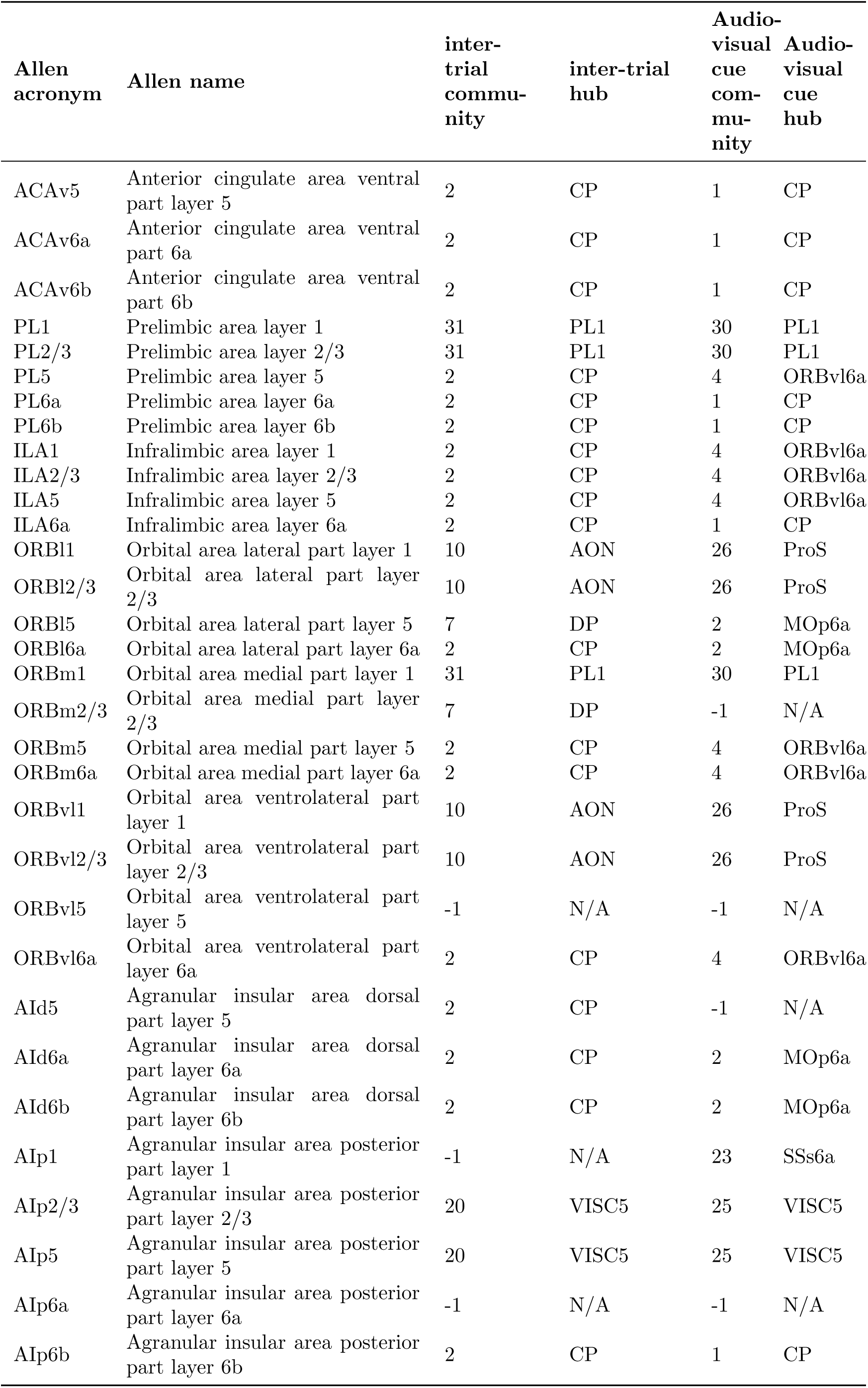
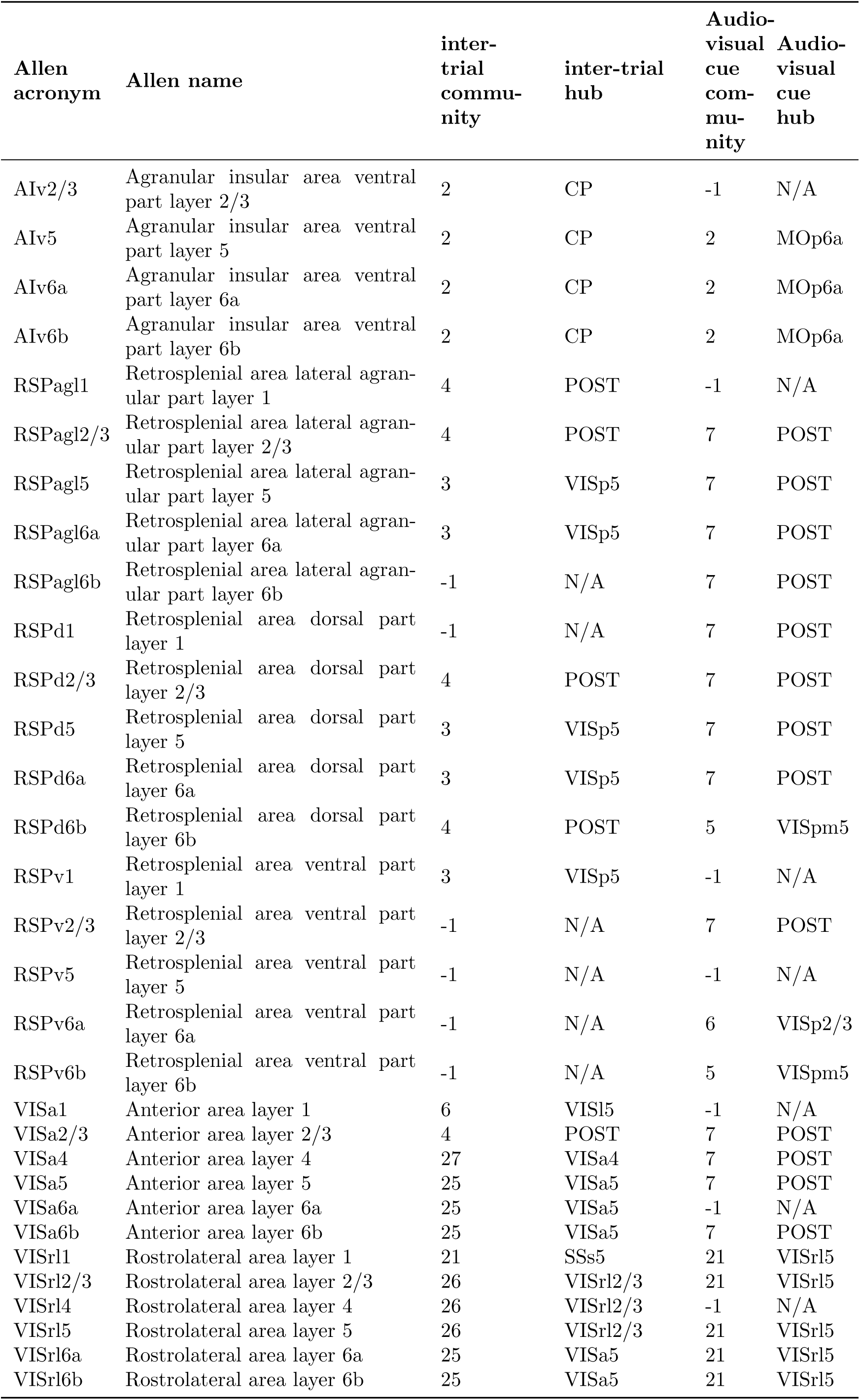
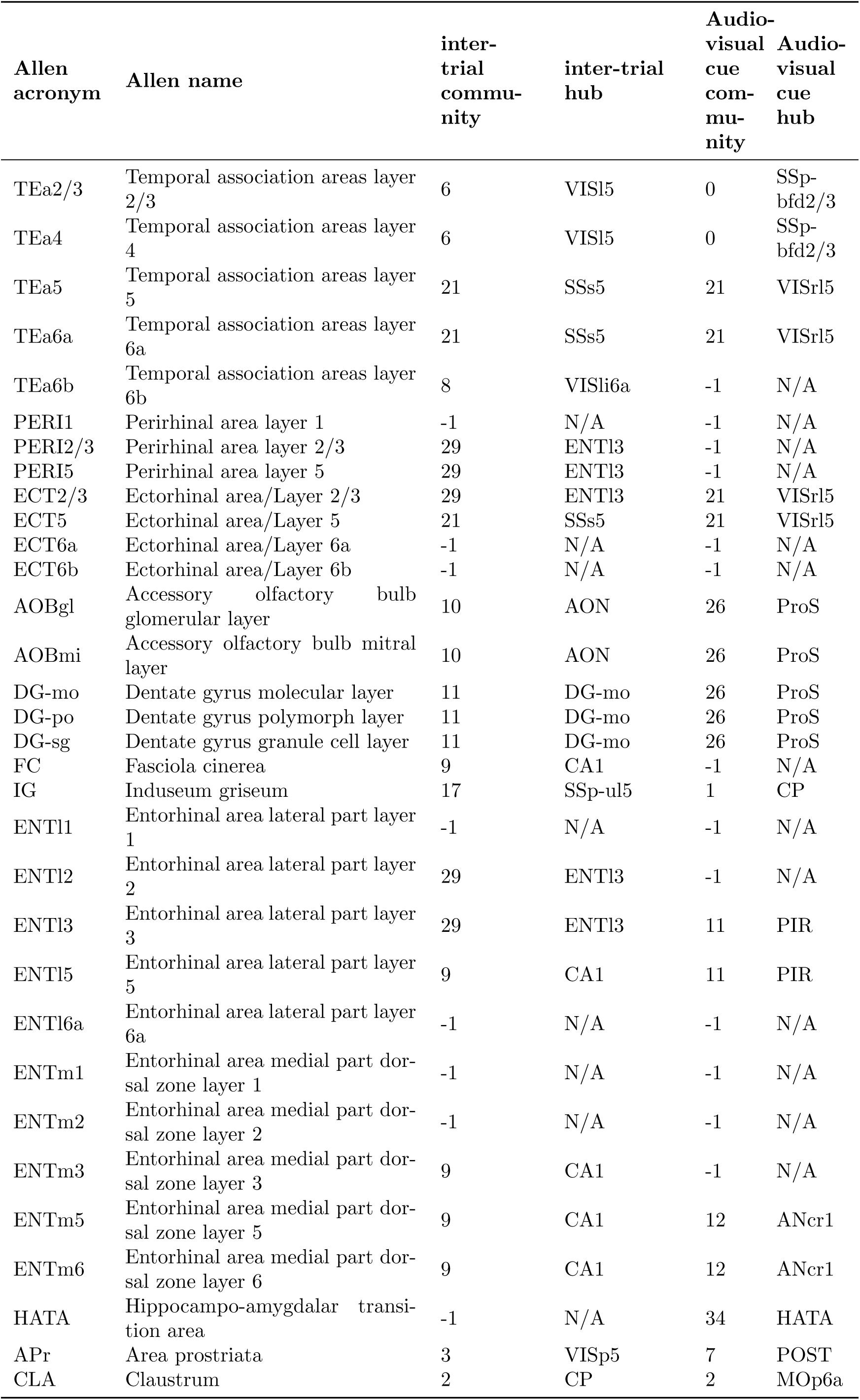
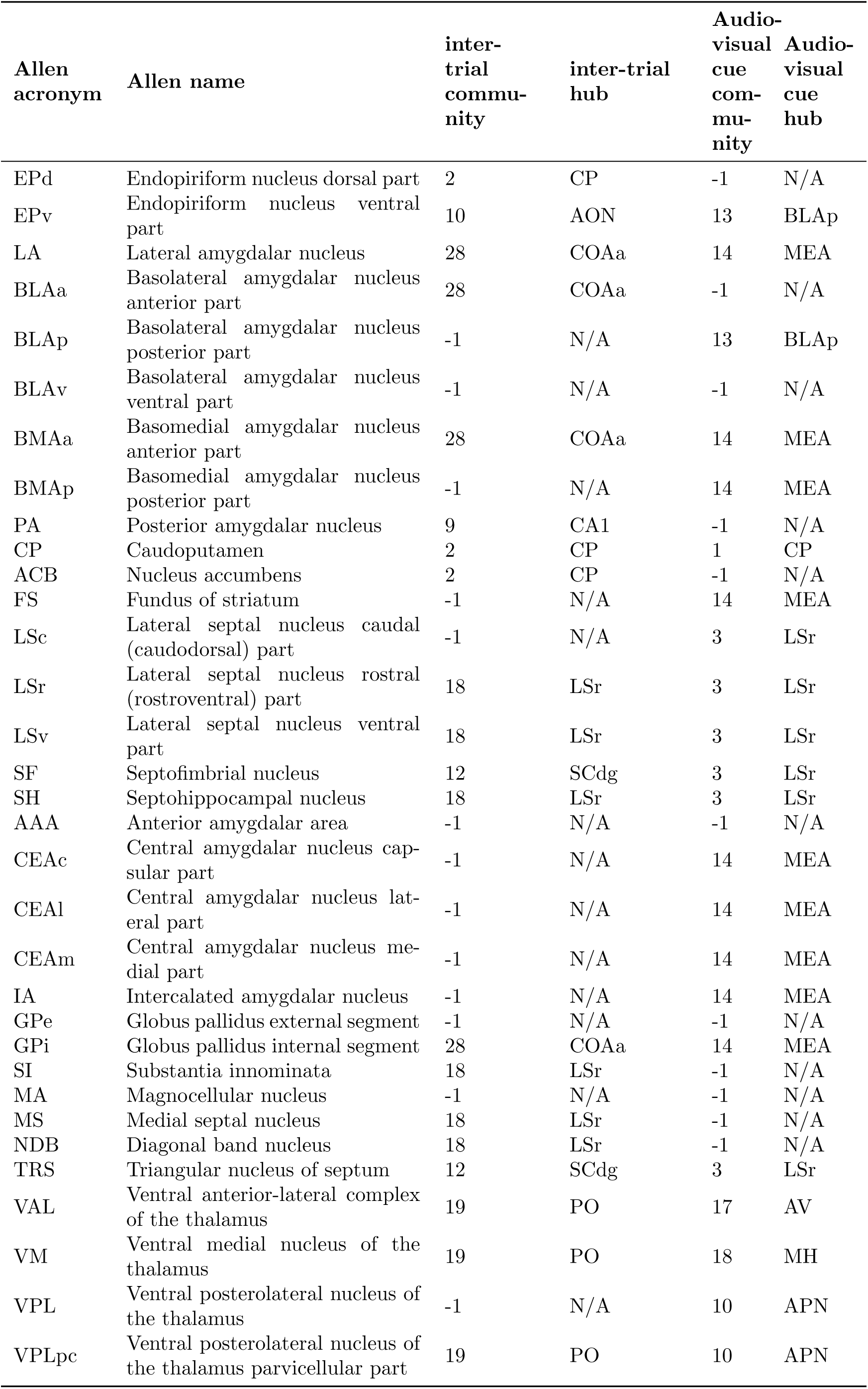
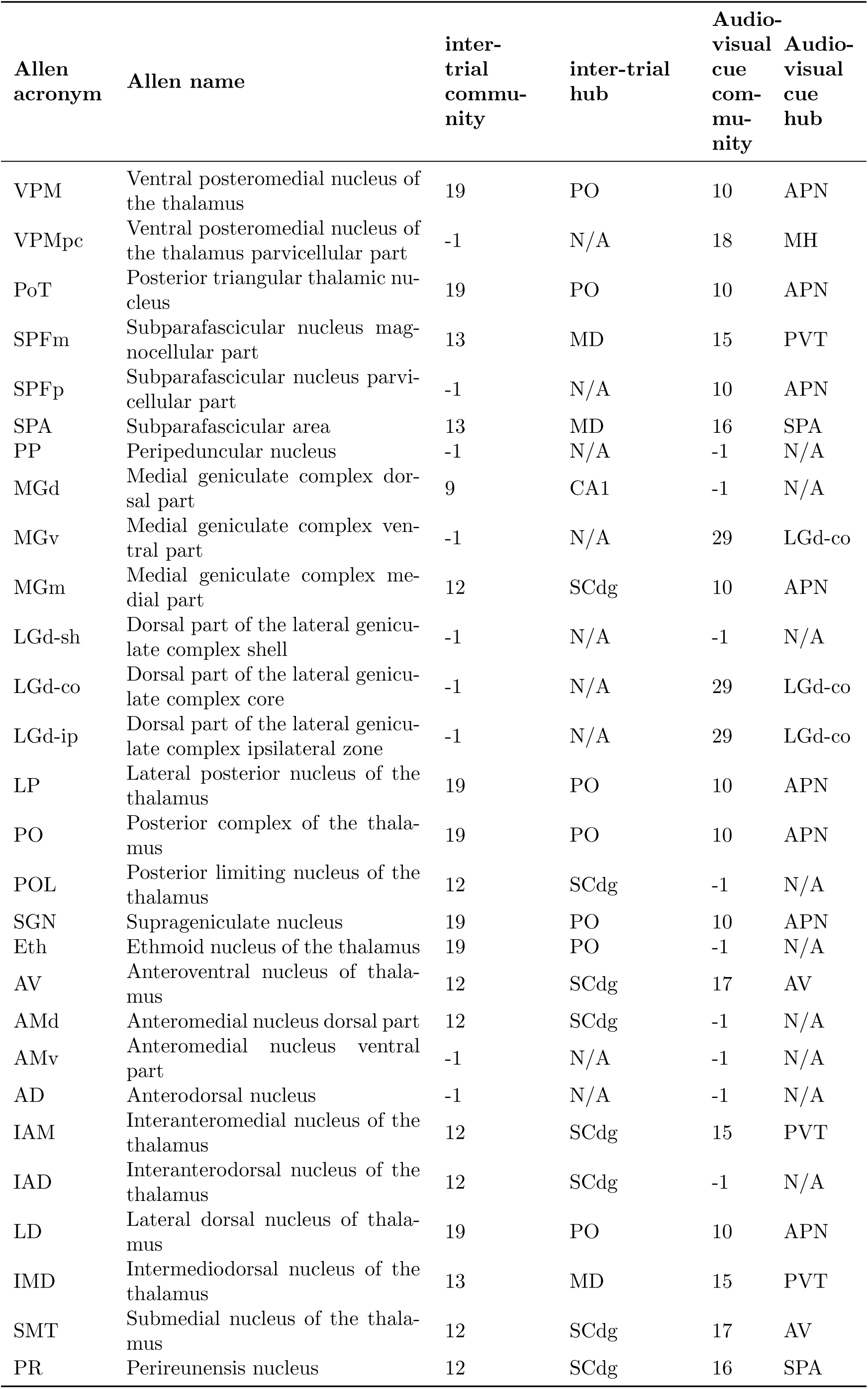
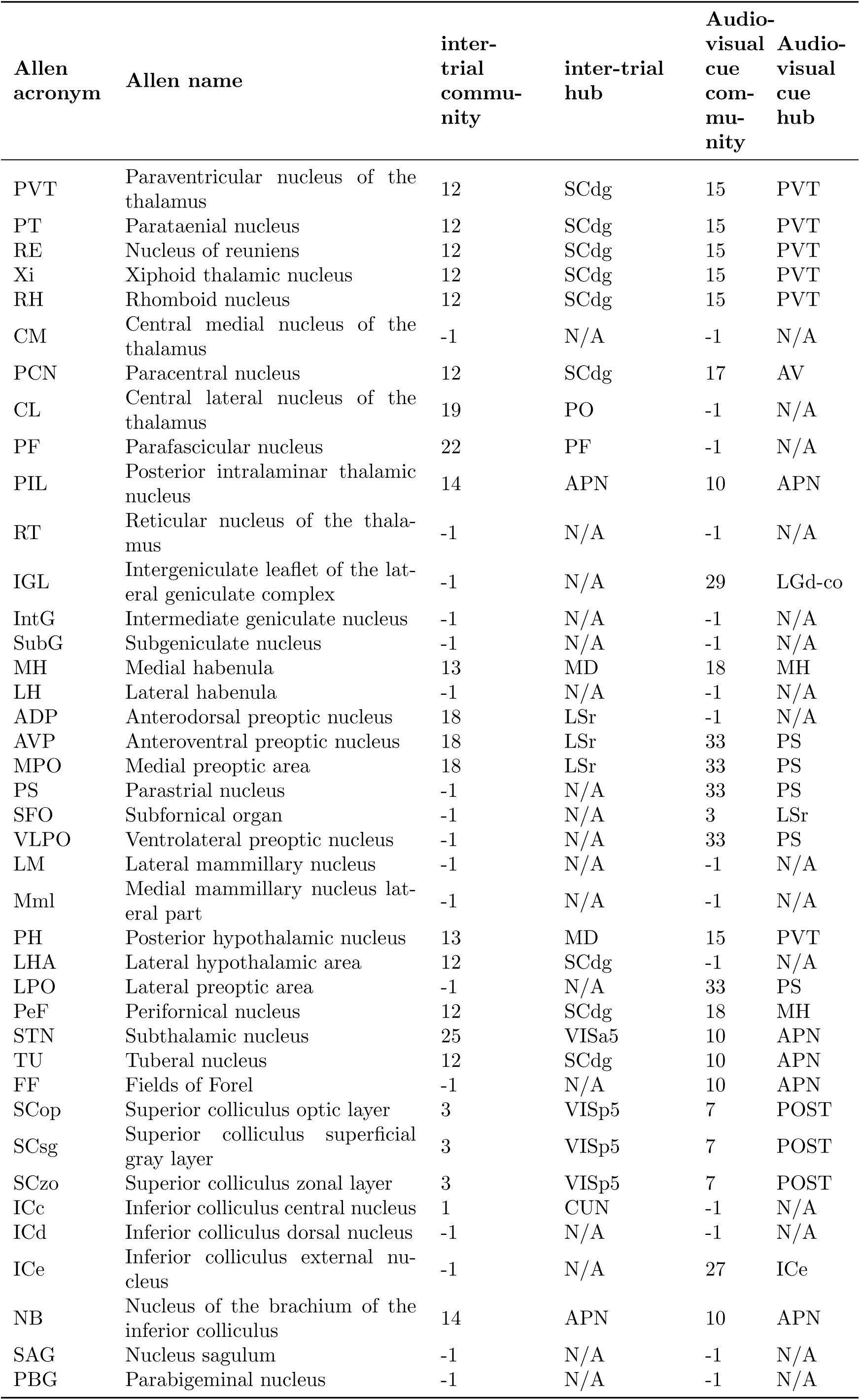
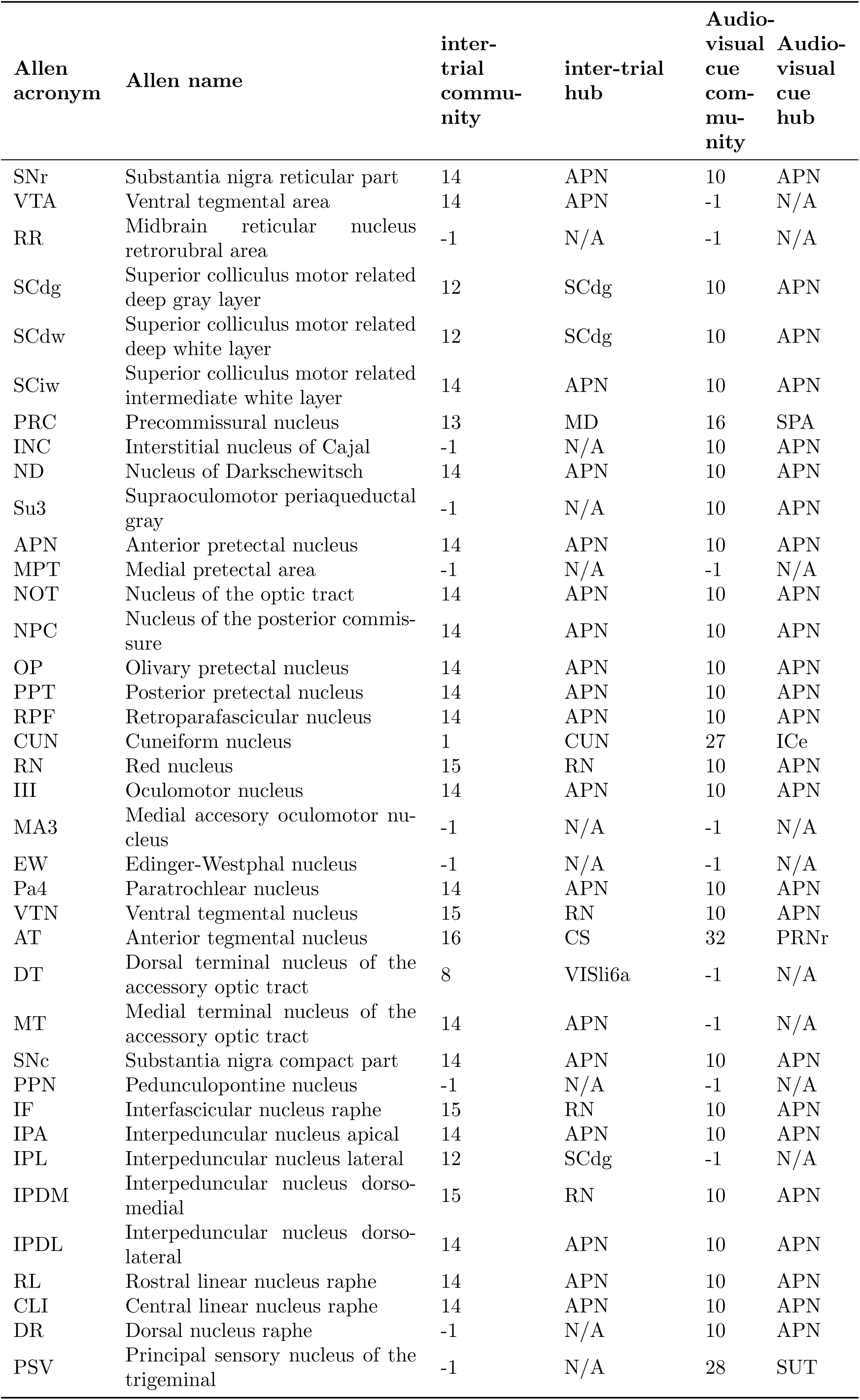
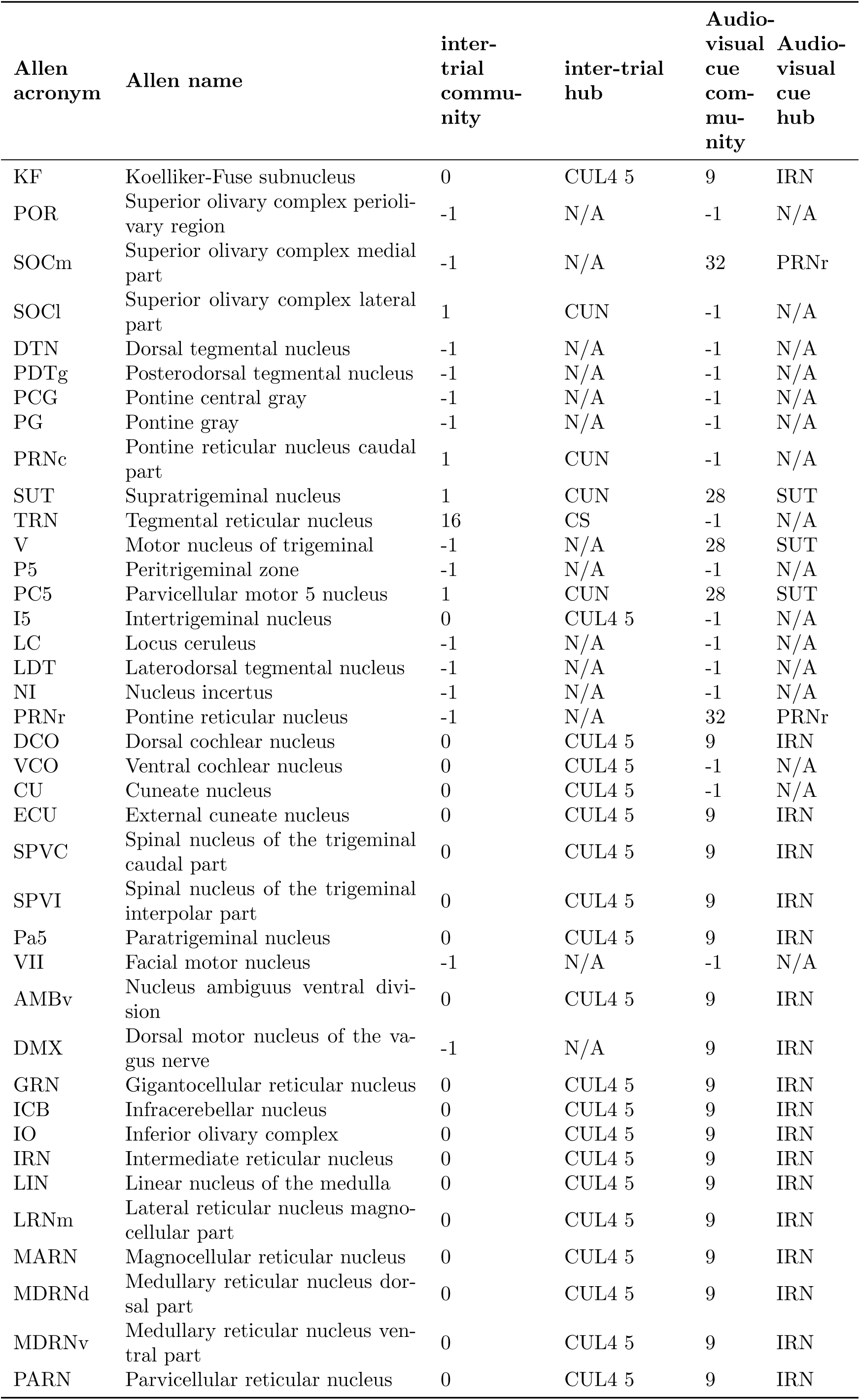
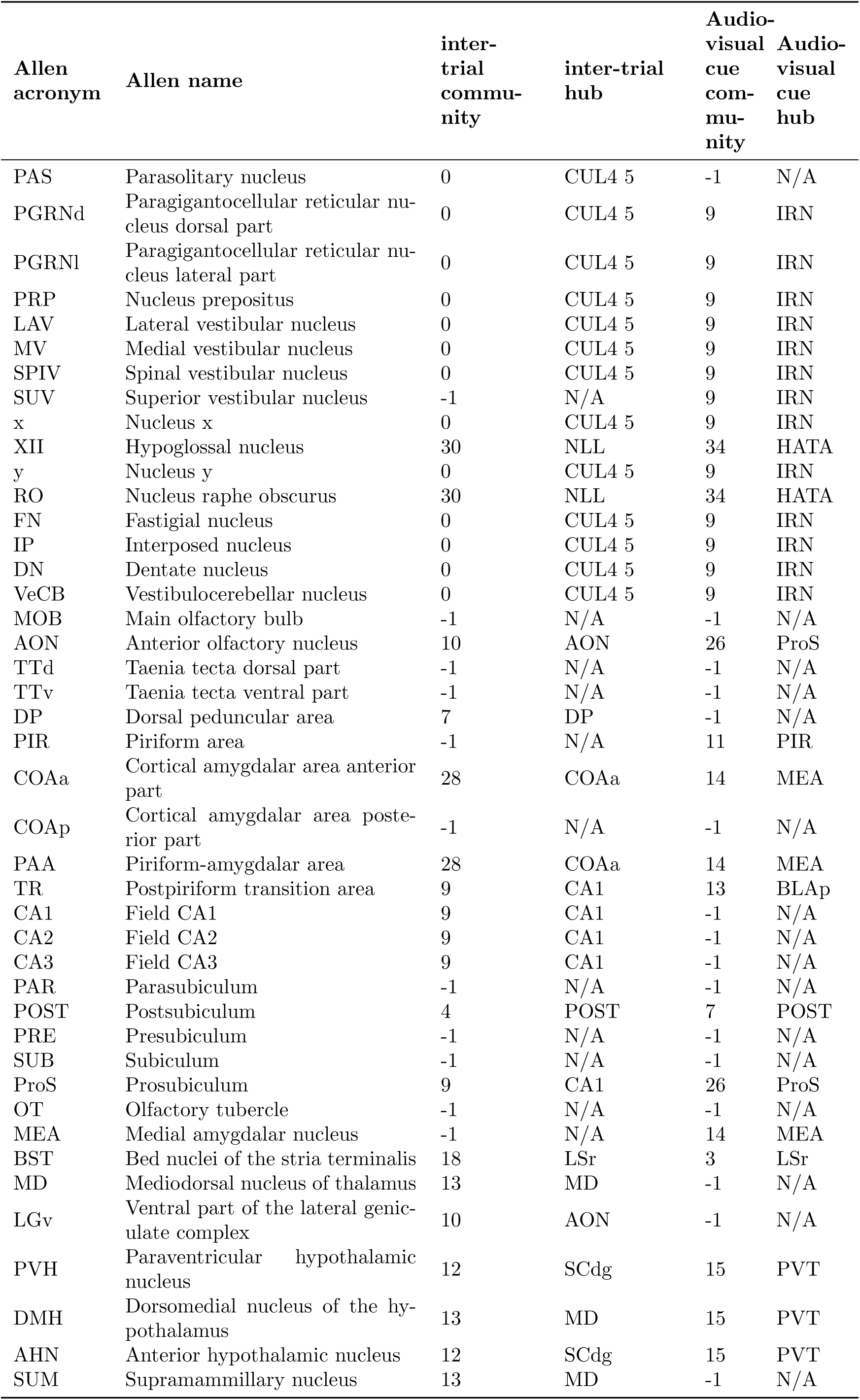
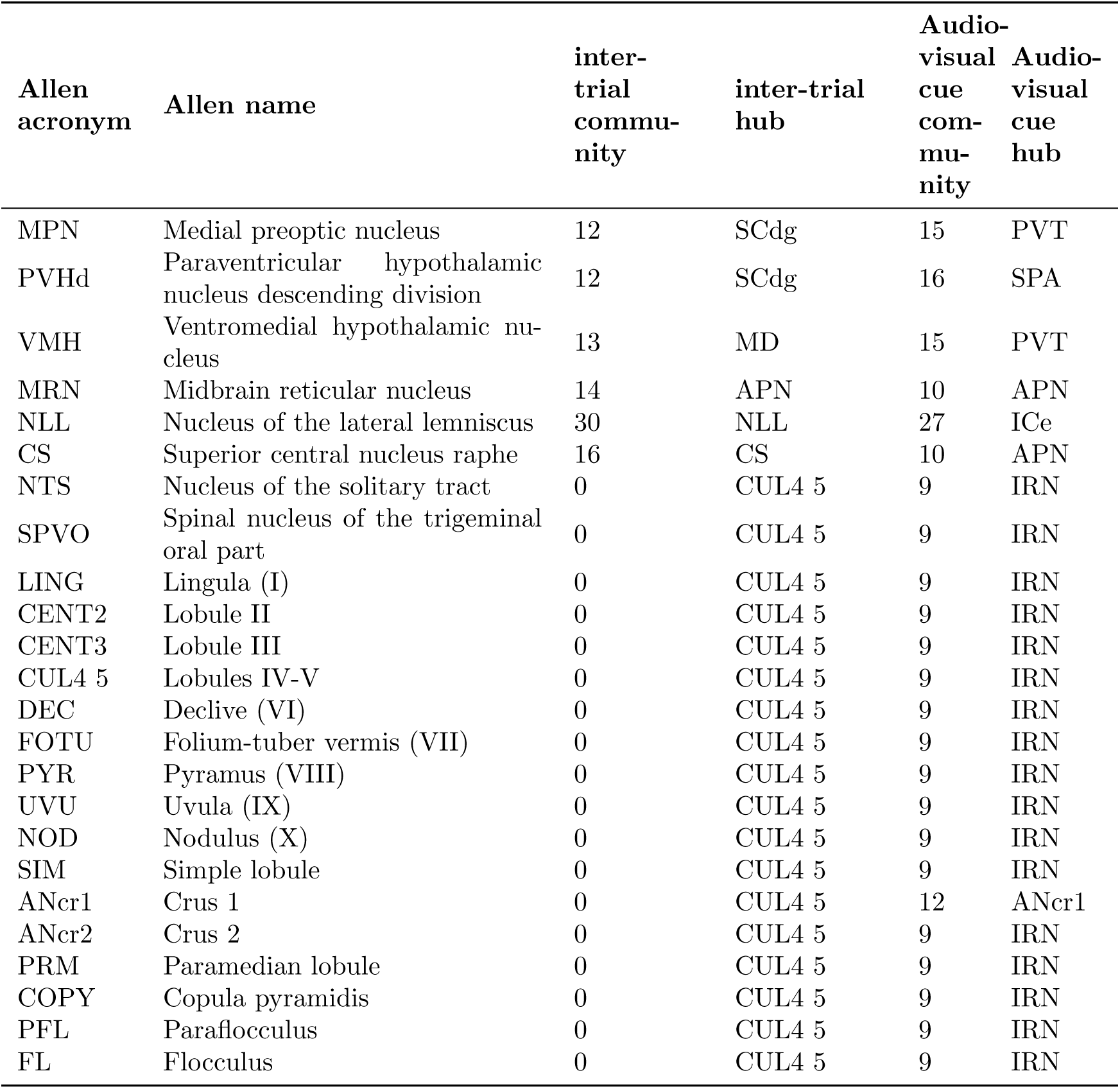
Table for community hub.

## References

[1] Nicholas A Steinmetz, Cagatay Aydin, Anna Lebedeva, Michael Okun, Marius Pachitariu, Marius Bauza, Maxime Beau, Jai Bhagat, Claudia Böhm, Martijn Broux, et al. Neuropixels 2.0: A miniaturized high-density probe for stable, long-term brain recordings. Science, 372(6539):eabf4588, 2021.

[2] Charles Findling, Felix Hubert, International Brain Laboratory, Luigi Acerbi, Brandon Benson, Julius Benson, Daniel Birman, Niccolòo Bonacchi, E Kelly Buchanan, Sebastian Bruijns, et al. Brain-wide representations of prior information in mouse decision-making. Nature, 645(8079):192–200, 2025.

[3] Jerry A Fodor. Precis of the modularity of mind. Behavioral and brain sciences, 8(1):1–5, 1985.

[4] David E Rumelhart, James L McClelland, PDP Research Group, et al. Parallel distributed processing, volume 1: Explorations in the microstructure of cognition: Foundations. The MIT press, 1986.

[5] Francis Jeffry Pelletier. The principle of semantic compositionality. Topoi, 13(1):11–24, 1994.

[6] Dieuwke Hupkes, Verna Dankers, Mathijs Mul, and Elia Bruni. Compositionality decomposed: How do neural networks generalise? Journal of Artificial Intelligence Research, 67:757–795, 2020.

[7] Bayanne Olabi, Ian Ellison-Wright, Andrew M McIntosh, Stephen J Wood, Ed Bullmore, and Stephen M Lawrie. Are there progressive brain changes in schizophrenia? a meta-analysis of structural magnetic resonance imaging studies. Biological psychiatry, 70(1):88–96, 2011.

[8] Jinseop S Kim and Marcus Kaiser. From caenorhabditis elegans to the human connectome: a specific modular organization increases metabolic, functional and developmental efficiency. Philosophical Transactions of the Royal Society B: Biological Sciences, 369(1653):20130529, 2014.

[9] David Meunier, Renaud Lambiotte, Alex Fornito, Karen Ersche, and Edward T Bullmore. Hierarchical modularity in human brain functional networks. Frontiers in neuroinformatics, 3:571, 2009.

[10] Olaf Sporns and Richard F Betzel. Modular brain networks. Annual review of psychology, 67(1):613–640, 2016.

[11] Christopher J Honey, Rolf Kötter, Michael Breakspear, and Olaf Sporns. Network structure of cerebral cortex shapes functional connectivity on multiple time scales. Proceedings of the National Academy of Sciences, 104(24):10240–10245, 2007.

[12] Brian Zingg, Houri Hintiryan, Lin Gou, Monica Y Song, Maxwell Bay, Michael S Bienkowski, Nicholas N Foster, Seita Yamashita, Ian Bowman, Arthur W Toga, et al. Neural networks of the mouse neocortex. Cell, 156(5):1096–1111, 2014.

[13] Julie A Harris, Stefan Mihalas, Karla E Hirokawa, Jennifer D Whitesell, Hannah Choi, Amy Bernard, Phillip Bohn, Shiella Caldejon, Linzy Casal, Andrew Cho, et al. Hierarchical organization of cortical and thalamic connectivity. Nature, 575(7781):195–202, 2019.

[14] Xiaoyin Chen, Stephan Fischer, Mara CP Rue, Aixin Zhang, Didhiti Mukherjee, Patrick O Kanold, Jesse Gillis, and Anthony M Zador. Whole-cortex in situ sequencing reveals input-dependent area identity. Nature, pages 1–10, 2024.

[15] James J Jun, Nicholas A Steinmetz, Joshua H Siegle, Daniel J Denman, Marius Bauza, Brian Barbarits, Albert K Lee, Costas A Anastassiou, Alexandru Andrei, Çăgatay Aydın, et al. Fully integrated silicon probes for high-density recording of neural activity. Nature, 551(7679):232–236, 2017.

[16] Angelique C Paulk, Yoav Kfir, Arjun R Khanna, Martina L Mustroph, Eric M Trautmann, Dan J Soper, Sergey D Stavisky, Marleen Welkenhuysen, Barundeb Dutta, Krishna V Shenoy, et al. Large-scale neural recordings with single neuron resolution using neuropixels probes in human cortex. Nature Neuroscience, 25(2):252–263, 2022.

[17] Larry F Abbott, Dora E Angelaki, Matteo Carandini, Anne K Churchland, Yang Dan, Peter Dayan, Sophie Deneve, Ila Fiete, Surya Ganguli, Kenneth D Harris, et al. An international laboratory for systems and computational neuroscience. Neuron, 96(6):1213–1218, 2017.

[18] International Brain Laboratory, Valeria Aguillon-Rodriguez, Dora Angelaki, Hannah Bayer, Nic-colòo Bonacchi, Matteo Carandini, Fanny Cazettes, Gaelle Chapuis, Anne K Churchland, Yang Dan, et al. Standardized and reproducible measurement of decision-making in mice. Elife, 10:e63711, 2021.

[19] György Buzśaki, Costas A Anastassiou, and Christof Koch. The origin of extracellular fields and currents—eeg, ecog, lfp and spikes. Nature reviews neuroscience, 13(6):407–420, 2012.

[20] Pieter R Roelfsema, Andreas K Engel, Peter König, and Wolf Singer. Visuomotor integration is associated with zero time-lag synchronization among cortical areas. Nature, 385(6612):157–161, 1997.

[21] Marius Schneider, Ana Clara Broggini, Benjamin Dann, Athanasia Tzanou, Cem Uran, Swathi Sheshadri, Hansjörg Scherberger, and Martin Vinck. A mechanism for inter-areal coherence through communication based on connectivity and oscillatory power. Neuron, 109(24):4050–4067, 2021.

[22] Theodoros P Zanos, Patrick J Mineault, Konstantinos T Nasiotis, Daniel Guitton, and Christopher C Pack. A sensorimotor role for traveling waves in primate visual cortex. Neuron, 85(3):615–627, 2015.

[23] J Jesús Herńandez-Péerez, Keiland W Cooper, and Ehren L Newman. Medial entorhinal cortex activates in a traveling wave in the rat. Elife, 9:e52289, 2020.

[24] Wolfgang Klimesch. Alpha-band oscillations, attention, and controlled access to stored information. Trends in cognitive sciences, 16(12):606–617, 2012.

[25] Charline Peylo, Yannik Hilla, and Paul Sauseng. Cause or consequence? alpha oscillations in visuospatial attention. Trends in Neurosciences, 44(9):705–713, 2021.

[26] Rosanne M Van Diepen, John J Foxe, and Ali Mazaheri. The functional role of alpha-band activity in attentional processing: the current zeitgeist and future outlook. Current opinion in psychology, 29:229–238, 2019.

[27] Supratim Ray and John HR Maunsell. Do gamma oscillations play a role in cerebral cortex? Trends in cognitive sciences, 19(2):78–85, 2015.

[28] Ole Jensen, Jochen Kaiser, and Jean-Philippe Lachaux. Human gamma-frequency oscillations associated with attention and memory. Trends in neurosciences, 30(7):317–324, 2007.

[29] Ole Jensen, Bart Gips, Til Ole Bergmann, and Mathilde Bonnefond. Temporal coding organized by coupled alpha and gamma oscillations prioritize visual processing. Trends in neurosciences, 37(7):357–369, 2014.

[30] Antonio Fernandez-Ruiz, Anton Sirota, Véıtor Lopes-dos Santos, and David Dupret. Over and above frequency: gamma oscillations as units of neural circuit operations. Neuron, 111(7):936–953, 2023.

[31] György Buzśaki. Hippocampal sharp wave-ripple: A cognitive biomarker for episodic memory and planning. Hippocampus, 25(10):1073–1188, 2015.

[32] Bernhard P Staresina and Maria Wimber. A neural chronometry of memory recall. Trends in cognitive sciences, 23(12):1071–1085, 2019.

[33] Yuta Senzai and György Buzśaki. Physiological properties and behavioral correlates of hippocampal granule cells and mossy cells. Neuron, 93(3):691–704, 2017.

[34] Diego Mendoza-Halliday, Alex James Major, Noah Lee, Maxwell J Lichtenfeld, Brock Carlson, Blake Mitchell, Patrick D Meng, Yihan Xiong, Jacob A Westerberg, Xiaoxuan Jia, et al. A ubiquitous spectrolaminar motif of local field potential power across the primate cortex. Nature Neuroscience, 27(3):547–560, 2024.

[35] Tianxiao He, Malhar Patel, Chenyi Li, Anna Maslarova, Mihéaly Vöröslakos, Nalini Ramanathan, Wei-Lun Hung, Gyorgy Buzsaki, and Erdem Varol. Self supervised learning for in vivo localization of microelectrode arrays using raw local field potential. In The Thirty-ninth Annual Conference on Neural Information Processing Systems.

[36] Dora Angelaki, Brandon Benson, Julius Benson, Daniel Birman, Niccolòo Bonacchi, Kćenia Bougrova, Sebastian A Bruijns, Matteo Carandini, Joana A Catarino, et al. A brain-wide map of neural activity during complex behaviour. Nature, 645(8079):177–191, 2025.

[37] Gemechu Bekele Tolossa, Aidan M Schneider, Eva Dyer, and Keith B Hengen. Neurons through-out the brain embed robust signatures of their anatomical location into spike trains. eLife, 13:RP101506, 2025.

[38] Alain Destexhe, Diego Contreras, and Mircea Steriade. Spatiotemporal analysis of local field potentials and unit discharges in cat cerebral cortex during natural wake and sleep states. Journal of Neuroscience, 19(11):4595–4608, 1999.

[39] Susan M Sunkin, Lydia Ng, Chris Lau, Tim Dolbeare, Terri L Gilbert, Carol L Thompson, Michael Hawrylycz, and Chinh Dang. Allen brain atlas: an integrated spatio-temporal portal for exploring the central nervous system. Nucleic acids research, 41(D1):D996–D1008, 2012.

[40] Ilija Radosavovic, Raj Prateek Kosaraju, Ross Girshick, Kaiming He, and Piotr Dolĺar. Designing network design spaces. In Proceedings of the IEEE/CVF conference on computer vision and pattern recognition, pages 10428–10436, 2020.

[41] Ashish Vaswani, Noam Shazeer, Niki Parmar, Jakob Uszkoreit, Llion Jones, Aidan N Gomez, Ƚukasz Kaiser, and Illia Polosukhin. Attention is all you need. Advances in neural information processing systems, 30, 2017.

[42] Alexey Dosovitskiy, Lucas Beyer, Alexander Kolesnikov, Dirk Weissenborn, Xiaohua Zhai, Thomas Unterthiner, Mostafa Dehghani, Matthias Minderer, Georg Heigold, Sylvain Gelly, et al. An image is worth 16x16 words: Transformers for image recognition at scale. arXiv preprint arXiv:2010.11929, 2020.

[43] Patel Dhruv and Subham Naskar. Image classification using convolutional neural network (cnn) and recurrent neural network (rnn): A review. Machine learning and information processing: proceedings of ICMLIP 2019, pages 367–381, 2020.

[44] Mark EJ Newman. Modularity and community structure in networks. Proceedings of the national academy of sciences, 103(23):8577–8582, 2006.

[45] Aaron Clauset, Mark EJ Newman, and Cristopher Moore. Finding community structure in very large networks. *Physical Review E—Statistical*, Nonlinear, and Soft Matter Physics, 70(6):066111, 2004.

[46] David Meunier, Renaud Lambiotte, and Edward T Bullmore. Modular and hierarchically modular organization of brain networks. Frontiers in neuroscience, 4:200, 2010.

[47] International Brain Laboratory, Dora Angelaki, Brandon Benson, Julius Benson, Daniel Birman, Niccolòo Bonacchi, Kćenia Bougrova, Sebastian A Bruijns, Matteo Carandini, Joana A Catarino, et al. A brain-wide map of neural activity during complex behaviour. Nature, 645(8079):177, 2025.

[48] Guido Meijer Huntenburg, Liam Paninski, Michael Schartner, Karel Svoboda, Miles Wells, Matthew R Whiteway, Olivier Winter, et al. Video hardware and software for the international brain laboratory.

[49] Alex Graves, Marcus Liwicki, Santiago Ferńandez, Roman Bertolami, Horst Bunke, and Jürgen Schmidhuber. A novel connectionist system for unconstrained handwriting recognition. IEEE transactions on pattern analysis and machine intelligence, 31(5):855–868, 2008.

[50] Aric Hagberg, Pieter J Swart, and Daniel A Schult. Exploring network structure, dynamics, and function using networkx. Technical report, Los Alamos National Laboratory (LANL), Los Alamos, NM (United States), 2008.

